# Musashi-2 causes cardiac hypertrophy and heart failure by inducing mitochondrial dysfunction through destabilizing *Cluh* and *Smyd1* mRNA

**DOI:** 10.1101/2023.03.01.530580

**Authors:** Sandhya Singh, Aakash Gaur, Renu Kumari, Shakti Prakash, Sunaina Kumari, Ayushi Devendrasingh Chaudhary, Rakesh Kumar Sharma, Pankaj Prasun, Priyanka Pant, Thomas Thum, Kumaravelu Jagavelu, Pragya Bharati, Kashif Hanif, Pragya Chitkara, Shailesh Kumar, Kalyan Mitra, Shashi Kumar Gupta

**Author notes:** Correspondence Address: Shashi Kumar Gupta, PhD, Pharmacology Division CSIR-CDRI, Sector 10, Jankipuram Extension, Lucknow, India 226031.

## Abstract

Regulation of RNA stability and translation by RNA-binding proteins (RBPs) is a crucial process altering gene expression. Musashi family of RBPs comprising *Msi1* and *Msi2* are known to control RNA stability and translation. However, despite the presence of MSI2 in the heart, its function remains entirely unknown. Here, we aim to explore the cardiac functions of MSI2. We confirmed the presence of MSI2 in the adult mouse, rat heart, and neonatal rat cardiomyocytes. Furthermore, *Msi2* was significantly enriched in the heart’s cardiomyocyte fraction. Next, using RNA-seq data and isoform-specific PCR primers, we identified, *Msi2* isoforms 1, 4, and 5 and two novel putative isoforms labeled as *Msi2* isoforms 6 and 7 to be expressed in the heart. Overexpression of *Msi2* isoforms led to cardiac hypertrophy in cultured cardiomyocytes. Additionally, *Msi2* was also found to be significantly increased in a pressure-overload model of cardiac hypertrophy. To validate the hypertrophic effects, we selected isoforms 4 and 7 due to their unique alternative splicing patterns. AAV9-mediated overexpression of *Msi2* isoforms 4 and 7 in murine hearts led to cardiac hypertrophy, dilation, heart failure, and eventually early death, confirming a pathological function for *Msi2*. Using global proteomics, gene ontology, transmission electron microscopy, and transmembrane potential measurement assays increased MSI2 was found to cause mitochondrial dysfunction in the heart. Mechanistically, we identified *Cluh* and *Smyd1* as direct downstream targets of *Msi2*. Overexpression of *Cluh* or *Smyd1* inhibited *Msi2*-induced hypertrophy and mitochondrial dysfunction in cardiomyocytes. Collectively, we show that *Msi2* induces hypertrophy, mitochondrial dysfunction, and heart failure.

## Introduction

Despite the availability of heart failure therapies, point-of-care, and our increasing understanding of disease mechanisms, cardiovascular diseases are still globally the leading cause of mortality [23]. Heart failure is the eventual end-point of all cardiovascular diseases leading to more than eighteen million deaths in 2019 around the globe [23]. Therefore, there is an unmet need to understand the pathophysiology of cardiovascular diseases better and identify novel druggable targets which would reduce the mortality burden. One key feature of the failing heart is energy deprivation due to abnormalities in mitochondrial structure, dynamics, and function [25]. Mitochondria is a semi-autonomous body with genome-encoding proteins, rRNAs, and tRNAs [28]. Besides the mitochondrial genome, nucleus-encoded regulatory factors regulate mitochondrial biogenesis, quality, and function [28]. These nucleus-encoded regulatory factors regulate the transcription, processing, and translation of mitochondrial genes and several other nuclear genes involved in the mitochondrial function directly or indirectly [28]. Alteration in these broad regulators affects the mitochondrial structure and function by altering the downstream gene expression.

Transcriptional or post-transcriptional processes are involved in gene expression regulation [6]. Post-transcriptional regulation of gene expression is mainly through RNA [6]. RNA-binding proteins (RBPs) are one of the primary mediators of post-transcriptional gene expression regulation by their RNA-binding function [6]. Humans and mice have over a thousand RBPs that bind to RNAs by canonical or non-canonical RNA-binding domains [13]. However, detailed studies describing the cardiovascular role of RBPs are scanty. RBPs regulate all critical steps during the life cycle of RNA, soon after its transcription until its decay [6]. RBPs regulate 5’ capping, splicing, 3’ polyadenylation, export, translation, decay, and stability of RNA molecules [6]. RNA stability and decay are crucial for cellular homeostasis, and its misregulation may lead to cardiovascular diseases [32]. Several RBPs promote the stability and decay of RNAs like PAIP2, YTHDF1-3, HuR, AUF1, TTP, KSRP, and MSI1-2 [31, 32] [15].

Musashi (MSI) family of RBPs is characterized by the presence of tandem RNA recognition motifs (RRM). Musashi was first identified as a regulator of adult sensory organ development in Drosophila [19]. In mammals, two Musashi isoforms, *MSI1* and *MSI2,* are found and have similar RNA binding specificities [15]. The function of MSI varies from translation inhibition or initiation to polyadenylation, alternative splicing, and mRNA stabilization or decay [15]. Musashi proteins regulate the self-renewal potential of hematopoietic stem cells and crypt base columnar stem cells in the intestine [15]. Furthermore, they also promote hematopoietic malignancies and colorectal carcinomas [15]. Additionally, *Msi2* regulates the mitochondrial distribution of microRNA miR-301a-3p in endothelial cells [11].

Recently, two contradictory reports have been published regarding MSI2 function in the skeletal muscle. Increased MSI2 levels were found to be responsible for muscle wasting and atrophy in myotonic dystrophy type 1 [24]. On the contrary, Wang et al. show upregulation of MSI2 promote skeletal muscle differentiation, and MSI2 KO mice show defective muscle regeneration [34]. Mechanistically, both studies report repression of miR-7a processing by MSI2 as the downstream mechanism warranting further investigations. In mice, *Msi1* was primarily expressed in the brain, small intestine, and ovary while absent in the heart [26]. On the contrary, *Msi2* showed more ubiquitous expression and was expressed in the heart [27]. Despite the expression of MSI2 in the heart, its role remains largely unknown. Here, we aim to study the cardiac function of MSI2.

Here, we report a pro-hypertrophic role for MSI2 in cardiomyocytes leading to heart failure and death in mice. AAV9 (Adeno-associated virus serotype 9) mediated overexpression of *Msi2* promoted degradation of *Cluh* and *Smyd1* and thus led to mitochondrial dysfunction. Overexpression of *Cluh* or *Smyd1* inhibits the pro-hypertrophic and mitochondrial dysfunction induced by *Msi2*.

## Methods

### Cell culture and lentivirus

Human cardiomyocyte cell line AC16 was cultured in DMEM/F-12 medium (Thermo Scientific, USA) together with 12.5% fetal bovine serum (Thermo Scientific, USA), penicillin and streptomycin (Thermo Scientific, USA), and 15 mM HEPES at 37°C in the presence of 5% CO_2_. Permanent AC16 cell lines were made by lentiviral-mediated transduction and puromycin selection. Lentiviruses were produced by transfection of HEK293T cells with lenti-plasmid, psPAX2, and pMD2.G using PEI Max 40000 (PolySciences, USA). The supernatant containing lentiviral particles was harvested and mixed with forty percent PEG 8000 (Sigma Aldrich, USA) and incubated for four hours at 4°C with 60 rpm rotation. Precipitated lentiviral particles were resuspended in 1X PBS. HEK293T cells were cultured in DMEM high glucose medium with 10% fetal bovine serum (Thermo Scientific, USA), penicillin and streptomycin (Thermo Scientific, USA).

### Neonatal rat cardiomyocytes

One to three-day-old SD rat neonates were used to isolate primary cardiomyocytes. Hearts were explanted and digested with Collagenase Type II at 1mg/ml (Thermo Scientific, USA) at 37°C. Digested cells were pellet down and resuspended in DMEM with 20% fetal bovine serum and penicillin/streptomycin. Resuspended cells were then plated in 10 cm dishes and incubated inside a cell culture incubator (Eppendorf, Germany) at 37°C with 5% CO2 for ninety minutes. Non-adherent cells were collected, and cardiomyocytes were counted. Counted cardiomyocytes were seeded in gelatin-coated cell culture plates with DMEM and 20% fetal bovine serum and kept inside the cell culture incubator. Forty-eight hours later, cells were washed with PBS, and a fresh DMEM medium with 1% fetal bovine serum was added. Cardiomyocytes were transduced with AAV6 at MOI of 0.1-1*10^5^ and cultured for four to five days. To measure mitochondrial transmembrane potential, we treated transduced cardiomyocytes with 100nM TMRE-Red (Thermo Scientific, USA). After thirty minutes of treatment, imaging was done using Leica DMI 6000 B (Leica Microsystems, USA) at 20X objective. Red fluorescence intensity was calculated with Image J. For isoproterenol treatment, cardiomyocytes were treated with 20μM isoproterenol hydrochloride (Sigma-Aldrich) for seventy-two hours. All the experiments were performed three independent times (biological replicate) with three replicates per group (technical replicate). For actinomycin D treatment, cardiomyocytes were treated at a dose of 5μg/ml after sixty hours of AAV6 transduction. Cells were collected at zero, four, and six hours post-treatment.

### Adeno-associated virus (AAV)

All the *Msi2* isoforms cDNA were cloned in AAV-CMV plasmid. Adeno-associated viruses were produced according to a previously published protocol [17]. In brief, HEK293T cells were transfected with AAV plasmid and helper plasmid using PEI Max 40000 (PolySciences, USA). Twenty-four hours later, the transfection medium was removed, and fresh DMEM (Thermo Scientific, USA) with 1% fetal bovine serum (Thermo Scientific, USA), penicillin/streptomycin (Thermo Scientific, USA), 10mM HEPES (Sigma Aldrich, USA), 0.075% NaHCO3 and 1X Glutamax (Thermo Scientific, USA). Cells were incubated for forty-eight hours, a culture medium containing AAV particles was harvested, and a fresh production medium was added. Once again, forty-eight hours later, culture medium with viral particles was harvested. HEK293T cells containing AAV viral particles were collected and lysed with citrate buffer. Medium with viral particles and cell lysate was mixed with 40% PEG 8000 and incubated overnight at 4°C. The next day precipitated viral particles were harvested by centrifugation and resuspended in PBS. AAV viral particles were then cleaned by chloroform and used for *in vitro* applications. For *in vivo* use, chloroform-cleaned viruses were loaded on Optipprep (Sigma Aldrich, USA) density gradient and ultracentrifuged (Beckman Coulter, USA). AAV viral particles were then collected from 40% gradient by puncturing the ultracentrifuge tube. The collected viral particles were then cleaned and concentrated using Amicon 100K (100kDa) cut-off columns (Sigma Aldrich, USA). AAV-GFP was used as a control.

### RNA and PCR

Total RNA from cell culture or heart tissue was isolated using RNA iso (Takara Bio, Japan) per the manufacturer’s protocol. A total of 1-2 μg RNA was reverse transcribed with random hexamers using PrimeScript 1^st^ strand cDNA synthesis kit (Takara Bio, Japan) per the manufacturer’s protocol on Verti 96 well thermocycler (Thermo Scientific, USA). Real-time quantitative PCR of mRNA was performed using TB Green Premix Extaq (Takara Bio) as per the manufacturer’s protocol on Quant Studio 12K Flex (Thermo Scientific, USA) using specific primers listed in **Supplementary Table 1**. For isoform detection, cDNA was amplified using Emerald Amp GT PCR mix (Takara Bio, Japan) on Verti 96 well thermocycler (Thermo Scientific, USA) by primer pair - forward primer 5’ CTACCCCAACTTTGTGGCAAC 3’ reverse primer 5’ GCCTGGACATCCAGGTATGC 3’. All the primers were synthesized from Sigma-Aldrich. Barcode Biosciences, Bangalore, India, did DNA sequencing.

### Western Blotting

Cell pellets were lysed in 1X RIPA lysis buffer and sonicated. A small piece was crushed in 1X RIPA buffer for heart tissue with liquid N2. Protein quantification of isolated lysates was done using a BCA kit (Thermo Scientific, USA). Twenty-five to forty micrograms of lysate were loaded on 12% SDS-PAGE gel to resolve the proteins. Resolved proteins were transferred to the PVDF membrane (BioRad, USA) using a Mini PROTEAN Tetra cell (Biorad, USA). The membrane was then incubated with specific antibodies for detection of MSI2 (#PA5-31024 Thermo Scientific, USA), GAPDH (#MA515738 Thermo Scientific, USA), and ACTB (#A00730-100 Genescript, USA). Secondary antibodies linked to HRP (Thermo Scientific, USA) were used for detection. Precision Plus Protein Western C ladder (BioRad) and Precision protein Streptactin-HRP conjugate (BioRad) were used to determine protein size on the membrane.

### Luciferase reporter assay

3’UTR of *Cluh* and *Smyd1* was amplified by PCR from mouse heart cDNA and cloned downstream to luciferase. HEK293T cells were transfected with luciferase plasmid, *Msi2* 4 or 7 encoding plasmids, and beta-gal plasmid using PEI Max 40000 (PolySciences, USA). The next day transfection medium was exchanged with fresh medium, and cells were further grown for forty-eight hours. Cells were lysed, and luciferase activity was measured with a GloMax Navigator microplate luminometer (Promega, USA) using a luciferase assay kit (#E1500 Promega, USA) as per the manufacturer’s protocol. Per the manufacturer’s instruction, the beta-galactosidase enzyme activity was measured using a kit (#E2000, Promega, USA). Luciferase readings were divided with beta-galactosidase readings to normalize for transfection differences. Three independent experiments were performed with three replicates per group each time.

### Cell size measurements

Neonatal rat cardiomyocytes were stained with alpha-sarcomeric actinin (#MA1-22863 Thermo Scientific, USA) and DAPI. AC16 cells were stained with wheat germ agglutinin (#MP00831 Thermo Scientific, USA) and DAPI. All images were taken with 20X objective using Leica DMI 6000 B (Leica Microsystems, USA). Cell size was calculated with ImageJ software. All the experiments were repeated three times (biological replicate) with three wells for each group (technical replicate). More than a hundred cells were counted from each well.

Paraffin-embedded heart sections were deparaffinized and stained with wheat germ agglutinin (#MP00831 Thermo Scientific, USA) and DAPI. Images from different regions of the heart were taken with a 20X objective using Leica DMI 6000 B (Leica Microsystems, USA). Cell size was measured using ImageJ software.

### Fibrosis staining

Picrosirius red (Sigma Aldrich, USA) staining was done on deparaffinized heart sections. Images were taken with Leica DFC 320 (Leica Microsystems, USA). The percentage fibrosis stain was calculated using Adobe Photoshop.

### Immunofluorescence

AC16 cells were fixed with four percent paraformaldehyde and permeabilized with 0.1% Triton-X-100. Cells were washed with PBS and incubated with 5% BSA for blocking. Blocked cells were incubated overnight with primary antibody against MSI2 (#PA5-31024 Thermo Scientific, USA). The following day cells were stained with Alexa-Flour 594 labeled secondary antibody (Thermo Scientific, USA) and DAPI. Images were taken with 20X objective using Leica DMI 6000 B (Leica Microsystems, USA) to check the localization of MSI2.

### Immunohistochemistry

Paraffin-embedded heart sections were cut using a Leica microtome and deparaffinized. Deparaffinized sections were then heated at 80°C in citrate buffer. Sections were then quenched for endogenous peroxidase activity by H2O2. Next sections were permeabilized using 0.3% Triton-X-100 and blocked with 5% BSA. Blocked sections were then incubated with MSI2 antibody overnight at 4°C. The next day, sections were incubated with HRP-labeled secondary antibody at room temperature for two hours. Sections were then exposed to DAB substrate and counter-stained with hematoxylin for nuclear staining.

### Transmission Electron microscopy

TEM experiments were performed as described with minor modifications [2]. The mouse heart’s tissue pieces (1mm3) were fixed in 2.5% glutaraldehyde and 4% paraformaldehyde in 0.1M phosphate buffer, pH 7.4, and post-fixed in 1% osmium tetroxide. This was followed by dehydration in ascending series of ethanol, infiltration, and embedding in Spurr resin. Ultra-thin sections (60-70nm) were obtained using Leica EM UC7 ultra-microtome (Wetzlar, Germany), picked up on 200 mesh copper grids, and dual stained with uranyl acetate and lead citrate. Grids were observed under JEOL JEM 1400 TEM, and data were collected using a Gatan Orius SC 200B CCD camera at 100kV with GATAN digital micrograph software.

### Animal experiments

All the animal experiments were carried out on C57BL/6 mice provided by National Laboratory Animal Centre, CSIR-CDRI, Lucknow, India. C57BL/6 adult male mice were injected with 1.8*10^12^ AAV9 viral particles intravenously. Echocardiography was done to analyze cardiac function using Vevo 1100 (Fujifilm VisualSonics, USA) under 2-3% isoflurane given by inhalation. Echocardiography was done after fourteen days post viral injection for AAV9-*Msi2* 4 and twenty-one days for AAV9-*Msi2* 7 injected animals, respectively. Mice were euthanized by cervical dislocation after anesthesia with isoflurane (3%), and the heart was explanted for further molecular and histological assays. The tibia was taken out, and the length was measured. For the TAC surgery, adult SD rats were used. The transverse aorta was partially occluded using an 18G needle. Post-surgery rats were initially recovered under a warming lamp and observed through the course of experiments. The local IAEC (Institutional Animal Ethics Committee) committee at CSIR-Central Drug Research Institute approved all the animal experiments (IAEC/2020/38) following the guidelines of the Committee for the Purpose of Control and Supervision of Experiments on Animals (CPCSEA), New Delhi, Government of India. All the animal procedures performed conform to the guidelines from Directive 2010/63/EU of the European Parliament on the protection of animals used for scientific purposes.

### Global Proteomics

AAV9-GFP and AAV9-*Msi2* 4 injected animals were sacrificed fourteen days post-injection, and hearts were explanted for proteomics. Vproteomics, Valerian Chem Private Limited, India, performed the proteomics per the following protocol. Trypsin digestion was done at 37°C for 16 hours with fifty micrograms of heart tissue. Digested lysates were cleaned using a C18 silica cartridge and dried. The dried pellet was resuspended in a buffer with 2% acetonitrile and 0.1% formic acid. Mass spectrometric analysis of the peptide mixtures was performed on Ultimate 3000 RSLC nano system coupled with a Tribrid Orbitrap Eclipse (Thermo Fisher Scientific, USA). 1μg of the sample was loaded on Acclaim PepMap 75 μm x 2 cm C18 guard column (3μm particle size). Peptides were eluted with a 0–40 % gradient of a buffer consisting of 80 % acetonitrile and 0.1 % formic acid and separated on a 50 cm, three μm Easy-spray C18 column (Thermo Fisher Scientific, USA) at a flow rate of 300 nl / min and injected for MS analysis. LC gradients were run for 110 min. MS1 spectra were acquired in the Orbitrap (R= 240k; AGQ target = 400,000; Max IT = 50 ms; RF Lens = 30%; mass range = 375-1500 m/z; centroid data). Dynamic exclusion was employed for 10 sec, excluding all charge states for a given precursor. MS 2 spectra were collected in the linear ion trap (rate = Rapid; AGQ target = 30000; MaxIT = 20 ms; NCEHCD = 30%). Generated raw data were analyzed with Proteome Discoverer (v2.2) against the UniProt *Mus musculus* proteome (UP000000589) database. For the Sequest search, the precursor and fragment mass tolerances were set at ten ppm and 1.0 Da, respectively. The protease used to generate peptides, i.e., enzyme specificity, was set for trypsin/P (cleavage at the C terminus of “K/R: unless followed by “P”) and with GluC on Glutamic acid along with maximum missed cleavages value of two. Carbamidomethyl on cysteine as fixed modification and oxidation of methionine and N-terminal acetylation were considered variable modifications for database search. Both peptide spectrum match and protein false discovery rate were set to 0.01 FDR. Raw abundance values were used for the statistical analysis. For each test, raw abundance values were filtered based on valid values (They should be present in at least 70% of samples within the group). Missing values were imputed based on standard deviation. Abundance values were log2 standardized, Median Based Quantile Normalisation followed by Z-score scaling matrix used to plot Heatmap, Boxplot. For comparison of the two groups, student t-test was used.

### MSI2 pull-down

HEK293T cells were seeded in 150 mm cell culture dishes and transfected with luciferase plasmids having 3’UTR of *Cluh* and *Smyd1* (used for luciferase assay) together with MSI2 isoform four overexpressing plasmid. Cells were cross-linked with 254nm UV exposure for 10min after sixty hours of transfection. The cells were harvested, washed with PBS, and stored at −80°C until use. MSI2 pull-down was performed using Dynabeads antibody coupling kit (Invitrogen 14311D) per the manufacturer’s instruction. Dyna beads M-270 were washed per kit instructions and then incubated with 10μg of MSI2 and IgG antibody separately at 37°C with continuous shaking overnight. Cells were thawed and lysed with 1X NP-40 lysis buffer (1mL) for 15 min, and lysates were collected by centrifugation at 13000Xg for 15min at 4°C. 80μl of input volume was taken out for RNA isolation and western blot, and the remaining volume proceeded for pull-down. Lysates were incubated with antibody-coupled Dyna beads at 4°C for 1hr with rotation. The beads were then washed in RIP buffer, and 30% of the fraction was taken for protein and 70% for RNA, which were further proceeded for RNA isolation with miRNeasy kit.

### RNA sequencing analysis

Publicly available transcriptomic data for nine sham mouse samples were retrieved from the NCBI GEO database (Gene Expression Omnibus) under the accession number GSE180794, where three replicates of each cell type were obtained from cardiomyocytes, fibroblasts, and endothelial cardiac cell types to investigate the expression of the *Msi2* gene in these cell fractions of the heart. *Msi2*’s different transcript expression and exon coverage were calculated from the cardiomyocyte fraction. The raw sequencing reads downloaded were preprocessed for adaptor trimming and quality control using the FASTP program (v0.21.0) [4]. The HISAT2 tool was used to index the reference genome and annotation file for *Mus musculus,* which was downloaded from the NCBI genome database (assembly GRCm39) [16]. The high-quality FASTQ files of each cell type were aligned to the indexed reference genome using the HISAT2 RNA-Seq aligner. The alignment was then assembled into potential transcripts using StringTie (v2.2.0) [21]. The per-base transcript and exon coverage for each *Msi2* transcript were calculated. The StringTie assembled transcripts were merged with the reference annotation file to generate a non-redundant set of transcripts observed in any previously assembled samples. The transcript abundance was then calculated using the merged annotation file to generate re-estimated normalized read counts for each transcript as fragments per kilobase of transcript per million reads (FPKM) values.

### Statistics

All the data were analyzed with GraphPad Prism software. All the data are presented as mean±sem. T-test was done to calculate the significance between the two groups. One-way ANOVA with post hoc Dunnett or Tukey test was used to calculate significance between more than two groups wherever required. P value ≤0.05 was considered significant.

## Results

### 1. *In vitro* functional characterization of *Msi2*

To explore the functions of the Musashi family of RNA-binding proteins, we checked the expression of both family members, *Msi1* and *Msi2,* in the C57BL6 mouse hearts. Similar to published reports, *Msi1* was confirmed to be absent in the heart, while *Msi2* was well expressed (**Fig. 1A**). We also confirmed MSI2 expression at the protein level in primary neonatal SD rat cardiomyocytes, adult SD rat hearts, and adult C57BL6 mouse hearts (**Fig. 1B**). Furthermore, to study the cell type enrichment of *Msi2* in the heart, we used a publicly available RNA sequencing dataset (GSE180794) from mouse hearts fractionated into cardiomyocytes, fibroblast, and endothelial cells [8]. *Msi2* was significantly enriched in cardiomyocyte fraction compared to the healthy heart’s fibroblast and endothelial cell fraction (**Fig. 1C**). Next, five different validated isoforms of *Msi2* are listed in NCBI. However, to understand the cardiac function of *Msi2,* it is necessary to know the isoforms expressed in the heart. Hence, we took the cardiomyocyte fraction RNA sequencing data and checked for different *Msi2* transcripts. We found *Msi2* isoforms NM_054043.3 (isoform 1), NM_001363195.1 (isoform 4), NM_001373923.1 (isoform 5), XM_036157063.1 and XM_036157068.1 to be well expressed (**Supplementary Table 2**). Moreover, we also assessed exon coverage for these five isoforms and found the highest exon coverage for XM_036157068.1, followed by XM_036157063.1, isoform 5, isoform 4, and isoform 1 being the least (**Supplementary Table 3**). For ease of labeling, we have further numbered XM_036157063.1 as isoform 6 and XM_036157068.1 as isoform 7 in the article. To understand the isoform-specific variations, an exon-wise schematic for all five isoforms is presented in **Fig. 1D**. All five isoforms at the protein level possess two canonical RRM domains for RNA binding with variations only at C-terminal (**Fig. 1D**). To validate the RNA sequencing data, we designed a PCR primer pair that can amplify all isoforms of *Msi2*(except isoform 2) with products of different sizes. We found three different isoforms of *Msi2* to be amplified from the mouse hearts (**Supplementary Fig. 1A**). DNA sequencing of the PCR products revealed the presence of isoforms 5, 6, and 7, shown by specific exon-exon junctional reads (**Supplementary Fig. 1A-B**). However, this PCR probably did not detect *Msi2* isoforms 1and 4 due to their low expression in the heart, which is already evident from their low exon coverage. Furthermore, we confirmed the presence of all five isoforms in the murine heart by utilizing specific primers for each isoform by real-time PCR (**Fig. 1E**). *Msi2* isoforms 5, 6, and 7 were highly expressed, while isoform 4 was less expressed, and isoform 1 was the least, confirming the earlier data (**Fig. 1E**).

**Fig. 1.**
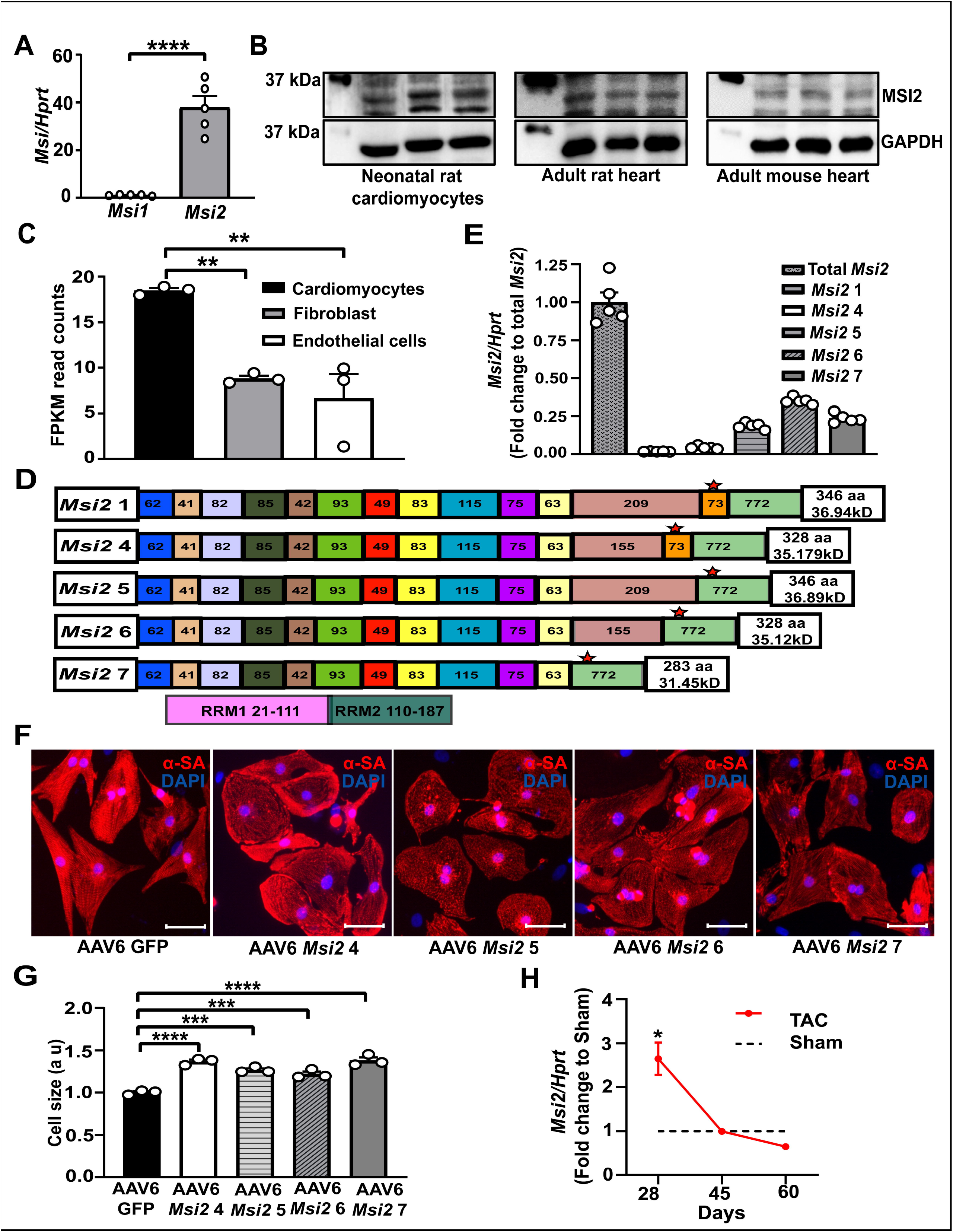
Functional characterization of *Msi2* in vitro. (**A**) Expression of Musashi family members *Msi1* and *Msi2* at mRNA level in the mouse heart (n=5). (**B**) Western blot showing the presence of MSI2 at the protein level in neonatal rat cardiomyocytes, adult rat hearts, and adult mice hearts. (**C**) FPKM read counts showing the expression level of *Msi2* in different cell fractions of the healthy heart (n=3). (**D**) Exon-wise schematic representation of Msi2 isoforms expressed in the mouse heart (region forming two RRM domains and predicted molecular weights are also shown). (**E**)Expression levels of total *Msi2* and its isoforms 1, 4, 5, 6, and 7 in mouse hearts from real-time PCR analysis (n=5). (**F**, **G**) Cell size of neonatal rat cardiomyocytes transduced with AAV6 encoding different isoforms of *Msi2* and GFP as control at MOI of 10^5^, where **F** shows representative image (n=3). (**H**) Expression level of *Msi2* in a rat pressure-overload model (TAC) of hypertrophy at different time points compared to respective sham animals. α-SA – alpha sarcomeric actinin, FPKM – fragment per kilobase of transcript per million read pairs, TAC – transverse aortic constriction. The scale bar represents 50μm. *p≤0.05, **p≤0.01, ***p≤0.001, ****p≤0.0001

Next, to study the function of *Msi2* in the heart, we overexpressed all four isoforms (4, 5, 6 and 7) found to be expressed in the heart; isoform 1 was left due to its very low expression (**Supplementary Fig. 1C**). AAV6-mediated overexpression of all *Msi2* isoforms led to a significant increase in primary neonatal rat cardiomyocyte size showing redundant function since all the isoforms have similar two RRM RNA-binding domains (**Fig. 1F-G**). The similar hypertrophic effect of *Msi2* isoforms on primary cardiomyocytes was even evident after transduction with ten times less MOI, confirming pro-hypertrophic function (**Supplementary Fig. 2**). Furthermore, human cardiac cell line AC16 also demonstrated a significant increase in cell size upon lentiviral-mediated overexpression of *Msi2* isoforms (**Supplementary Fig. 3**). These *in vitro* results suggest *Msi2* has a pro-hypertrophic function. Therefore, we checked the levels of MSI2 in primary cardiomyocytes and hearts during the hypertrophic condition. MSI2 was significantly upregulated in primary cardiomyocytes treated with isoproterenol (**Supplementary Fig. 4**). Similar to neonatal rat primary cardiomyocytes, the *Msi2* expression level was significantly induced after four weeks of pressure-overload-induced hypertrophy in rats (**Fig. 1H**). However, *Msi2* expression follows a downward trend during longer durations of pressure overload (**Fig. 1H**). These results indicate a pro-hypertrophic role for *Msi2*.

### 2. *In vivo* cardiac overexpression of *Msi2* causes heart-failure

Next, to confirm the *in vitro* pro-hypertrophic effect of *Msi2*, we selected two *Msi2* isoforms 4 and 7. One of the apparent differences between abundantly expressed isoforms (5,6,7) and low expressed isoforms (1,4) is the exclusion and inclusion of the penultimate exon of 73 base pairs, respectively, leading to differences at the protein c-terminal (**Fig. 1D**). Therefore, isoform 4 was selected from low expressed as it includes the penultimate exon. Isoform 7 was chosen among the abundant isoforms due to its uniqueness of excluding both alternatively spliced penultimate exons (209/155 base pairs and 73 base pairs), providing an opportunity to elucidate the effects of these exons on MSI2 cardiac functions (**Fig. 1D**). To study the *in vivo* effects, we injected adult mice with 1.8*10^12^ AAV9-*Msi2* 4/7 and AAV9-GFP control as previously done [12]. MSI2 isoforms 4 and 7 were successfully overexpressed in the heart injected with AAV9-*Msi2* compared to controls (**Fig. 2A-C**). Both isoforms of *Msi2* led to early death in mice compared to the control (**Fig. 2D**). *Msi2* isoform 4 and 7 overexpression caused massive cardiac hypertrophy evident from the gross morphology (**Fig. 2E**) and significantly increased heart weight to tibia length ratio (**Fig. 2F**). AAV9-*Msi2* transduced hearts revealed heart-failure phenotype during the echocardiographic analysis of cardiac function (**Fig. 2G-K**). *Msi2* overexpressed hearts displayed significantly reduced ejection fraction (**Fig. 2G**), fractional shortening (**Fig. 2H**), and cardiac output (**Fig. 2I**). Additionally, increased left ventricular mass (**Supplementary Fig. 5A**), left ventricular end-systolic diameter (**Fig. 2J**), left ventricular end-diastolic diameter (**Fig. 2K**), left ventricular systolic volume (**Supplementary Fig. 5B**), and left ventricular diastolic volume (**Supplementary Fig. 5C**) was found suggesting hypertrophy together with dilation. Next, we measured the cardiomyocyte cross-sectional area in heart sections to confirm the hypertrophy by wheat germ agglutinin staining. *Msi2* overexpressing cardiomyocytes depicted increased area (**Fig. 3A-B**), demonstrating cardiac hypertrophy. Furthermore, *Msi2* overexpressing hearts showed increased fibrosis by picrosirus red staining under bright fields (**Fig. 3C-D**). At the molecular level, *Msi2* overexpression in hearts activated the cardiac remodeling gene program, as seen by significantly increased *Nppa*, *Nppb*, and *Myh7/Myh6* levels (**Fig. 3E-G**). Our data show that cardiac overexpression of *Msi2* leads to heart failure due to hypertrophy, dilation, and activation of pathological cardiac remodeling.

**Fig. 2.**
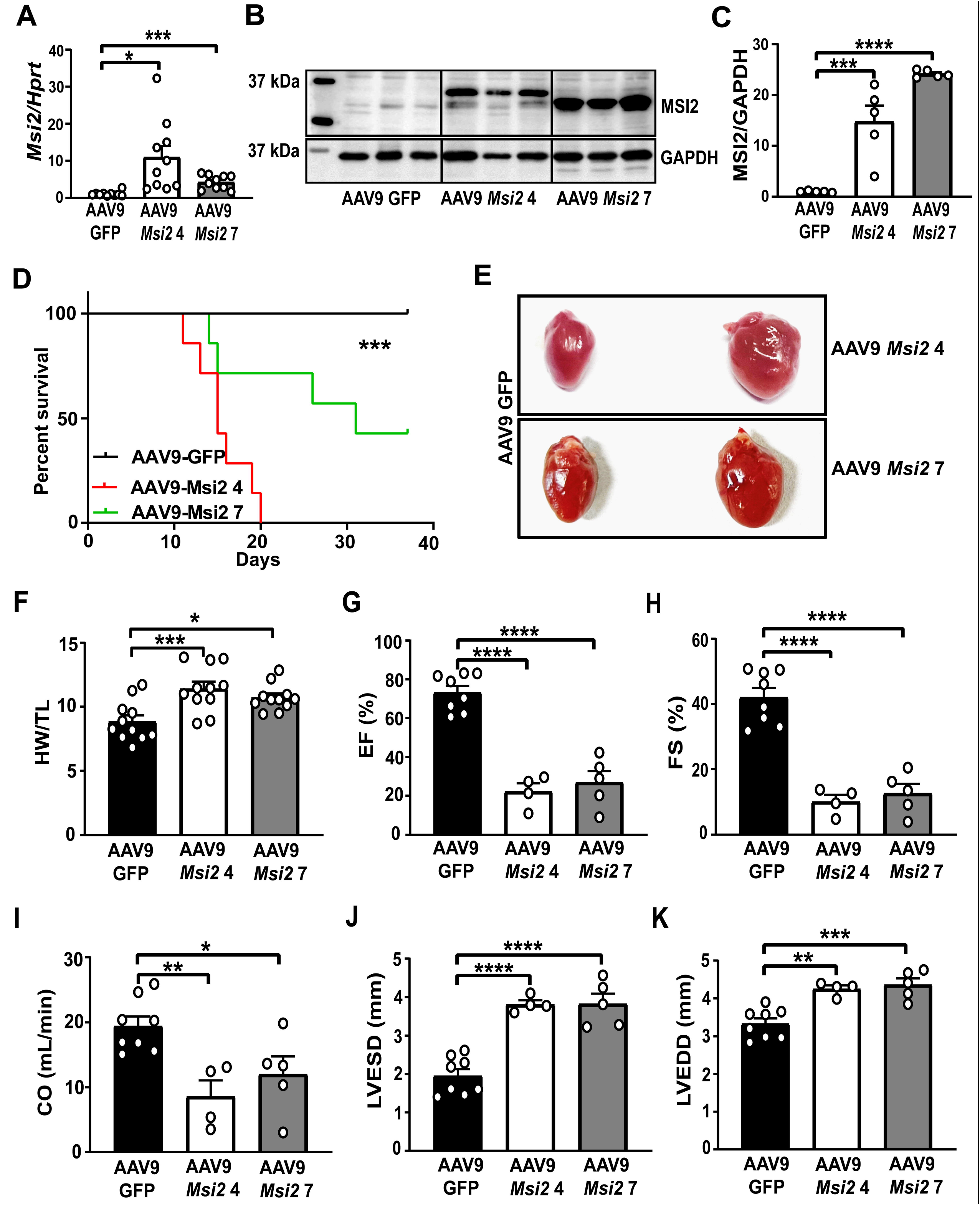
AAV9-mediated overexpression of *Msi2* causes heart failure. (**A**) The expression level of *Msi2* at mRNA in murine hearts overexpressing MSI2 or GFP control. (**B**, **C**) Western blot showing MSI2 and GAPDH in heart lysate from AAV9-injected animals encoding for *Msi2* isoform 4 and 7 and GFP as control (n=5). (**D**) Kaplan-Meier survival curve for mice injected with AAV9 encoding *Msi2* isoform 4 and 7 and GFP as control (n=7). (**E**) Representative heart images with overexpression of *Msi2* isoform 4 and 7 and GFP. (**F**) Heart weight to tibia length ratio of mice treated with AAV9 encoding *Msi2* isoform 4 and 7 and GFP (n=11). (**G**-**K**) Echocardiographic parameters showing the cardiac function of AAV9-*Msi2* isoform 4 and 7 treated animals (n=8 GFP, n=4 *Msi2* 4, n=5 *Msi2* 7). HW-heart weight, TL-tibia length, EF-ejection fraction, FS – fractional shortening, CO – cardiac output, LVESD – left ventricular end-systolic diameter, LVEDD – left ventricular end-diastolic diameter. *p≤0.05, **p≤0.01, ***p≤0.001, ****p≤0.0001

**Fig. 3.**
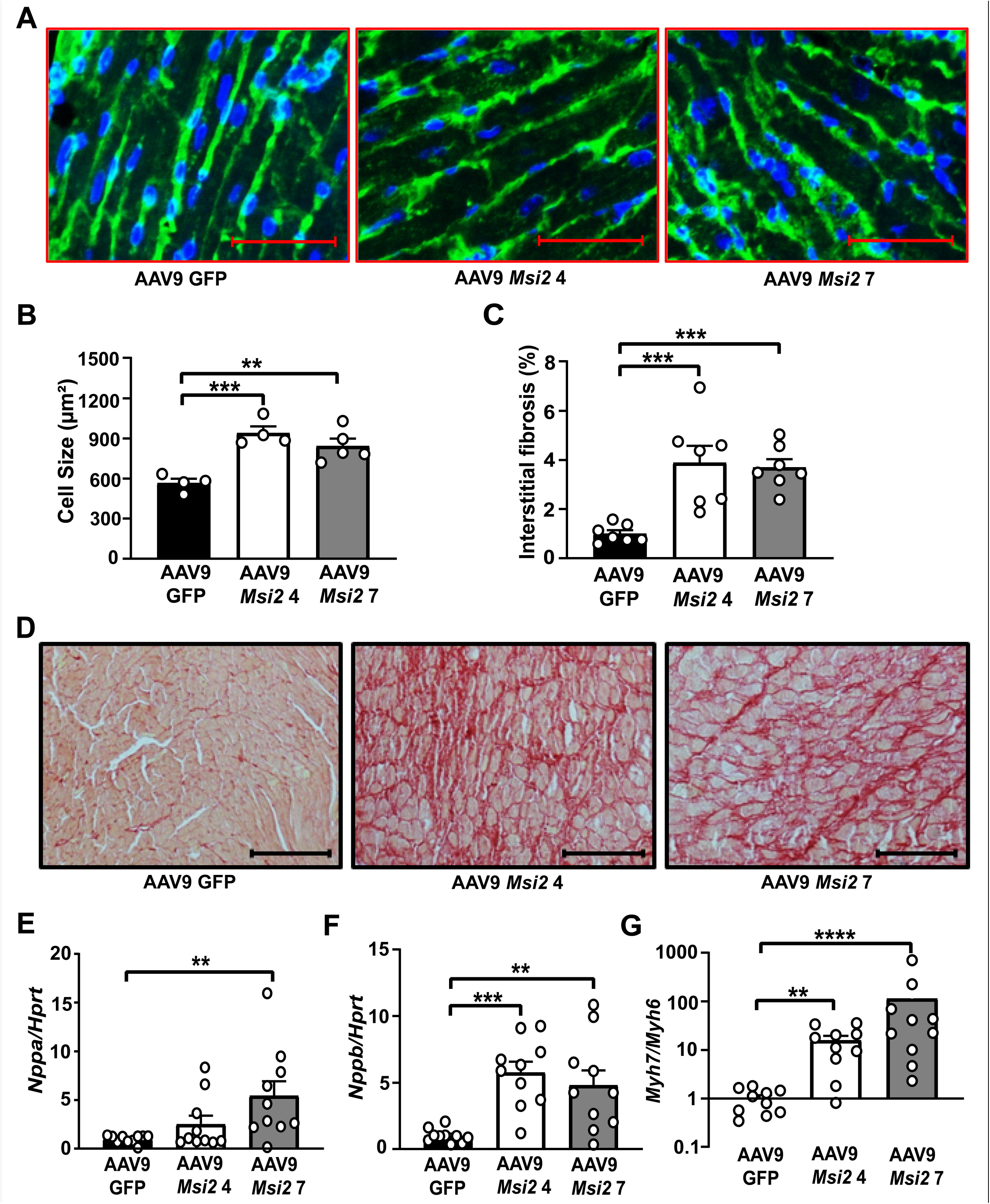
*Msi2* induces pathological remodeling of the heart. (**A**, **B**) Cardiomyocyte size measurement in heart sections of AAV9-*Msi2* isoform 4 and 7 and GFP using wheat germ agglutinin staining (n=4-5). (**C**). (C, **D**) Fibrosis measurement by picrosirius staining of hearts overexpressing *Msi2* isoform 4 and 7 and GFP (n=7). (**E**-**G**) Expression levels of *Nppa* (**E**), *Nppb* (**F**), and *Myh7/Myh6* (**G**) mRNA in hearts overexpressing *Msi2* isoform 4 and 7 and GFP (n=10). μm – micrometer. The scale bar for **3A** represents 50μm, and the scale bar for **3D** represents 100μm. *p≤0.05, **p≤0.01, ***p≤0.001, ****p≤0.0001

### 3. Overexpression of *Msi2* causes global downregulation of nuclear-encoded mitochondrial genes

*Msi2* functions in diverse ways, from regulating pri-microRNA processing, mRNA decay, and translational inhibition depending upon its localization in cells [1, 5, 14]. Therefore, to explore the cardiac functions of *Msi2*, we checked the cellular localization of MSI2 protein in human cardiac cell line AC16 by immunofluorescence. Basal MSI2 and its isoforms 4 and 7 were found mainly localized in the cytoplasm (**Supplementary Fig. 6A**) in the AC16 cell line. Furthermore, MSI2 localization was also checked in murine heart sections. MSI2 was primarily found to be localized in the cytoplasm at the basal level and after overexpression (**Supplementary Fig. 6B**). Based on cytoplasmic localization, *Msi2* can be predicted to function either as an inducer of mRNA decay or a translational inhibitor [1, 14]. As *Msi2* could regulate gene expression at the RNA or protein level, we performed an LC-MS/MS-based global proteomic profiling of AAV9-*Msi2* 4 overexpressed and control hearts. Proteomic profiling identified 3170 proteins, with 2692 proteins common to both groups, while others were unique (**Fig. 4A**). Principal component analysis of the proteomics data exhibited two completely separated clusters representing each group (**Fig. 4B**). *Msi2* overexpression significantly altered 1037 proteins, with 495 upregulated and 542 downregulated (**Fig. 4C-D**) (**Supplementary Table 4**).

**Fig. 4.**
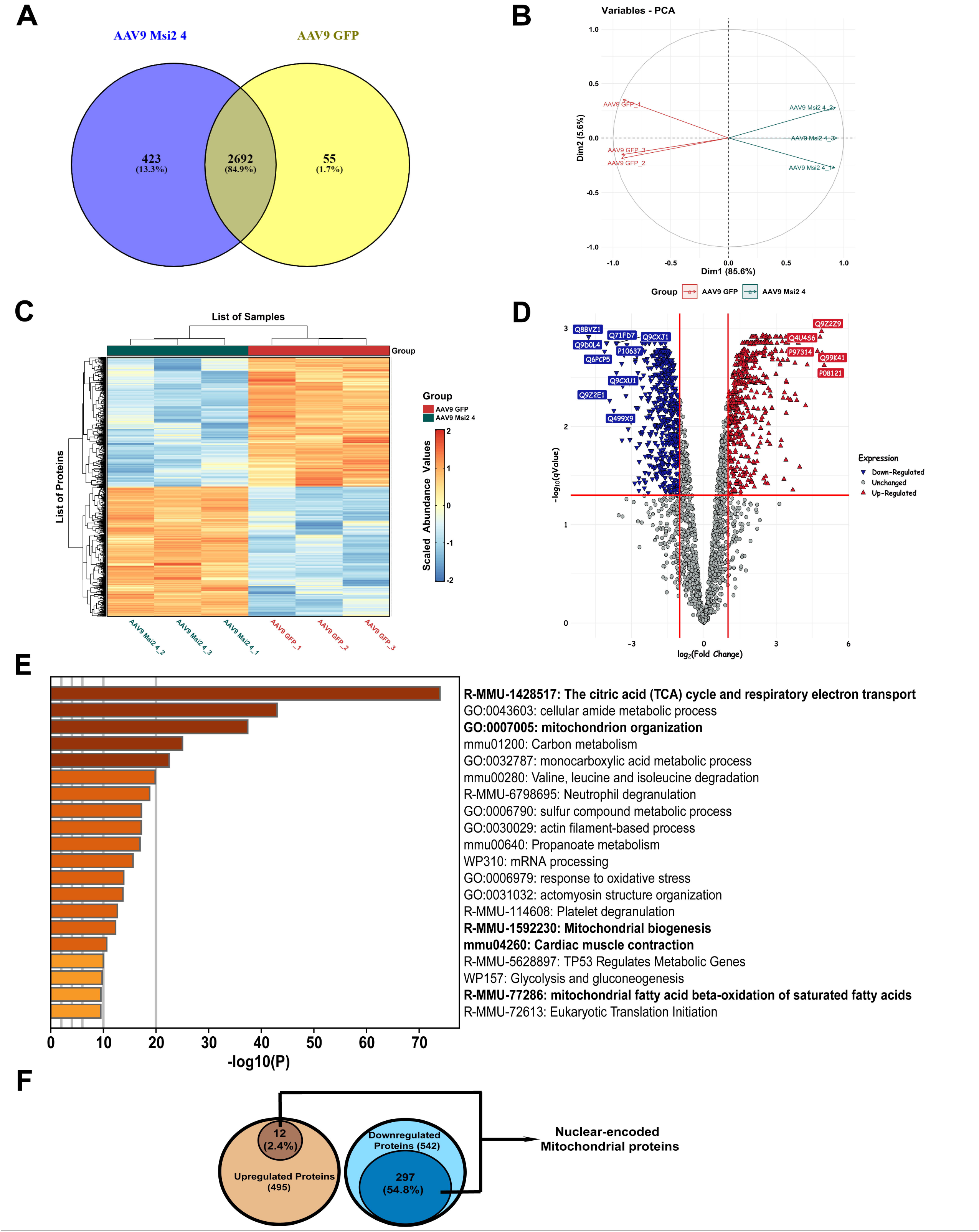
*Msi2* induced deregulation of the cardiac proteome. (**A**-**D**) Global proteomics analysis in heart lysate from AAV9-*Msi2* 4 and AAV9-GFP shows the number of proteins identified (**A**), principal component analysis of complete proteome (**B**), heatmap (**C**), and a volcano plot (**D**) showing differentially expressed genes with fold change cut-off of two and p-value ≤0.05 (n=3). The scale bar represents 50μm. (**E**) Gene ontology (GO) term analysis of differentially expressed proteins after *Msi2* 4 overexpression in the heart (mitochondria-related terms are highlighted in bold). (**F**) Schematics show the number of nuclear-encoded mitochondrial proteins among up and downregulated proteins from the proteomics data.

We performed gene ontology (GO) analysis using Metascape online tool to understand the significance of differentially expressed proteins. GO analysis revealed enrichment of several processes related to mitochondria like mitochondrion organization, mitochondrial biogenesis, mitochondrial fatty acid beta-oxidation of saturated fatty acids, TCA cycle, and respiratory electron transport, and cardiac muscle contraction suggestive of dysfunctional mitochondria and energy-deprived failing heart (**Fig. 4E**). Mitochondria only encode thirteen protein-coding genes and depend on several nuclear-encoded proteins for proper functioning [28]. Interestingly, among the downregulated proteins in *Msi2* overexpressing hearts, more than half were nuclear-encoded mitochondrial proteins (**Fig. 4F and Supplementary Fig. 7**), thus supporting the GO outcome of mitochondrial dysfunction. Next, to validate the GO outcome of mitochondrial dysfunction in MSI2 overexpressing hearts, we performed transmission electron microscopic (TEM) imaging of AAV9-*Msi2* isoform 4 and 7 treated hearts and control. Using the thin sectioning TEM technique, we demonstrated a disorganized arrangement of mitochondria and mitochondria with wider and less packaged cristae with bulge tips in *Msi2* overexpressing hearts (**Fig. 5A**).

**Fig. 5.**
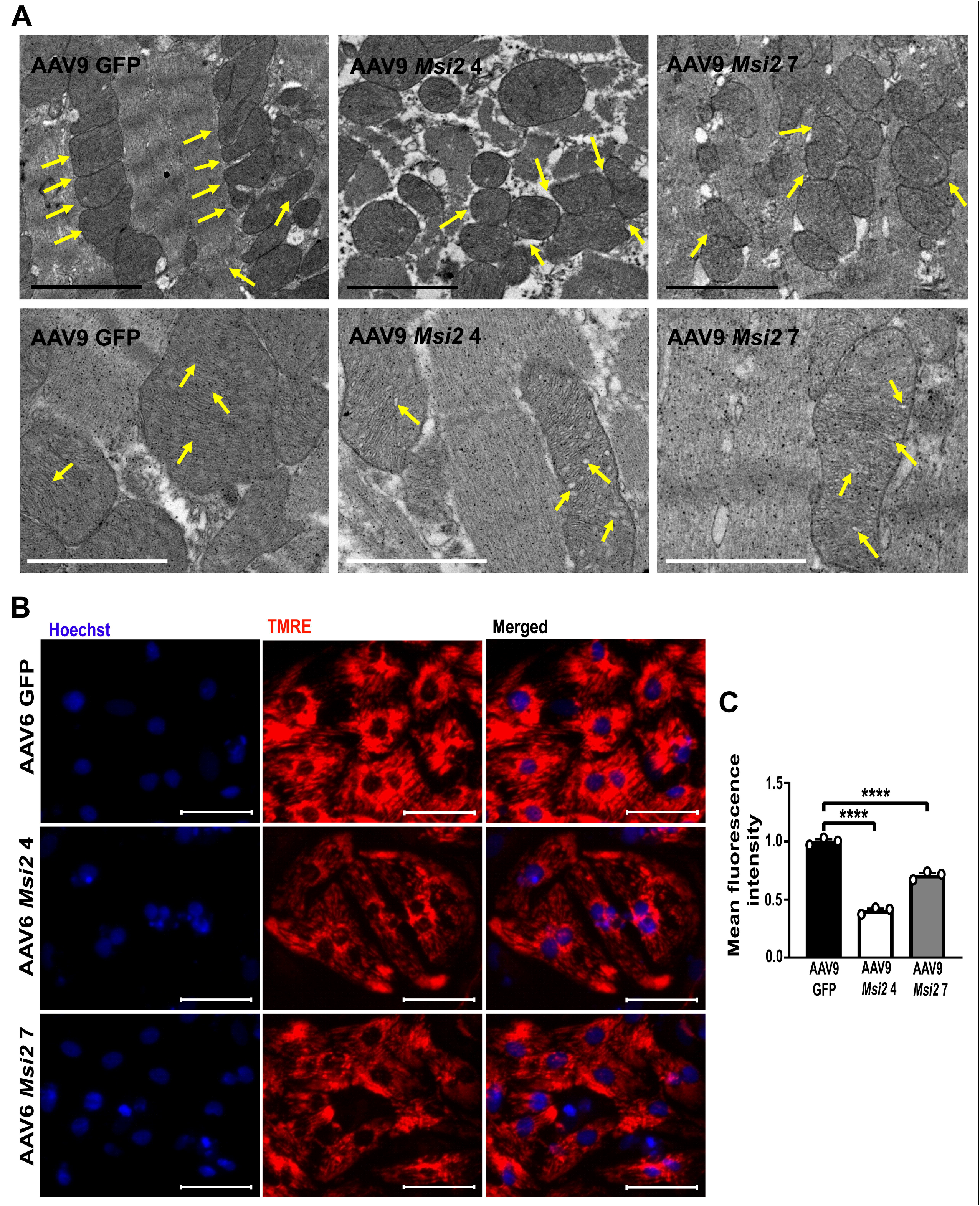
*Msi2* induces mitochondrial dysfunction. (**A**) TEM micrographs of mouse hearts with overexpression of *Msi2* 4 and 7 and GFP control (arrows highlight mitochondrial distribution and structural changes observed). The black scale bar represents 2μm while the white one represents 1 μm. (**B, C**) TMRE staining showing mitochondrial transmembrane potential (function) in primary cardiomyocyte overexpressing *Msi2* isoforms 4 and 7 and GFP. The scale bar represents 50μm. ****p≤0.0001

On the contrary, control AAV9-GFP treated hearts have a regular mitochondrial arrangement parallel to sarcomeres with preserved structural integrity and tightly packed cristae, thus confirming mitochondrial dysfunction in MSI2 overexpressing hearts (**Fig. 5A**). Mitochondrial dysfunction is a common phenomenon in failing hearts. Therefore, to verify that the mitochondrial dysfunction phenotype is a direct effect of MSI2 overexpression and is not confounded by the underlying failing cardiac phenotype, we measured the mitochondrial transmembrane potential with TMRE red dye *in vitro* in primary cardiomyocytes. After overexpression of *Msi2* 4 and 7, primary cardiomyocytes showed significantly reduced red fluorescence confirming lower mitochondrial transmembrane potential, compared to GFP control (**Fig. 5B-C**). These results demonstrate that increased levels of both *Msi2* isoforms 4 and 7 induce mitochondrial dysfunction leading to cardiac malfunctioning.

### 4. *Msi2* targets nuclear-encoded regulatory factors *Cluh* and *Smyd1*

MSI2 is known to directly bind to the 3’UTR of its targets and destabilize them, leading to the downregulation of the target at mRNA and protein. Hence, to identify downstream targets of MSI2, we focussed on downregulated proteins from proteomics. Based on the proteomics data showing enrichment for nuclear-encoded mitochondrial proteins among downregulated proteins, we hypothesize that MSI2 may directly target broad regulators, which may affect the expression of nuclear-encoded mitochondrial genes at the transcriptional or post-transcriptional level. Therefore, we checked the molecular function of all downregulated proteins from GO (gene ontology) molecular function and identified seventy-two proteins with DNA or RNA binding and transcriptional or translational regulatory function (**Supplementary Table 5**). Next, among these proteins, we identified two nuclear-encoded broad regulatory factors, *Cluh* and *Smyd1*, which are known to control mitochondrial structure and function. *Cluh* is an RNA-binding protein that regulates the stability and translation of various nucleus-encoded mitochondrial genes [29]. *Smyd1* is a muscle-specific histone methyltransferase known to regulate mitochondrial energetics by controlling the expression of several regulatory factors [20, 33]. Hence, these proteins may act as downstream targets of *Msi2*, controlling the expression of several nucleus-encoded mitochondrial genes. Like proteomics, AAV9-*Msi2* 4 and 7 treated hearts have significantly lower *Cluh* and *Smyd1* mRNA levels than the control (**Fig. 6A-B**). Furthermore, overexpression of both isoforms of *Msi2* in primary cardiomyocytes also significantly downregulated *Cluh* and *Smyd1* levels, showing them as putative targets of *Msi2* (**Fig. 6C-D**). Next, we assessed whether lower levels of *Cluh* and *Smyd1* mRNAs are due to destabilization induced by MSI2. Primary cardiomyocytes overexpressing MSI2 treated with actinomycin D showed significantly lower levels of *Cluh* and *Smyd1* compared to the control (**Fig. 6E**). Thus declined levels of *Cluh* and *Smyd1* mRNAs are due to the effect induced by MSI2 on their stability. Additionally, *Hadha*, *Pcca*, *Pdha1*, *Acat1*, and *Mccc1* validated targets of *Cluh*, and *Ppargc1a* and *Perm1* validated targets of *Smyd1* were significantly downregulated in *Msi2* overexpressing hearts (**Fig. 6F-G**). These results indicate *Cluh* and *Smyd1* as the potential downstream target of *Msi2*, regulated by mRNA decay. *Msi2* recognizes its target mRNA for degradation by the presence of multiple copies of the UAG sequence in its 3’UTR [1]. In silico analysis of 3’UTR of *Cluh* and *Smyd1* revealed eight and twenty-one putative UAG binding motifs, respectively. Therefore to confirm *Cluh* and *Smyd1* as the direct targets, we cloned 3’UTR of both genes downstream to luciferase. The overexpression of *Msi2* isoforms 4 and 7 significantly decreased luciferase activity with *Cluh* and *Smyd1* 3’UTR, confirming them as direct targets (**Fig. 6H-I**). Furthermore, to ensure that the decline in luciferase activity is due to the direct binding of MSI2 to the *Cluh* and *Smyd1* 3’UTR, we performed ribonucleoprotein immunoprecipitation of MSI2. Significant enrichment of *Cluh* and *Smyd1* 3’UTR regions was found in the MSI2 pull-down fraction compared to IgG control, validating *Cluh* and *Smyd1* as the direct downstream target of MSI2 (**Fig. 6J**). Furthermore, overexpression of either *Cluh* or *Smyd1* prevented *Msi2*-induced cardiac hypertrophy in primary cardiomyocytes (**Fig. 7A-B**). Additionally, mitochondrial dysfunction induced by MSI2 was partially rescued by overexpression of downstream targets *Cluh* and *Smyd1* (**Fig. 7C-D**). Thus, these results establish *Cluh* and *Smyd1* as direct downstream targets of *Msi2*.

**Fig. 6.**
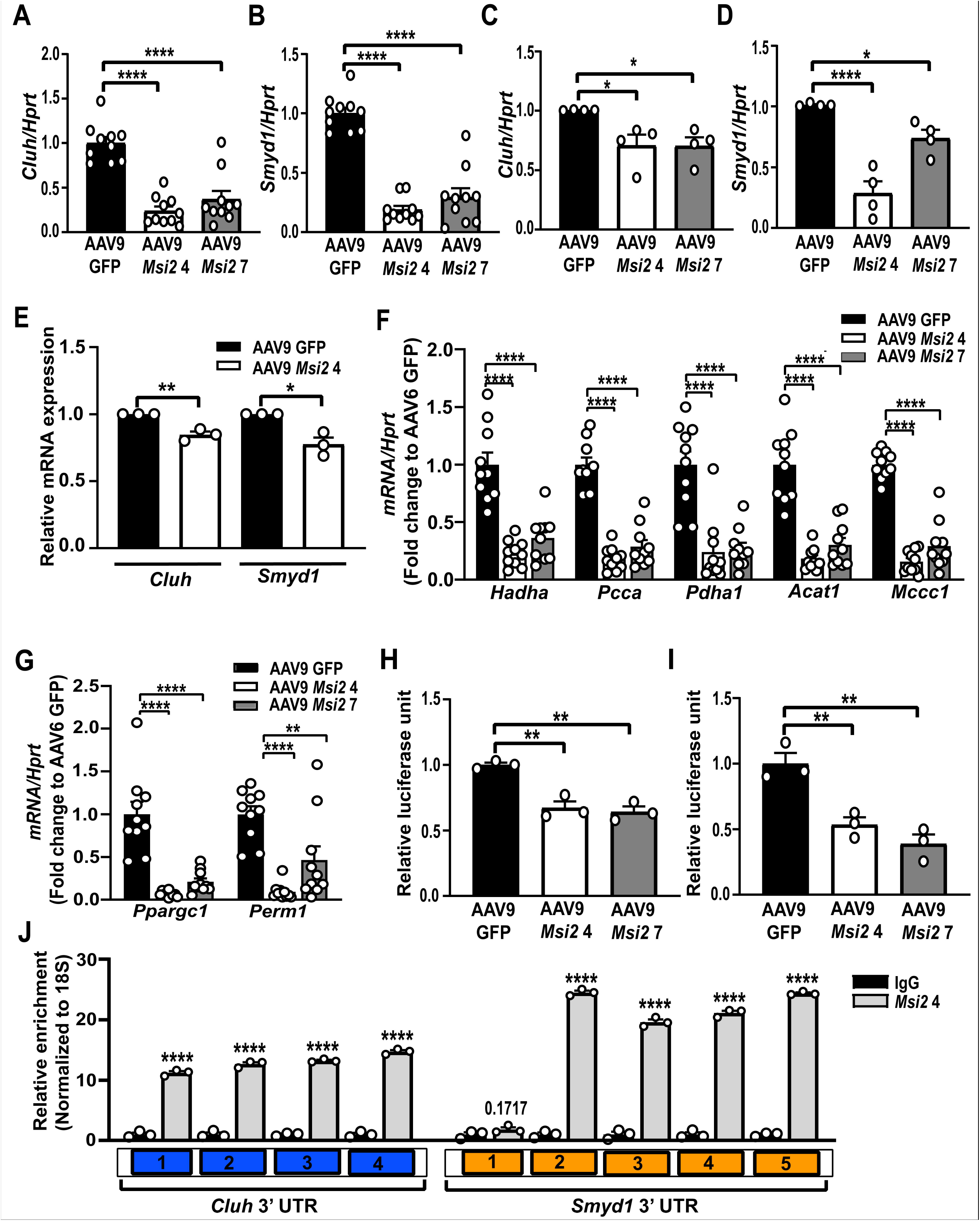
*Cluh* and *Smyd1* are novel targets of *Msi2*. (**A**, **B**) Expression levels of *Cluh* and *Smyd1* mRNA in hearts overexpressing *Msi2* 4 and 7 and GFP control (n=10). (**C**, **D**) Cluh and Smyd1 mRNA expression levels in neonatal rat cardiomyocytes after AAV6-induced overexpression of *Msi2* isoform 4 and 7 (n=4). (**E**) Expression level of *Cluh* and *Smyd1* in neonatal rat cardiomyocytes transduced with AAV6-*Msi2* 4 after four to six hours of Actinomycin D treatment. (**F**) Expression levels of *Cluh* target genes *Hadha*, *Pcca*, *Pdha1*, *Acat1*, and *Mccc1* at mRNA in hearts overexpressing *Msi2* 4 and 7 and GFP control (n=10). (**G**) Expression levels of *Ppargc1a* and *Perm1* (targets of *Smyd1*) in hearts with overexpression of *Msi2* isoform 4 and 7 (n=10). (**H**, **I**) Luciferase reporter assay with 3’UTR of *Cluh* (**H**) and *Smyd1* (**I**) after overexpression of *Msi2* isoform 4 and 7 and GFP control (n=3). (**J**) Enrichment level of different regions of Cluh and Smyd1 3’UTR in MSI2 or IgG pull-down RNA. *p≤0.05, **p≤0.01, ****p≤0.0001

**Fig. 7.**
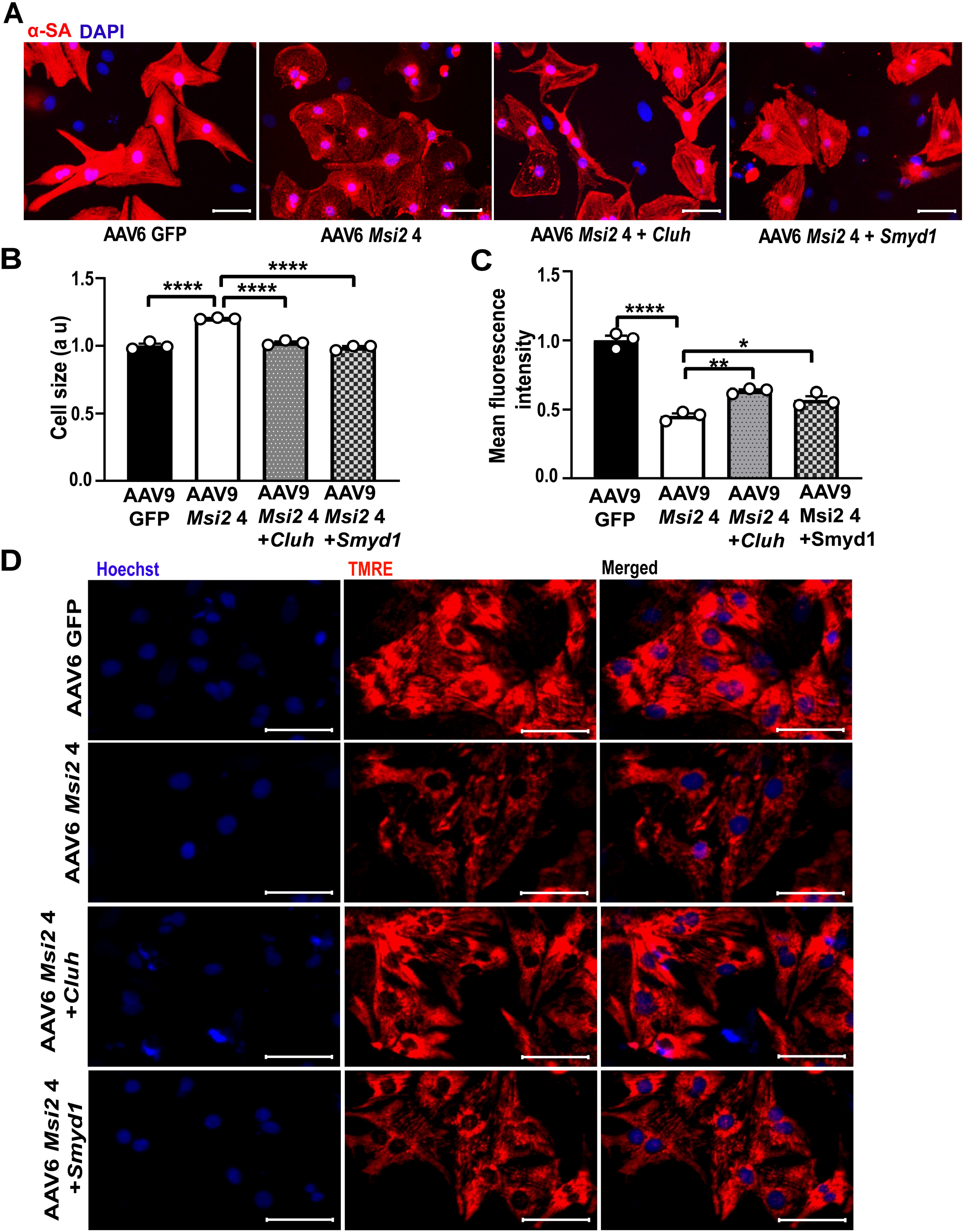
CLUH and SMYD1 are mediators of the MSI2 function. (**A, B**) Neonatal rat cardiomyocyte cell size after AAV6-mediated overexpression of *Msi2* isoform 4 alone or with *Cluh* or *Smyd1* (n=3). (**C, D**) TMRE staining showing mitochondrial transmembrane potential (function) in primary cardiomyocyte overexpressing *Msi2* isoforms 4 alone or together with *Cluh* or *Smyd1* (n=3). Au – arbitrary unit, α-SA – alpha sarcomeric actinin. The scale bar represents 50μm. *p≤0.05, **p≤0.01, ****p≤0.0001

## Discussion

A cell’s amount of translatable mRNA depends upon its synthesis and decay [30]. The purpose of mRNA decay is either to function as a quality control mechanism to eliminate unwanted proteins or to regulate the abundance of proteins by altering their half-lives [30]. RBPs regulate mRNA half-lives and their translation by forming mRNA ribonucleoprotein complexes [30]. RBPs involved in these processes mainly bind to mRNA 3’UTRs through specific sequence motifs and can promote or hinder the binding of decay factors or regulate translatable status [30]. RBPs-mediated control of mRNA decay and translation plays a pivotal role in cardiovascular development and disease; however, its role in cardiovascular diseases remains less explored [9]. Recently, Zhou A et al. have shown that an RBP HuR increases levels of *Mef2c* in cardiomyocytes by binding and stabilizing its mRNA through ARE elements in 3’ UTR [36]. Likewise, another RBP, *Pcbp2*, was shown to have an anti-hypertrophic function in cardiomyocytes by promoting the degradation of *Gpr56* mRNA [35]. Similarly, RBP *Celf1* binds to *Cx43* (connexion 43) mRNA by binding UG-rich regions in its 3’ UTR, promoting dilated cardiomyopathy [3]. Here, we demonstrate a novel pro-hypertrophic cardiac function for an RBP *Msi2* via regulating mRNA decay. MSI2 was found to be expressed in mice and rat hearts, and rat primary cardiomyocytes at protein level. Similar to our results MSI2 was found to be expressed at protein level in heart by Sakakibara et al. [27]. Recently, Reichert et al. have shown enrichment of MSI2 protein in cardiomyocyte RBPome (RNA-binding proteome) confirming RNA-binding activity of MSI2 in cardiomyocytes [22]. Additionally, we found *Msi2* to be enriched in cardiomyocytes compared to other cell types suggesting an important cardiac function.

*Msi2* isoform 4 and 7 overexpression in heart-induced cardiac hypertrophy, dilation, heart failure, and death. In line with the hypertrophic effect, *Msi2* was found to be significantly increased in pressure-overload model of cardiac hypertrophy. Global proteomics analysis demonstrated the downregulation of several nuclear-encoded mitochondrial proteins after *Msi2* overexpression. Electron microscopic imaging in heart sections and *in vitro* measurement of transmembrane potential in cardiomyocytes confirmed mitochondrial dysfunction. Mechanistically, *Cluh* and *Smyd1*, broad regulators of mitochondrial structure and functions, were identified as novel direct targets of *Msi2. Msi2* destabilizes *Cluh* and *Smyd1* mRNAs by directly binding to their 3’UTR. Overexpression of either target *Cluh* or *Smyd1* partially prevented *Msi2*-induced cardiac hypertrophy and mitochondrial dysfunction in primary cardiomyocytes. Thus, confirming *Cluh* and *Smyd1* as direct targets and downstream mediators of *Msi2* pro-hypertrophic function (summarized in **Fig. 8**).

**Fig. 8.**
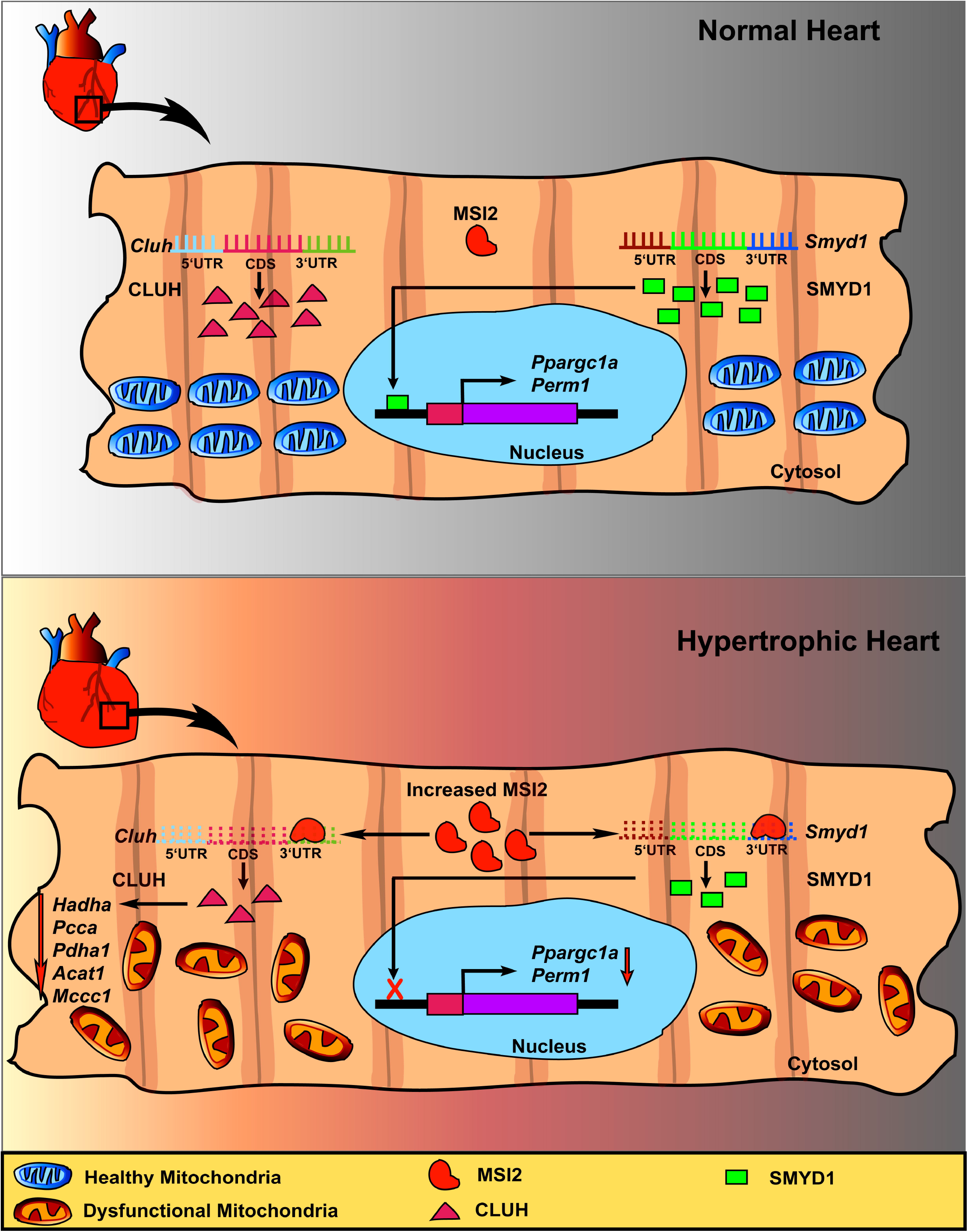
Schematic summarizing the function of MSI2 in healthy and hypertrophic hearts.

Both *Cluh* and *Smyd1* regulate mitochondrial structure and function by controlling the expression of several nuclear-encoded mitochondrial genes. *Cluh* is an RBP that helps survive post-birth starvation in neonates, and knockout mice die shortly after birth due to hypoglycemia [29]. Liver-specific knockout in adult liver shows its role in metabolic adaptation during conditions of high-energy demand like starvation [29]. *Cluh* exerts its metabolic function by promoting stability and translation of several nuclear-encoded mitochondrial genes [10, 29]. Some known targets of Cluh, like HADHA, PCCA, PDHA1, ACAT1, and MCCC1, were downregulated in our cardiac proteomics data and at the mRNA level, illustrating the probable existence of similar molecular mechanisms in the heart. We also showed that *Cluh* could partially inhibit *Msi2*-induced cardiac hypertrophy and mitochondrial dysfunction, demonstrating a need to explore the cardiac role of *Cluh* further. Another *Msi2* target, *Smyd1*, is a muscle-specific histone methyltransferase whose deletion in the adult heart causes cardiac hypertrophy and heart failure [7]. Loss of *Smyd1* in the adult heart results in mitochondrial dysfunction due to the downregulation of *Ppargc1a* and *Perm1* [20, 33]. Similarly, we have found decreased *Ppargc1a* and *Perm1* upon *Msi2*-induced downregulation of *Smyd1* in the heart.

*Msi2* has different isoforms due to alternative splicing [18]. Here, we confirmed the expression of two novel predicted isoforms, XM_036157063.1 (isoform 6) and XM_036157068.1 (isoform 7), together with isoforms 5, 4, and 1 in the heart. All these three isoforms (5, 6, and 7) highly expressed in the heart have a common exclusion of the penultimate exon of 73 base pairs. On the other hand, isoforms 4 and 1, which are low expressed, include this 73 base pair exon. Similarly, in a different study from our lab, alternative splicing analysis from RNA-seq data of balbc hearts demonstrated an inclusion-to-exclusion ratio of 0.15 for this 73 base pairs exon, confirming exon exclusion as a dominant phenotype in the heart (**data not shown**).

Collectively, we have shown a novel pro-hypertrophic function for *Msi2* in the heart leading to mitochondrial dysfunction, heart failure, and death in mice. Furthermore, we identified two novel targets of *Msi2*, namely *Cluh* and *Smyd1* (**Fig. 8**). Additionally, we validated the expression of two predicted *Msi2* isoforms in the heart.

## Non-standard Abbreviations and Acronyms

RBP: RNA-binding protein
AAV: Adeno-associated virus
GFP: Green fluorescent protein
UTR: Untranslated region
DNA: Deoxyribonucleic acid
RNA: Ribonucleic acid
rRNA: Ribosomal RNA
tRNA: Transfer RNA
mRNA: messenger RNA
NCBI: National Center for Biotechnology Information
PCR: Polymerase chain reaction
DAPI: 4’,6-diamidino-2-phenylindole
TEM: Transmission Electron Microscope
RRM: RNA recognition motifs
LC-MS/MS: Liquid Chromatography with tandem mass spectrometry
GO: Gene Ontology
TCA: Tricarboxylic Acid
ARE: Adenylate-uridylate-rich elements
TMRE: Tetramethylrhodamine, ethyl ester
TAC: Trans-aortic constriction
FPKM: Fragment per kilobase of transcript per million read pairs
MOI: Multiplicity of Infection

## Acknowledgments

We want to thank Dr. Rajdeep Guha, Head of Animal Facility, CSIR-CDR, for his help with animal experiments. We want to thank Dr. Manoj Kumar Barthwal, Dr. Chandra Prakash Pandey, and Dr. Hobby Aggarwal, Pharmacology Division, for their help with the instruments and training. We acknowledge the FACS facility, CSIR-CDRI, for using their FACS machine. We acknowledge the valuable suggestions of Dr. Regalla Kumarswamy, CSIR-Centre for Cellular and Molecular Biology, Hyderabad, India, Dr. Marisol Ruiz-Meana, Valld’Hebron Hospital Universitari, Barcelona, Spain, and Dr. Gaurav Ahuja, Indraprastha Institute of Information Technology, Delhi, India.

## Funding

This work was funded by Ramalingaswami Re-entry Fellowship (BT/RLF/re-entry/14/2019) from the Department of Biotechnology, Government of India, to SKG. SS, SP, SK, and RKS avail JRF fellowship from CSIR (Council of Scientific and Industrial Research), Government of India. RK avails SRF fellowship from ICMR (Indian Council of Medical Research), and AG and ADC avail JRF fellowship from UGC (University Grant Commission), Government of India.

### Disclosure

TT has filed and licensed patents about non-coding RNAs and is the founder and shareholder of Cardior Pharmaceuticals GmbH (outside of this manuscript). SKG holds patents about non-coding RNAs (outside of this manuscript).

### Author contribution

SKG has developed the concept, designed the study, planned experiments, analyzed results, and prepared the manuscript. SS designed the study, performed most experiments, analyzed the results, and drafted the manuscript. AG, RK, SP, SK, ADC, and PP helped with neonatal rat cardiomyocyte isolation, lentivirus, and AAV production. RKS and KM performed electron microscopy experiments on heart samples. PP and KJ helped with echocardiography and animal experiments. PB and KH provided the rat TAC heart samples. PC and SK performed the RNA sequencing analysis. TT gave critical inputs and helped in manuscript writing.

## References

1. Bennett CG, Riemondy K, Chapnick DA, Bunker E, Liu X, Kuersten S, Yi R (2016) Genome-wide analysis of Musashi-2 targets reveals novel functions in governing epithelial cell migration. Nucleic Acids Res 44:3788–3800 doi:10.1093/nar/gkw207

2. Bhattacharjee A, Hasanain M, Kathuria M, Singh A, Datta D, Sarkar J, Mitra K (2018) Ormeloxifene-induced unfolded protein response contributes to autophagy-associated apoptosis via disruption of Akt/mTOR and activation of JNK. Sci Rep 8:2303 doi:10.1038/s41598-018-20541-8

3. Chang KT, Cheng CF, King PC, Liu SY, Wang GS (2017) CELF1 Mediates Connexin 43 mRNA Degradation in Dilated Cardiomyopathy. Circ Res 121:1140–1152 doi:10.1161/CIRCRESAHA.117.311281

4. Chen S, Zhou Y, Chen Y, Gu J (2018) fastp: an ultra-fast all-in-one FASTQ preprocessor. Bioinformatics 34:i884–i890 doi:10.1093/bioinformatics/bty560

5. Choudhury NR, de Lima Alves F, de Andres-Aguayo L, Graf T, Caceres JF, Rappsilber J, Michlewski G (2013) Tissue-specific control of brain-enriched miR-7 biogenesis. Genes Dev 27:24–38 doi:10.1101/gad.199190.112

6. Corbett AH (2018) Post-transcriptional regulation of gene expression and human disease. Curr Opin Cell Biol 52:96–104 doi:10.1016/j.ceb.2018.02.011

7. Franklin S, Kimball T, Rasmussen TL, Rosa-Garrido M, Chen H, Tran T, Miller MR, Gray R, Jiang S, Ren S, Wang Y, Tucker HO, Vondriska TM (2016) The chromatin-binding protein Smyd1 restricts adult mammalian heart growth. Am J Physiol Heart Circ Physiol 311:H1234–H1247 doi:10.1152/ajpheart.00235.2016

8. Froese N, Cordero J, Abouissa A, Trogisch FA, Grein S, Szaroszyk M, Wang Y, Gigina A, Korf-Klingebiel M, Bosnjak B, Davenport CF, Wiehlmann L, Geffers R, Riechert E, Jurgensen L, Boileau E, Lin Y, Dieterich C, Forster R, Bauersachs J, Ola R, Dobreva G, Volkers M, Heineke J (2022) Analysis of myocardial cellular gene expression during pressure overload reveals matrix based functional intercellular communication. iScience 25:103965 doi:10.1016/j.isci.2022.103965

9. Gao C, Wang Y (2020) mRNA Metabolism in Cardiac Development and Disease: Life After Transcription. Physiol Rev 100:673–694 doi:10.1152/physrev.00007.2019

10. Gao J, Schatton D, Martinelli P, Hansen H, Pla-Martin D, Barth E, Becker C, Altmueller J, Frommolt P, Sardiello M, Rugarli EI (2014) CLUH regulates mitochondrial biogenesis by binding mRNAs of nuclear-encoded mitochondrial proteins. J Cell Biol 207:213–223 doi:10.1083/jcb.201403129

11. Guo QQ, Gao J, Wang XW, Yin XL, Zhang SC, Li X, Chi LL, Zhou XM, Wang Z, Zhang QY (2020) RNA-Binding Protein MSI2 Binds to miR-301a-3p and Facilitates Its Distribution in Mitochondria of Endothelial Cells. Front Mol Biosci 7:609828 doi:10.3389/fmolb.2020.609828

12. Gupta SK, Garg A, Avramopoulos P, Engelhardt S, Streckfuss-Bomeke K, Batkai S, Thum T (2019) miR-212/132 Cluster Modulation Prevents Doxorubicin-Mediated Atrophy and Cardiotoxicity. Mol Ther 27:17–28 doi:10.1016/j.ymthe.2018.11.004

13. Hentze MW, Castello A, Schwarzl T, Preiss T (2018) A brave new world of RNA-binding proteins. Nat Rev Mol Cell Biol 19:327–341 doi:10.1038/nrm.2017.130

14. Katz Y, Li F, Lambert NJ, Sokol ES, Tam WL, Cheng AW, Airoldi EM, Lengner CJ, Gupta PB, Yu Z, Jaenisch R, Burge CB (2014) Musashi proteins are post-transcriptional regulators of the epithelial-luminal cell state. Elife 3:e03915 doi:10.7554/eLife.03915

15. Kharas MG, Lengner CJ (2017) Stem Cells, Cancer, and MUSASHI in Blood and Guts. Trends Cancer 3:347–356 doi:10.1016/j.trecan.2017.03.007

16. Kim D, Paggi JM, Park C, Bennett C, Salzberg SL (2019) Graph-based genome alignment and genotyping with HISAT2 and HISAT-genotype. Nat Biotechnol 37:907–915 doi:10.1038/s41587-019-0201-4

17. Kimura T, Ferran B, Tsukahara Y, Shang Q, Desai S, Fedoce A, Pimentel DR, Luptak I, Adachi T, Ido Y, Matsui R, Bachschmid MM (2019) Production of adeno-associated virus vectors for in vitro and in vivo applications. Sci Rep 9:13601 doi:10.1038/s41598-019-49624-w

18. Li M, Li AQ, Zhou SL, Lv H, Wei P, Yang WT (2020) RNA-binding protein MSI2 isoforms expression and regulation in progression of triple-negative breast cancer. J Exp Clin Cancer Res 39:92 doi:10.1186/s13046-020-01587-x

19. Nakamura M, Okano H, Blendy JA, Montell C (1994) Musashi, a neural RNA-binding protein required for Drosophila adult external sensory organ development. Neuron 13:67–81 doi:10.1016/0896-6273(94)90460-x

20. Oka SI, Sabry AD, Horiuchi AK, Cawley KM, O’Very SA, Zaitsev MA, Shankar TS, Byun J, Mukai R, Xu X, Torres NS, Kumar A, Yazawa M, Ling J, Taleb I, Saijoh Y, Drakos SG, Sadoshima J, Warren JS (2020) Perm1 regulates cardiac energetics as a downstream target of the histone methyltransferase Smyd1. PLoS One 15:e0234913 doi:10.1371/journal.pone.0234913

21. Pertea M, Pertea GM, Antonescu CM, Chang TC, Mendell JT, Salzberg SL (2015) StringTie enables improved reconstruction of a transcriptome from RNA-seq reads. Nat Biotechnol 33:290–295 doi:10.1038/nbt.3122

22. Riechert E, Kmietczyk V, Stein F, Schwarzl T, Sekaran T, Jurgensen L, Kamuf-Schenk V, Varma E, Hofmann C, Rettel M, Gur K, Olschlager J, Kuhl F, Martin J, Ramirez-Pedraza M, Fernandez M, Doroudgar S, Mendez R, Katus HA, Hentze MW, Volkers M (2021) Identification of dynamic RNA-binding proteins uncovers a Cpeb4-controlled regulatory cascade during pathological cell growth of cardiomyocytes. Cell Rep 35:109100 doi:10.1016/j.celrep.2021.109100

23. Roth GA, Mensah GA, Johnson CO, Addolorato G, Ammirati E, Baddour LM, Barengo NC, Beaton AZ, Benjamin EJ, Benziger CP, Bonny A, Brauer M, Brodmann M, Cahill TJ, Carapetis J, Catapano AL, Chugh SS, Cooper LT, Coresh J, Criqui M, DeCleene N, Eagle KA, Emmons-Bell S, Feigin VL, Fernandez-Sola J, Fowkes G, Gakidou E, Grundy SM, He FJ, Howard G, Hu F, Inker L, Karthikeyan G, Kassebaum N, Koroshetz W, Lavie C, Lloyd-Jones D, Lu HS, Mirijello A, Temesgen AM, Mokdad A, Moran AE, Muntner P, Narula J, Neal B, Ntsekhe M, Moraes de Oliveira G, Otto C, Owolabi M, Pratt M, Rajagopalan S, Reitsma M, Ribeiro ALP, Rigotti N, Rodgers A, Sable C, Shakil S, Sliwa-Hahnle K, Stark B, Sundstrom J, Timpel P, Tleyjeh IM, Valgimigli M, Vos T, Whelton PK, Yacoub M, Zuhlke L, Murray C, Fuster V, Group G-N-JGBoCDW (2020) Global Burden of Cardiovascular Diseases and Risk Factors, 1990-2019: Update From the GBD 2019 Study. J Am Coll Cardiol 76:2982–3021 doi:10.1016/j.jacc.2020.11.010

24. Sabater-Arcis M, Bargiela A, Moreno N, Poyatos-Garcia J, Vilchez JJ, Artero R (2021) Musashi-2 contributes to myotonic dystrophy muscle dysfunction by promoting excessive autophagy through miR-7 biogenesis repression. Mol Ther Nucleic Acids 25:652–667 doi:10.1016/j.omtn.2021.08.010

25. Sabbah HN (2020) Targeting the Mitochondria in Heart Failure: A Translational Perspective. JACC Basic Transl Sci 5:88–106 doi:10.1016/j.jacbts.2019.07.009

26. Sakakibara S, Imai T, Hamaguchi K, Okabe M, Aruga J, Nakajima K, Yasutomi D, Nagata T, Kurihara Y, Uesugi S, Miyata T, Ogawa M, Mikoshiba K, Okano H (1996) Mouse-Musashi-1, a neural RNA-binding protein highly enriched in the mammalian CNS stem cell. Dev Biol 176:230–242 doi:10.1006/dbio.1996.0130

27. Sakakibara S, Nakamura Y, Satoh H, Okano H (2001) Rna-binding protein Musashi2: developmentally regulated expression in neural precursor cells and subpopulations of neurons in mammalian CNS. J Neurosci 21:8091–8107

28. Scarpulla RC (2012) Nucleus-encoded regulators of mitochondrial function: integration of respiratory chain expression, nutrient sensing and metabolic stress. Biochim Biophys Acta 1819:1088–1097 doi:10.1016/j.bbagrm.2011.10.011

29. Schatton D, Pla-Martin D, Marx MC, Hansen H, Mourier A, Nemazanyy I, Pessia A, Zentis P, Corona T, Kondylis V, Barth E, Schauss AC, Velagapudi V, Rugarli EI (2017) CLUH regulates mitochondrial metabolism by controlling translation and decay of target mRNAs. J Cell Biol 216:675–693 doi:10.1083/jcb.201607019

30. Schoenberg DR, Maquat LE (2012) Regulation of cytoplasmic mRNA decay. Nat Rev Genet 13:246–259 doi:10.1038/nrg3160

31. Shi R, Ying S, Li Y, Zhu L, Wang X, Jin H (2021) Linking the YTH domain to cancer: the importance of YTH family proteins in epigenetics. Cell Death Dis 12:346 doi:10.1038/s41419-021-03625-8

32. Suresh Babu S, Joladarashi D, Jeyabal P, Thandavarayan RA, Krishnamurthy P (2015) RNA-stabilizing proteins as molecular targets in cardiovascular pathologies. Trends Cardiovasc Med 25:676–683 doi:10.1016/j.tcm.2015.02.006

33. Warren JS, Tracy CM, Miller MR, Makaju A, Szulik MW, Oka SI, Yuzyuk TN, Cox JE, Kumar A, Lozier BK, Wang L, Llana JG, Sabry AD, Cawley KM, Barton DW, Han YH, Boudina S, Fiehn O, Tucker HO, Zaitsev AV, Franklin S (2018) Histone methyltransferase Smyd1 regulates mitochondrial energetics in the heart. Proc Natl Acad Sci U S A 115:E7871–E7880 doi:10.1073/pnas.1800680115

34. Yang W, Yang L, Wang J, Zhang Y, Li S, Yin Q, Suo J, Ma R, Ye Y, Cheng H, Li J, Hui J, Hu P (2022) Msi2-mediated MiR7a-1 processing repression promotes myogenesis. J Cachexia Sarcopenia Muscle 13:728–742 doi:10.1002/jcsm.12882

35. Zhang Y, Si Y, Ma N, Mei J (2015) The RNA-binding protein PCBP2 inhibits Ang II-induced hypertrophy of cardiomyocytes though promoting GPR56 mRNA degeneration. Biochem Biophys Res Commun 464:679–684 doi:10.1016/j.bbrc.2015.06.139

36. Zhou A, Shi G, Kang GJ, Xie A, Liu H, Jiang N, Liu M, Jeong EM, Dudley SC, Jr. (2018) RNA Binding Protein, HuR, Regulates SCN5A Expression Through Stabilizing MEF2C transcription factor mRNA. J Am Heart Assoc 7 doi:10.1161/JAHA.117.007802

