## Supplementary Material for "Musashi-2 causes cardiac hypertrophy and heart failure by inducing mitochondrial dysfunction through destabilizing *Cluh* and *Smyd1* mRNA"

### Supplementary Fig. 1

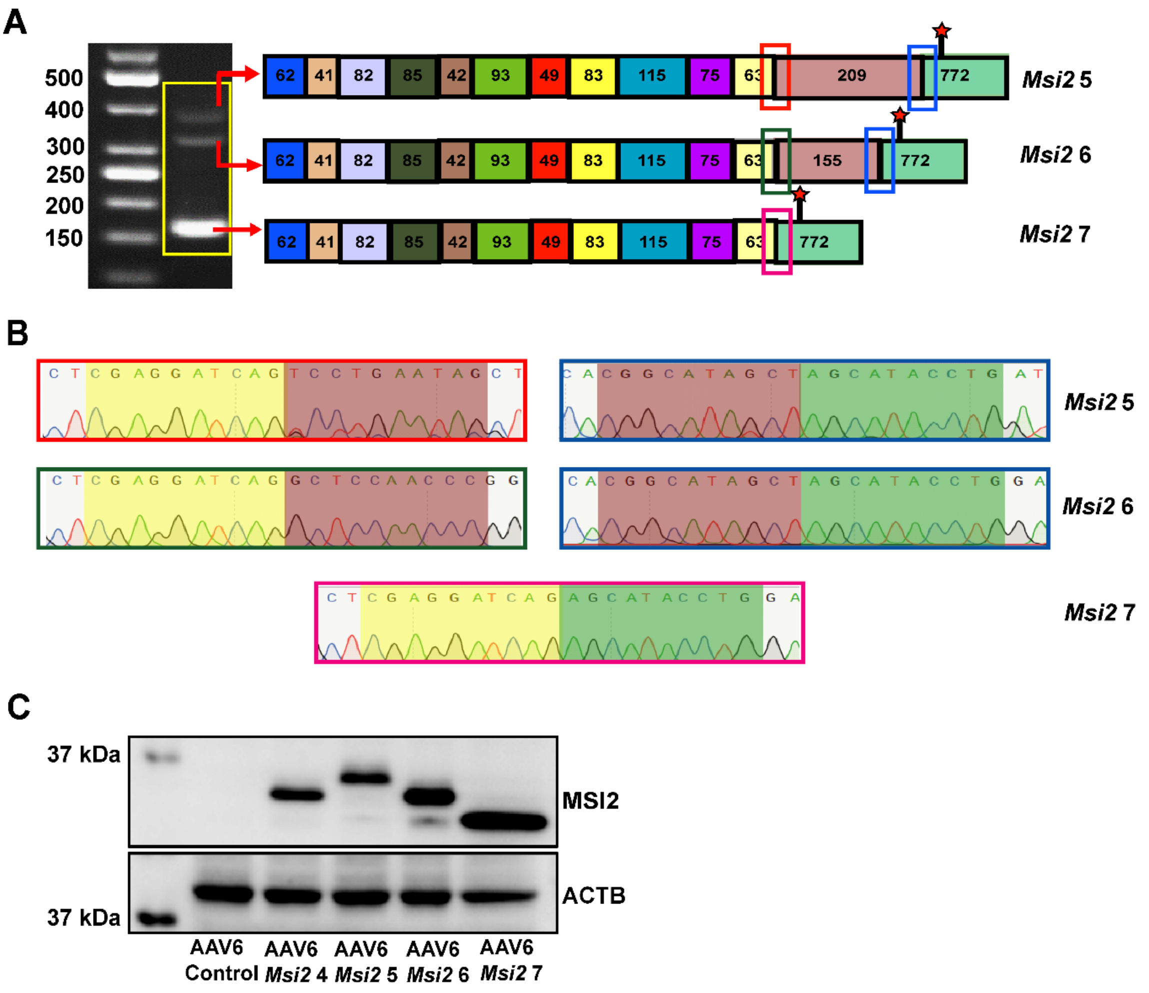


**Supplementary Fig. 1. (A)** *Msi2* isoforms amplified by PCR from the murine heart. (**B**) Sequencing reads depicting the unique exon junctions (highlighted by the box in figure **1B**) of each isoform of *Msi2*. (**C**) Western blot showing MSI2 levels in primary neonatal rat cardiomyocytes transduced with AAV6 encoding different isoforms of *Msi2* and GFP as control.

### Supplementary Fig. 2

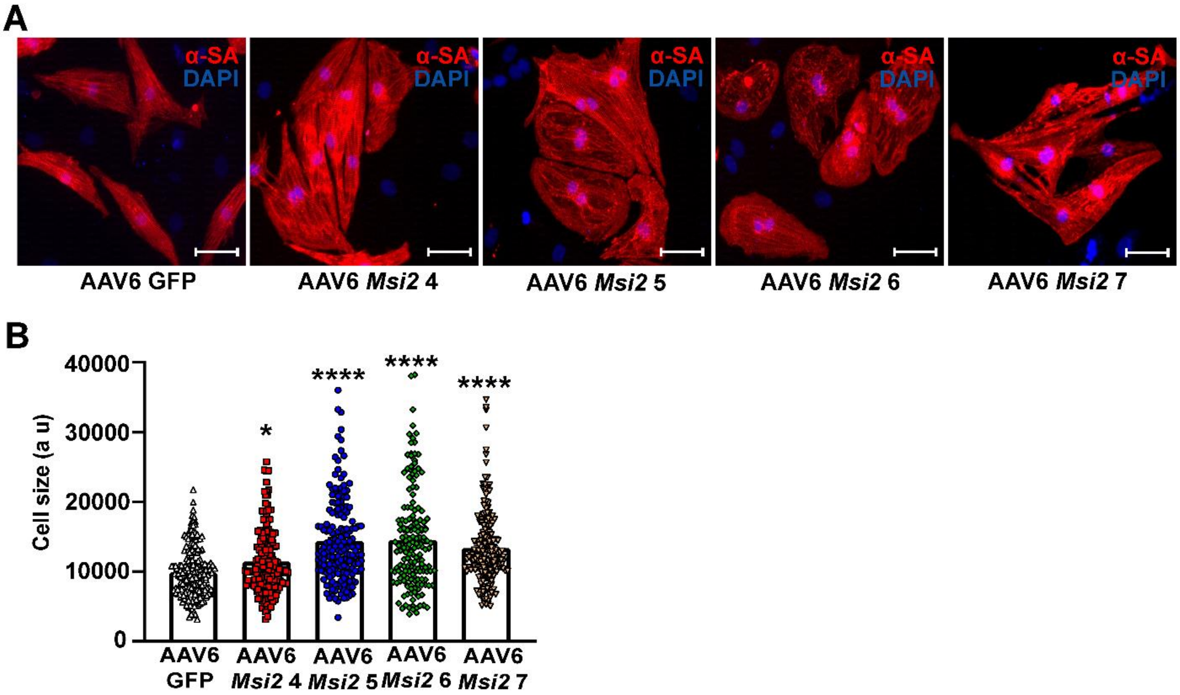


**Supplementary Fig. 2**. (**A**, **B**) Neonatal rat cardiomyocytes cell size after transduction with AAV6-*Msi2* isoforms and GFP control at MOI $10^4$. *$p \leq 0.05$, ****$p \leq 0.0001$

### Supplementary Fig. 3

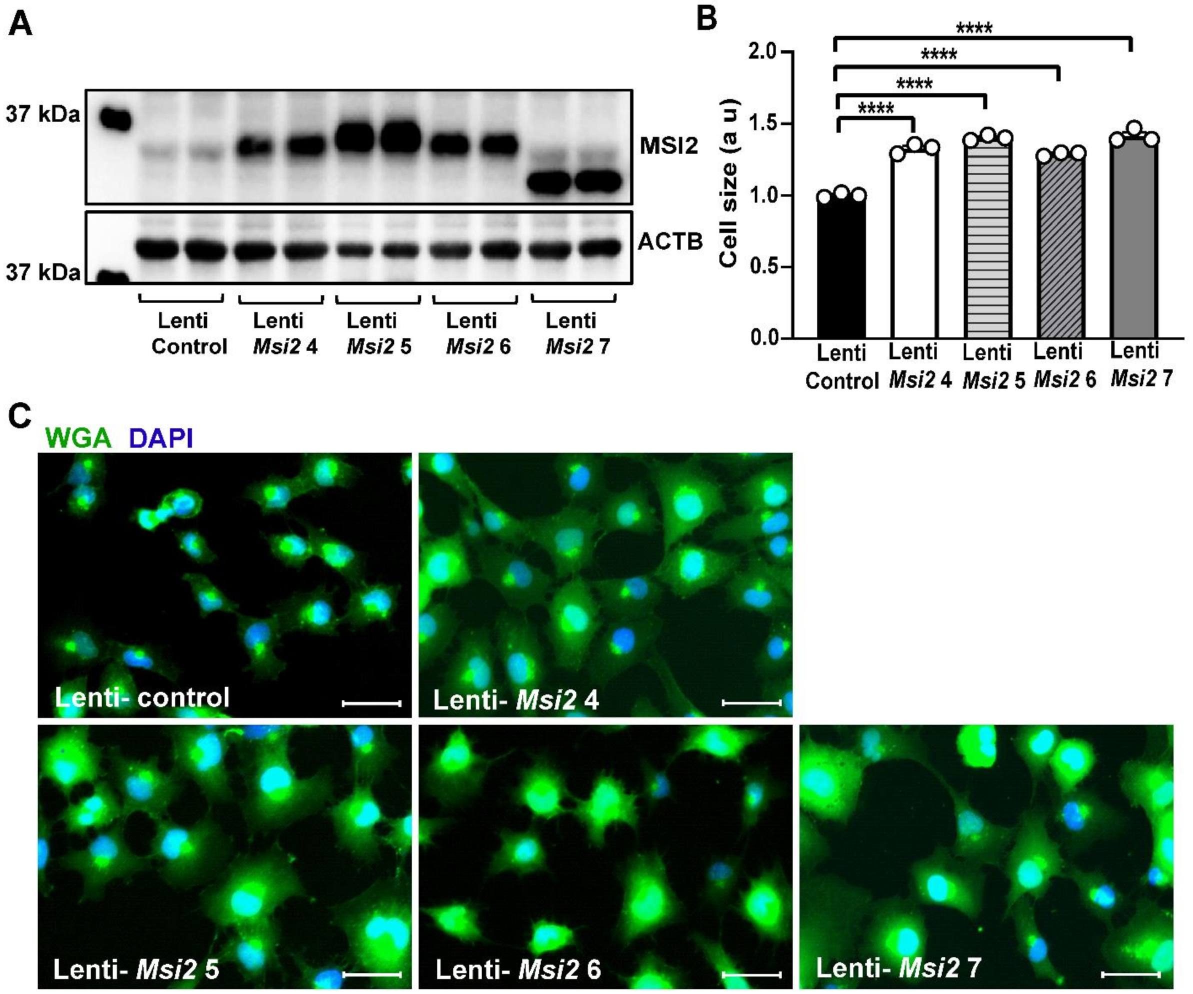


**Supplementary Fig. 3**. Human cardiomyocyte cell line AC16 cells stably overexpressing different isoforms of MSI2 and control (**A**), and their cell size (**B, C**) (n=3). The scale bar represents 50µm. ****p≤0.0001

### Supplementary Fig. 4

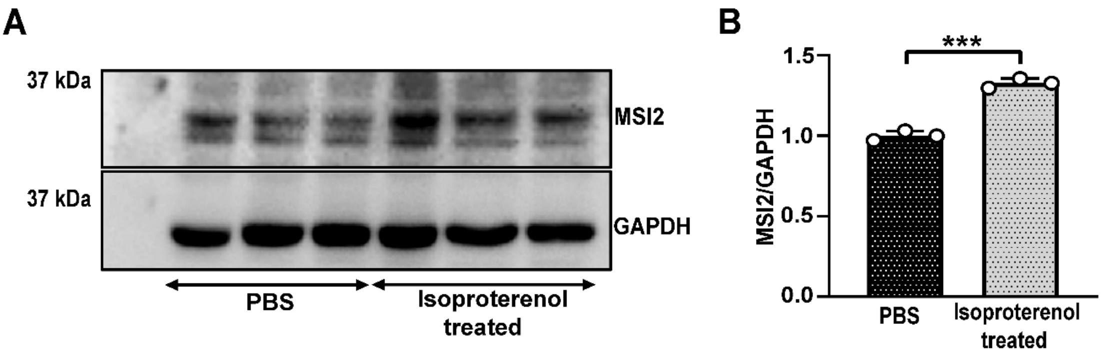


**Supplementary Fig. 4**. (**A**, **B**) MSI2 levels in neonatal rat cardiomyocytes after treatment with 20µM isoproterenol for seventy-two hours (n=3) ***p≤0.001

### Supplementary Fig. 5

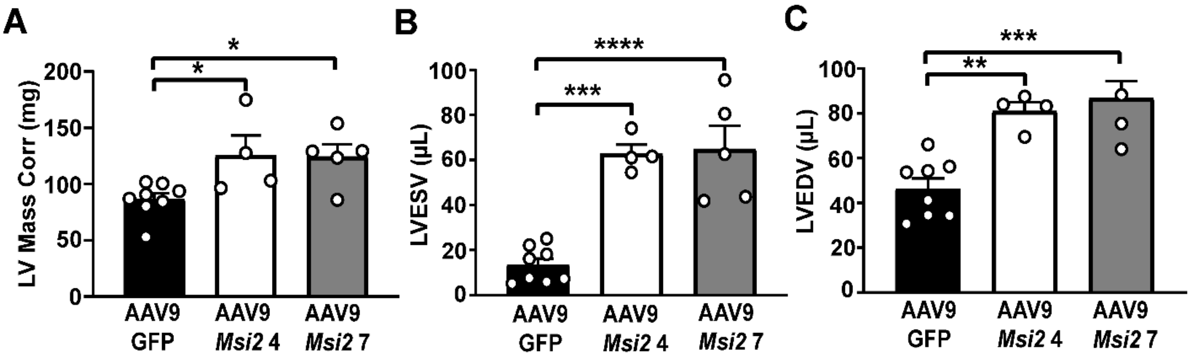


**Supplementary Fig. 5**. Echocardiographic data showing the cardiac function of *Msi2* overexpressing hearts compared to GFP control. LV- Left ventricular, LVESV – left ventricular end-systolic volume, LVEDD – left ventricular end-diastolic volume. *p≤0.05, **p≤0.01, ***p≤0.001, ****p≤0.0001

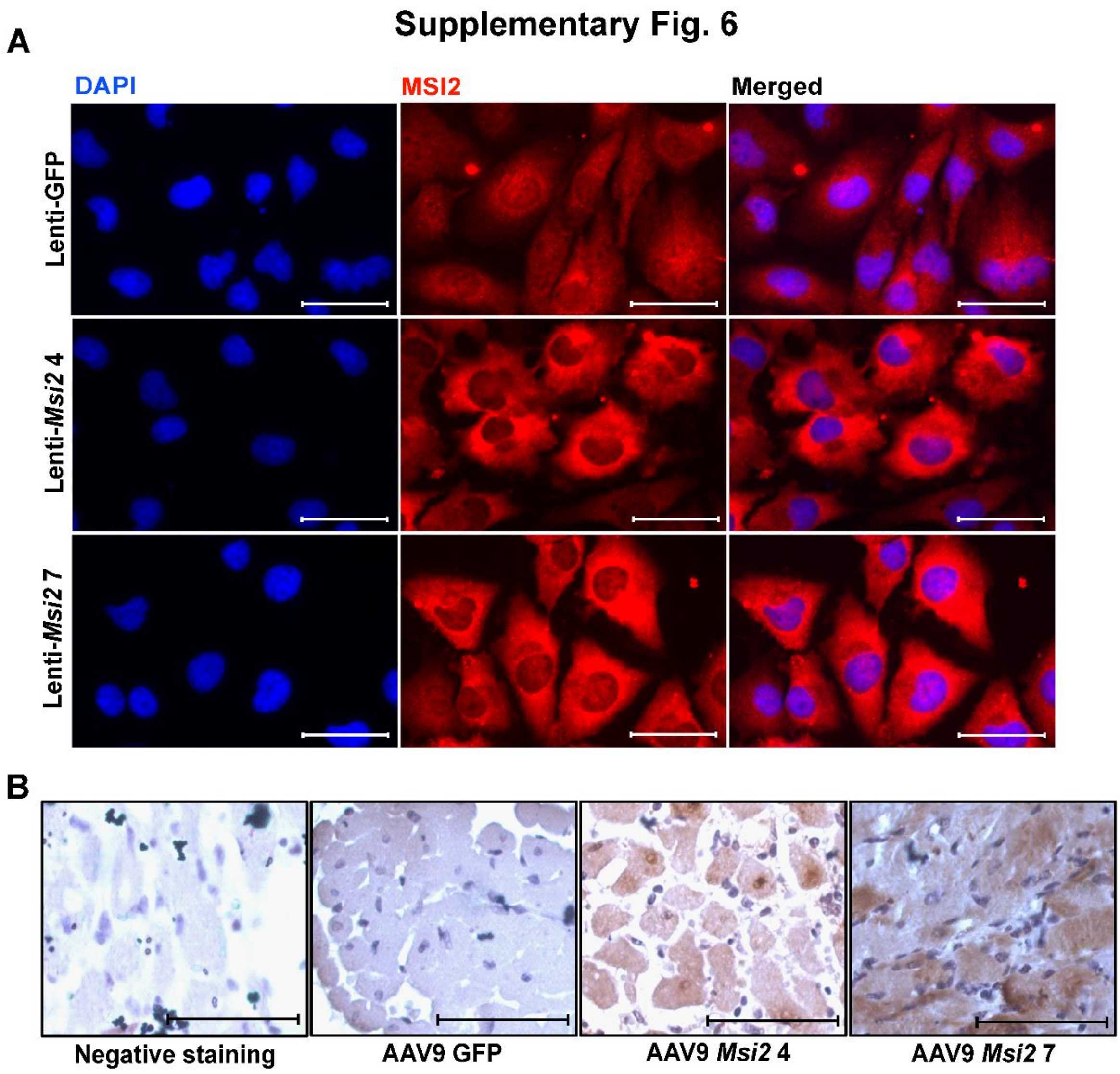


**Supplementary Fig. 6. Localization of MSI2**. (**A**) Immunofluoresence staining of MSI2 in AC16 cell line. (**B**) Immunohistochemistry staining of MSI2 in murine heart sections. The scale bar represents 50μm.

### Supplementary Fig. 7

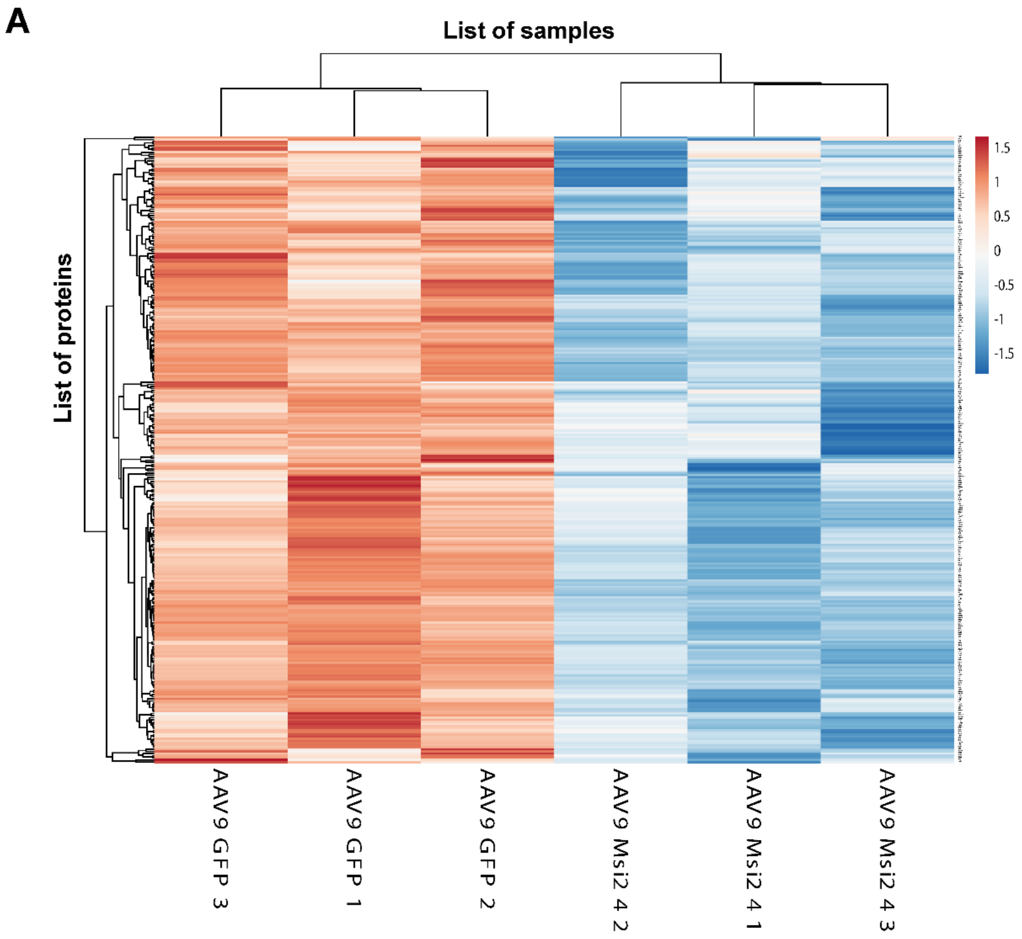


**Supplementary Fig. 7**. Heatmap showing expression pattern of nuclear-encoded mitochondrial genes in AAV9-*Msi2* 4 or GFP control transduced heart from proteomics analysis.

**Supplementary Table 1.** Primer list for Real-time PCR

| **Primer** | | **5' Sequence 3'** |
|---|---|---|
| *Msi1* | Fwd | CGGGGAGGTGAAAGAGTGTC |
| | Rev | TTCGAGGAAAGGCCACCTTG |
| *Msi2* | Fwd | CTACCCCAACTTTGTGGCAAC |
| | Rev | GCTGCCACTGGTCCATAAG |
| *Msi2 Iso1* | Fwd | TCGAGGATCAGTCCTGAATAGC |
| | Rev | TCAGTGGTATCCATTTGTAAAG |
| *Msi2 Iso4* | Fwd | CGAGGATCAGGCTCCAACCC |
| | Rev | TCAGTGGTATCCATTTGTAAAG |
| *Msi2 Iso5* | Fwd | TCGAGGATCAGTCCTGAATAGC |
| | Rev | CCAGGTATGCTAGCTATGCCG |
| *Msi2 Iso6* | Fwd | CGAGGATCAGGCTCCAACCC |
| | Rev | CCAGGTATGCTAGCTATGCCG |
| *Msi2 Iso7* | Fwd | CTACCCCAACTTTGTGGCAAC |
| | Rev | CCAGGTATGCTCTGATCCTCG |
| *Msi2 Variants* | Fwd | CTACCCCAACTTTGTGGCAAC |
| | Rev | GCCTGGACATCCAGGTATGC |
| *Mmu HPRT* | Fwd | GCGTCGTGATTAGCGATGAT |
| | Rev | TCCTTCATGACATCTCGAGCA |
| *Rat HPRT* | Fwd | CAGTCAACGGGGGACATAA |
| | Rev | GCTGTACTGCTTGACCAAGG |
| *Nppa* | Fwd | CCTGTGTACAGTGCGGTGTC |
| | Rev | CCTAGAAGCACTGCCGTCTC |
| *Nppb* | Fwd | CTGAAGGTGCTGTCCAGAT |
| | Rev | GTTCTTTTGTGAGGCCTTGG |
| *Myh6* | Fwd | GGTCCACATTCTTCAGGATTCTC |
| | Rev | GCGTTCCTTCTCTGACTTTCG |
| *Myh7* | Fwd | TCTCCTGCTGTTTCCTTACTTGCT |
| | Rev | CAGGCCTGTAGAAGAGCTGTACTC |

| | | |
|---|---|---|
| *Cluh* | Fwd | TCAGAGCTACGCAGTGTGGA |
| | Rev | AAGGCATCAGATGGGTCCAAG |
| *Hadha* | Fwd | AAATCTGGGCCAACGACCAAA |
| | Rev | CTTCCTGTGATATTCGTGTTGCT |
| *Mccc1* | Fwd | ACCATGAAGTATGGAACAACCC |
| | Rev | TGCACACCCATCTTTTTGGCT |
| *Pdha1* | Fwd | ACCACTCCTTTTGAGTGCCT |
| | Rev | AGCTGATCCGCCTTTAGCTC |
| *Acat1* | Fwd | TGTAAAAGACGGGCTAACTGATG |
| | Rev | TGTTCCTGCCGTGAGATATTCAT |
| *Pcca* | Fwd | GGTTTTAGGGGATAAACATGGCA |
| | Rev | CCATTGCTTGGCGAGTTTCA |
| *Smyd1* | Fwd | ATAGACAAGGCTCGCTCCGA |
| | Rev | CCATCAACCATCCTCCTGGC |
| *Ppargc1a* | Fwd | AGTGGTGTAGCGACCAATCG |
| | Rev | GTGTCTCTGTGAGAACCGCT |
| *Perm1* | Fwd | CTGAGGAAGGATTTGGGCACA |
| | Rev | CAGAGATGGTGCCAATGTTGG |
| *Cluh_3'UTR_1* | Fwd | GATGGACAGTCCAGCAGCCC |
| | Rev | AGGAAGGACGGAGCAAGACATC |
| *Cluh_3'UTR_2* | Fwd | CCTGATGTACATAGCCCAGTCAG |
| | Rev | GGAGGCAAGTCAAAGCCTGT |
| *Cluh_3'UTR_3* | Fwd | TGCTCCCTGTACCCCAGGTG |
| | Rev | GGCAGTAGGAGCCCCAACAC |
| *Cluh_3'UTR_4* | Fwd | GTGTTGGGGCTCCTACTGCC |
| | Rev | GCAGGTTTCAAAAACTCACTTTATTCC |
| *Smyd1_3'UTR_1* | Fwd | AGCAGTGAGGACCTTGTTGG |
| | Rev | AGCCCCAGAGATGCCTACAC |
| *Smyd1_3'UTR_2* | Fwd | CTGCTGGTCTCATCTACAGTTATG |
| | Rev | GTCTTGGAAGCCCCTACACAC |
| *Smyd1_3'UTR_3* | Fwd | TGAGAAGTCTGCCTCAGCCTAG |

| | | |
|---|---|---|
| | Rev | AGAGTTCTGGGAGATGTCAGTG |
| *Smyd1_3'UTR_4* | Fwd | AACATCTTGAAGTCACATGATTCCC |
| | Rev | GGCTTGTTTACTGTGGCTCTC |
| *Smyd1_3'UTR_5* | Fwd | TAAGAGTTCCACGCACCCCC |
| | Rev | TCCGAGAGCATCCAAGGGAAT |

**Supplementary Table 2.** FPKM counts for different *Msi2* variants expressed in cardiomyocytes

| Transcript IDs | Cardiomyocyte 1 | Cardiomyocyte 2 | Cardiomyocyte 3 |
|---|---|---|---|
| XM_011249302.3 | 0 | 0 | 0 |
| XM_036157065.1 | 0 | 0.102998 | 0.128234 |
| XM_006534440.4 | 0 | 0.266345 | 0.17102 |
| XM_006534435.4 | 0 | 0.336725 | 0.111734 |
| XM_036157064.1 | 0 | 0 | 0 |
| XM_006534438.4 | 0 | 0 | 0 |
| XM_006534439.4 | 0 | 0 | 0 |
| XM_006534431.4 | 0 | 0 | 0 |
| XM_006534432.4 | 0 | 0 | 0 |
| XM_006534441.5 | 0 | 0 | 0 |
| XM_011249305.4 | 0 | 0 | 0 |
| XM_030246417.1 | 0 | 0.106979 | 1.360224 |
| XM_036157066.1 | 0 | 0 | 0 |
| XM_036157067.1 | 0 | 0.19272 | 0.462448 |
| XM_036157062.1 | 0 | 0.157003 | 0.203143 |
| XM_006534434.4 | 0 | 0.070402 | 0.099925 |
| XM_011249304.4 | 0 | 0 | 0 |
| XM_030246418.2 | 0 | 0.003464 | 0 |
| XM_036157063.1 | 1.103833 | 1.282824 | 3.810156 |
| XM_036157068.1 | 1.237758 | 1.401667 | 4.299333 |
| NM_001201341.1 | 0 | 0 | 0 |
| NM_001363194.1 | 0 | 0 | 0 |
| NM_001363195.1 | 4.264986 | 2.330235 | 1.516321 |
| NM_001373923.1 | 5.203801 | 11.656748 | 4.54784 |
| NM_054043.3 | 3.853275 | 1.931582 | 1.966839 |

**Supplementary Table 3.** Average exon coverage of *Msi2* isoforms

| **Transcript ID** | **Cardiomyocyte 1** | **Cardiomyocyte 2** | **Cardiomyocyte 3** |
|---|---|---|---|
| NM_054043.3 | 2.456958357 | 3.622366143 | 2.427603 |
| NM_001363195.1 | 2.468968429 | 4.225165643 | 5.627823071 |
| NM_001373923.1 | 4.996078462 | 22.46611508 | 6.578222923 |
| XM_036157063.1 | 17.60338231 | 18.38252423 | 21.20132992 |
| XM_036157068.1 | 41.59262067 | 24.83498092 | 43.1145585 |

**Supplementary Table 4.** List of differentially expressed proteins in AAV9 *Msi2* 4 hearts compared to AAV9 GFP control

| SN | Upregulated Proteins | logFC | adj.P.Val |
|---|---|---|---|
| 1 | **Fcgr2** | 5.838026768 | 0.00193293 |
| 2 | **Cotl1** | 5.621166257 | 0.001227373 |
| 3 | **Col3a1** | 4.987707929 | 0.002384229 |
| 4 | **Col1a2** | 4.980211858 | 0.000892995 |
| 5 | **Emilin1** | 4.90616053 | 0.001789432 |
| 6 | **Gfpt2** | 4.876932873 | 0.001075829 |
| 7 | **Postn** | 4.840527513 | 0.000892995 |
| 8 | **Xirp2** | 4.817846909 | 0.001227373 |
| 9 | **Csrp2** | 4.710474735 | 0.001715592 |
| 10 | **Tnc** | 4.469737768 | 0.000892995 |
| 11 | **Aurkaip1** | 4.285271801 | 0.002355372 |
| 12 | **Capg** | 4.282222773 | 0.000892995 |
| 13 | **Tubb6** | 4.218435659 | 0.006095987 |
| 14 | **Hspa1b** | 4.210237499 | 0.000892995 |
| 15 | **Plbd1** | 4.14476941 | 0.001227373 |
| 16 | **Lcp1** | 4.125499349 | 0.001227373 |
| 17 | **Msi2** | 4.071410336 | 0.000892995 |
| 18 | **Plod1** | 3.972163066 | 0.025764045 |
| 19 | **Lgals3** | 3.916136225 | 0.001411055 |
| 20 | **H2-Ab1** | 3.852155854 | 0.005389611 |
| 21 | **Tmem43** | 3.823723528 | 0.001789432 |
| 22 | **Hexb** | 3.784985518 | 0.004689724 |
| 23 | **Mta2** | 3.778099765 | 0.002258852 |
| 24 | **Actg1** | 3.688284415 | 0.044165162 |
| 25 | **Ass1** | 3.660776113 | 0.001227373 |
| 26 | **Smc1a** | 3.608436946 | 0.001876125 |
| 27 | **Ltf** | 3.603802498 | 0.005389611 |
| 28 | **Basp1** | 3.580610792 | 0.001784651 |
| 29 | **Asah1** | 3.5799797 | 0.005324987 |
| 30 | **Naga** | 3.564156017 | 0.001227373 |
| 31 | **Tgtp2** | 3.532066714 | 0.001652328 |
| 32 | **Hpcal1** | 3.484304456 | 0.005177803 |
| 33 | **Ppic** | 3.474388143 | 0.001411055 |
| 34 | **S100a4** | 3.473662041 | 0.002108928 |
| 35 | **S100a9** | 3.452059272 | 0.001723939 |
| 36 | **Sgpl1** | 3.4262423 | 0.003575683 |
| 37 | **Coro1a** | 3.421162838 | 0.000892995 |
| 38 | **Ssr1** | 3.414119485 | 0.00345049 |
| 39 | **Elmo1** | 3.411478761 | 0.007272328 |
| 40 | **Rhog** | 3.403026564 | 0.001411055 |
| 41 | **Atp6v1h** | 3.401331656 | 0.021820028 |
| 42 | **Sf3a1** | 3.394365846 | 0.001227373 |

| SN | Downregulated Proteins | logFC | adj.P.Val |
|---|---|---|---|
| 1 | **Cox7a1** | -1.003027858 | 0.04203784 |
| 2 | **Gsta4** | -1.010024024 | 0.019826301 |
| 3 | **Hmgcl** | -1.011620341 | 0.047077118 |
| 4 | **Naca** | -1.01199629 | 0.00628681 |
| 5 | **Eno2** | -1.013676712 | 0.011225625 |
| 6 | **Dock2** | -1.018120435 | 0.020099706 |
| 7 | **Fah** | -1.022760281 | 0.018356261 |
| 8 | **Atp5mk** | -1.024259245 | 0.006852624 |
| 9 | **Pygb** | -1.025749077 | 0.012235034 |
| 10 | **Adh5** | -1.028382046 | 0.019617695 |
| 11 | **Ethe1** | -1.029210971 | 0.008645795 |
| 12 | **Dnm1l** | -1.037928753 | 0.008641207 |
| 13 | **Mrpl12** | -1.039974123 | 0.035176836 |
| 14 | **Gatd3** | -1.041944798 | 0.011359408 |
| 15 | **Tigar** | -1.043278077 | 0.015245017 |
| 16 | **Sod1** | -1.044154988 | 0.006671056 |
| 17 | **Ppm1a** | -1.044881381 | 0.02270553 |
| 18 | **Fxn** | -1.04556941 | 0.021204314 |
| 19 | **Suclg2** | -1.047641553 | 0.007083373 |
| 20 | **Ndufb3** | -1.047822747 | 0.011612404 |
| 21 | **Mug1** | -1.050183266 | 0.013191152 |
| 22 | **Atad3** | -1.05539425 | 0.021501323 |
| 23 | **Cox5a** | -1.055420756 | 0.009323093 |
| 24 | **Samm50** | -1.056205982 | 0.012433221 |
| 25 | **Cox7c** | -1.060689778 | 0.020347515 |
| 26 | **Ndufb8** | -1.061994637 | 0.009142182 |
| 27 | **Csrp1** | -1.066759353 | 0.012867304 |
| 28 | **Alyref** | -1.067463747 | 0.01314478 |
| 29 | **Nmnat3** | -1.067816168 | 0.006096852 |
| 30 | **Pdcd5** | -1.072346737 | 0.024742012 |
| 31 | **Trap1** | -1.074216713 | 0.004933539 |
| 32 | **Prdx5** | -1.077199117 | 0.007530902 |
| 33 | **Sgca** | -1.078208536 | 0.009595491 |
| 34 | **Gpc1** | -1.079379563 | 0.006310413 |
| 35 | **Fam162a** | -1.088100433 | 0.038620529 |
| 36 | **Mrps2** | -1.088746959 | 0.013686997 |
| 37 | **Glrx3** | -1.090278666 | 0.029877207 |
| 38 | **Ptges2** | -1.090708688 | 0.010599131 |
| 39 | **Smyd1** | -1.091337783 | 0.023463067 |
| 40 | **Palmd** | -1.097113503 | 0.012235034 |
| 41 | **Mpst** | -1.098920349 | 0.041103626 |
| 42 | **Nit2** | -1.101539433 | 0.043221493 |

| | | | |
|---|---|---|---|
| 43 | **Slc9a3r1** | 3.37961975 | 0.000892995 |
| 44 | **Rbp1** | 3.353259211 | 0.00091976 |
| 45 | **Klhl40** | 3.339738439 | 0.004500364 |
| 46 | **Ostf1** | 3.319779263 | 0.034198823 |
| 47 | **M6pr** | 3.285042442 | 0.005108636 |
| 48 | **Sh3bgrl3** | 3.268299824 | 0.002108928 |
| 49 | **Spcs3** | 3.253559843 | 0.001366147 |
| 50 | **Clpb** | 3.23485893 | 0.000928364 |
| 51 | **Dhx15** | 3.226628529 | 0.001715592 |
| 52 | **Psap** | 3.205435821 | 0.013772291 |
| 53 | **Serpinb1a** | 3.193130704 | 0.001227373 |
| 54 | **S100a11** | 3.18005215 | 0.015712596 |
| 55 | **Klc1** | 3.172782077 | 0.001227373 |
| 56 | **Col4a1** | 3.158916554 | 0.002355372 |
| 57 | **Ptprc** | 3.158312235 | 0.00229262 |
| 58 | **Tgfbi** | 3.096002441 | 0.000892995 |
| 59 | **Col14a1** | 3.064392595 | 0.002087105 |
| 60 | **Iigp1** | 3.054954788 | 0.000892995 |
| 61 | **Uap1l1** | 3.031897716 | 0.001419231 |
| 62 | **Gbp2** | 3.029415617 | 0.001227373 |
| 63 | **Rpl32** | 2.988043719 | 0.002926368 |
| 64 | **H1-1** | 2.972245881 | 0.001723939 |
| 65 | **Lipa** | 2.959705498 | 0.001227373 |
| 66 | **Fasn** | 2.959415395 | 0.001652328 |
| 67 | **H2-Aa** | 2.900505933 | 0.000892995 |
| 68 | **Clic1** | 2.844384123 | 0.000928364 |
| 69 | **C1qa** | 2.839239336 | 0.001974852 |
| 70 | **C1qb** | 2.836852236 | 0.002093653 |
| 71 | **Cstb** | 2.826421068 | 0.006484299 |
| 72 | **Col1a1** | 2.822820073 | 0.006444507 |
| 73 | **Snrpe** | 2.812053448 | 0.001789432 |
| 74 | **Ddx39a** | 2.792074048 | 0.001411055 |
| 75 | **Gbp5** | 2.78166821 | 0.001419231 |
| 76 | **Samhd1** | 2.771524342 | 0.004892855 |
| 77 | **Scamp2** | 2.741461089 | 0.004955233 |
| 78 | **Mtpn** | 2.741252627 | 0.00432025 |
| 79 | **Iqgap1** | 2.738582933 | 0.001227373 |
| 80 | **Trim28** | 2.738274336 | 0.015070922 |
| 81 | **Lyz1** | 2.718040939 | 0.001852133 |
| 82 | **Sf3a3** | 2.713061376 | 0.001227373 |
| 83 | **Mgst1** | 2.695366208 | 0.011696402 |
| 84 | **Eea1** | 2.673258397 | 0.009323728 |
| 85 | **Abhd12** | 2.669724531 | 0.001411055 |
| 86 | **Psmb10** | 2.667746562 | 0.004082264 |
| 87 | **Atp6v0c** | 2.648407454 | 0.038106543 |
| 88 | **Eif4a3** | 2.63592947 | 0.014114886 |
| 89 | **Nrap** | 2.633299246 | 0.001253158 |
| 90 | **Rrbp1** | 2.618439414 | 0.001366147 |

| | | | |
|---|---|---|---|
| 43 | **Ig lambda-1** | -1.104065846 | 0.042369332 |
| 44 | **Gfm2** | -1.104308375 | 0.035176836 |
| 45 | **Phgdh** | -1.106168776 | 0.014458496 |
| 46 | **Macrod1** | -1.107886052 | 0.006867105 |
| 47 | **Hspa4l** | -1.108950901 | 0.005970115 |
| 48 | **Myh14** | -1.114107264 | 0.018356261 |
| 49 | **Unc45b** | -1.118025902 | 0.00469793 |
| 50 | **Parvb** | -1.121959285 | 0.04964057 |
| 51 | **Isoc1 1** | -1.128124046 | 0.014139473 |
| 52 | **Mix23** | -1.132962855 | 0.023244473 |
| 53 | **Sorbs1** | -1.141327908 | 0.004730759 |
| 54 | **Mrps18a** | -1.145668587 | 0.030518216 |
| 55 | **Cavin4** | -1.146676699 | 0.006310413 |
| 56 | **Glo1** | -1.147353505 | 0.022918342 |
| 57 | **Adipoq** | -1.148083435 | 0.015234655 |
| 58 | **Ndufb11** | -1.152325147 | 0.019587136 |
| 59 | **Hsd17b10** | -1.152852451 | 0.005752091 |
| 60 | **Tmem256** | -1.153874464 | 0.008508975 |
| 61 | **Sqor** | -1.15493905 | 0.016883653 |
| 62 | **Cycs** | -1.156560297 | 0.005694914 |
| 63 | **Gys1** | -1.157458368 | 0.020081846 |
| 64 | **Mgll** | -1.162101082 | 0.004272328 |
| 65 | **Cyb5r1** | -1.162788113 | 0.014139473 |
| 66 | **Mrpl49** | -1.163227317 | 0.02597907 |
| 67 | **Dnaja3** | -1.163581291 | 0.003157642 |
| 68 | **Oplah** | -1.168275117 | 0.013772291 |
| 69 | ***Got2*** | -1.169634581 | 0.006644354 |
| 70 | **Serpina1d** | -1.171695883 | 0.045078872 |
| 71 | **Pzp** | -1.173144545 | 0.015608905 |
| 72 | **Flad1** | -1.177591217 | 0.0120702 |
| 73 | **Sdhb** | -1.178915095 | 0.006231956 |
| 74 | **Atp1a1** | -1.18286747 | 0.003240174 |
| 75 | **Hibch 1** | -1.183442923 | 0.004069716 |
| 76 | **Krt2** | -1.18980253 | 0.01187943 |
| 77 | **Macf1** | -1.197754909 | 0.012235034 |
| 78 | **Krt20** | -1.199243586 | 0.005389611 |
| 79 | **Hspd1** | -1.20072134 | 0.008926329 |
| 80 | **Vdac3** | -1.203611101 | 0.004194692 |
| 81 | **Serpina1b** | -1.208353933 | 0.024206957 |
| 82 | **Dtna** | -1.208426948 | 0.004352985 |
| 83 | **Lonp1** | -1.212703382 | 0.020347515 |
| 84 | **Apip** | -1.214706308 | 0.013527194 |
| 85 | **Dhrs4** | -1.21786051 | 0.0119182 |
| 86 | **Surf1** | -1.220887499 | 0.015608905 |
| 87 | **Acad9** | -1.226296139 | 0.005797345 |
| 88 | **Grpel1** | -1.22929311 | 0.006199316 |
| 89 | **Serhl** | -1.229869171 | 0.034694498 |
| 90 | **Popdc2** | -1.232153211 | 0.021766598 |

| | | | |
|---|---|---|---|
| 91 | **Sf3b3** | 2.617140097 | 0.003198889 |
| 92 | **Fn1** | 2.595882412 | 0.001202217 |
| 93 | **Arf4** | 2.590135605 | 0.003157642 |
| 94 | **C4orf54** | 2.572360993 | 0.007024432 |
| 95 | **G6pdx** | 2.572258992 | 0.003198889 |
| 96 | **Pak2** | 2.568541928 | 0.003198889 |
| 97 | **PSec61g** | 2.566064314 | 0.032177193 |
| 98 | **Clu** | 2.560736273 | 0.001411055 |
| 99 | **Adpgk** | 2.554845504 | 0.001227373 |
| 100 | **Ptgis** | 2.549881115 | 0.037405375 |
| 101 | **Arhgdib** | 2.544515164 | 0.001419231 |
| 102 | **Ptbp1** | 2.537469363 | 0.001587411 |
| 103 | **H1-5** | 2.535880245 | 0.001336876 |
| 104 | **Eif6** | 2.531119143 | 0.001723939 |
| 105 | **Acly** | 2.528450051 | 0.007272328 |
| 106 | **Stat1** | 2.504301591 | 0.001913633 |
| 107 | **Wasf2** | 2.503869781 | 0.003615006 |
| 108 | **Camp** | 2.50358228 | 0.004596101 |
| 109 | **Arpc1b** | 2.501865001 | 0.001227373 |
| 110 | **Uggt1** | 2.501408216 | 0.006580593 |
| 111 | **Ube2d3** | 2.478520558 | 0.033995739 |
| 112 | **Ssr4** | 2.47241245 | 0.001784651 |
| 113 | **U2af1** | 2.469920277 | 0.003011038 |
| 114 | **Ckap4** | 2.462790705 | 0.000928364 |
| 115 | **Anxa1** | 2.45887718 | 0.001928588 |
| 116 | **Arpc5** | 2.456366903 | 0.001411055 |
| 117 | **Aldh1a2** | 2.45213817 | 0.001199851 |
| 118 | **Sfpq** | 2.446070704 | 0.001170994 |
| 119 | **Cyrib** | 2.432303148 | 0.000928364 |
| 120 | **Eef1a1** | 2.419653066 | 0.001784651 |
| 121 | **Coro1c** | 2.41601143 | 0.008052273 |
| 122 | **Gvin1** | 2.404903263 | 0.003587143 |
| 123 | **Kctd12** | 2.397996802 | 0.001227373 |
| 124 | **Mybbp1a** | 2.389050084 | 0.001411055 |
| 125 | **Erp29** | 2.387189407 | 0.001411055 |
| 126 | **Copz1** | 2.378675972 | 0.008576062 |
| 127 | **H2-K1** | 2.368459021 | 0.001411055 |
| 128 | **Lrrc59** | 2.359718091 | 0.00181256 |
| 129 | **Myof** | 2.355341218 | 0.01034688 |
| 130 | **Tsn** | 2.350874819 | 0.012954478 |
| 131 | **Thy1** | 2.344002248 | 0.001411055 |
| 132 | **Lyar** | 2.334524072 | 0.014644977 |
| 133 | **Stmn1** | 2.32005414 | 0.002578606 |
| 134 | **Pdxk** | 2.300101089 | 0.003438574 |
| 135 | **Dpysl3** | 2.294117441 | 0.001411055 |
| 136 | **Gpld1** | 2.28963812 | 0.001784651 |
| 137 | **Psmb9** | 2.250800303 | 0.016065684 |
| 138 | **Fnbp1** | 2.244487023 | 0.006593127 |

| | | | |
|---|---|---|---|
| 91 | **Dnaja2** | -1.233097061 | 0.038620529 |
| 92 | **Nebl** | -1.233340913 | 0.020512605 |
| 93 | **Afg3l2** | -1.234354993 | 0.010733171 |
| 94 | **Gstm5** | -1.234615518 | 0.021820028 |
| 95 | **Gadd45gip1** | -1.234806147 | 0.039762646 |
| 96 | **Dmac1** | -1.235542452 | 0.044229718 |
| 97 | **Sirt5** | -1.239985184 | 0.005709066 |
| 98 | **Hspa9** | -1.240101511 | 0.002355372 |
| 99 | **Sod2** | -1.240716985 | 0.002434267 |
| 100 | **Huwe1** | -1.241788498 | 0.010519525 |
| 101 | **Msrb2** | -1.247373693 | 0.012469793 |
| 102 | **Gfm1** | -1.251756257 | 0.009474818 |
| 103 | **Nlrx1** | -1.252188205 | 0.015204686 |
| 104 | **Pnpo** | -1.254070388 | 0.037298635 |
| 105 | **Ecpas** | -1.256772145 | 0.03749847 |
| 106 | **Ndufa12** | -1.258689409 | 0.002694088 |
| 107 | **Dglucy** | -1.260207701 | 0.007365926 |
| 108 | **Atp5pb** | -1.262604732 | 0.008926329 |
| 109 | **Hspa12b** | -1.265398585 | 0.009954278 |
| 110 | **Srl** | -1.266788946 | 0.007404874 |
| 111 | **Mrpl28** | -1.268828914 | 0.003294322 |
| 112 | **Myo1d** | -1.269259157 | 0.045535704 |
| 113 | **Alad** | -1.270916785 | 0.041653686 |
| 114 | **Ndufb9** | -1.27292169 | 0.002405932 |
| 115 | **Ppp1r12b** | -1.278144979 | 0.002506478 |
| 116 | **Cavin2** | -1.281571266 | 0.003723688 |
| 117 | **Obscn** | -1.28382311 | 0.003147872 |
| 118 | **Ndufb5** | -1.284494292 | 0.004585192 |
| 119 | **Pdpr** | -1.285672597 | 0.020765124 |
| 120 | **Dlst** | -1.291992635 | 0.02489373 |
| 121 | **Cast** | -1.292182348 | 0.012863589 |
| 122 | **Slc25a4** | -1.293031651 | 0.019513909 |
| 123 | **Gpd1l** | -1.294122664 | 0.006964891 |
| 124 | **Isca2** | -1.296357652 | 0.005304655 |
| 125 | **Hccs** | -1.29891215 | 0.015107416 |
| 126 | **Naxe** | -1.306441654 | 0.011813334 |
| 127 | **Uqcrc1** | -1.30765815 | 0.003438574 |
| 128 | **Pdlim5** | -1.307796953 | 0.014741591 |
| 129 | **Myh7b** | -1.308764225 | 0.013506933 |
| 130 | **Pfkfb2** | -1.309562336 | 0.020314865 |
| 131 | **Cbr1** | -1.313115134 | 0.002228601 |
| 132 | **Cisd1** | -1.314606972 | 0.007247386 |
| 133 | **Grsf1** | -1.315684418 | 0.014435403 |
| 134 | **Pdhx** | -1.317842856 | 0.007642505 |
| 135 | **Actn2** | -1.318947085 | 0.003244867 |
| 136 | **Mybpc3** | -1.319045862 | 0.002108928 |
| 137 | **Obsl1** | -1.31974921 | 0.012892768 |
| 138 | **Lpin1** | -1.319751003 | 0.018098773 |

| | | | |
|---|---|---|---|
| 139 | **Hnrnpf** | 2.222421071 | 0.00346669 |
| 140 | **Atp6v1a** | 2.215152565 | 0.001411055 |
| 141 | **Niban1** | 2.214589125 | 0.001411055 |
| 142 | **Naa15** | 2.207683101 | 0.0226528 |
| 143 | **Pdia4** | 2.206123849 | 0.001307391 |
| 144 | **Anp32e** | 2.201909452 | 0.001411055 |
| 145 | **Rpl3** | 2.192424338 | 0.001411055 |
| 146 | **Tubb3** | 2.190551107 | 0.004498873 |
| 147 | **Ube2o** | 2.181585685 | 0.00175894 |
| 148 | **Srrt** | 2.174012782 | 0.001227373 |
| 149 | **Pgm2** | 2.173986954 | 0.018438357 |
| 150 | **Ctsa** | 2.173696858 | 0.003198889 |
| 151 | **Uchl1** | 2.166425821 | 0.001852133 |
| 152 | **Bzw1** | 2.161962297 | 0.001411055 |
| 153 | **Snx5** | 2.157504107 | 0.002108928 |
| 154 | **Sparc** | 2.145470564 | 0.002058898 |
| 155 | **Bgn** | 2.14203545 | 0.001813703 |
| 156 | **Sart1** | 2.142005912 | 0.009110296 |
| 157 | **Ap1g1** | 2.138429182 | 0.008299011 |
| 158 | **Sub1** | 2.136467681 | 0.002228601 |
| 159 | **Glipr2** | 2.131935531 | 0.002228601 |
| 160 | **Stat3** | 2.13152364 | 0.001227373 |
| 161 | **Lama4** | 2.125787158 | 0.003607292 |
| 162 | **Yars1** | 2.121592733 | 0.027646256 |
| 163 | **Mvp** | 2.112505771 | 0.001227373 |
| 164 | **Anxa4** | 2.093840658 | 0.001411055 |
| 165 | **Cyfip1** | 2.093821547 | 0.023463067 |
| 166 | **Sdcbp** | 2.091441503 | 0.041037931 |
| 167 | **Ctsb** | 2.090574377 | 0.002061334 |
| 168 | **Septin11** | 2.086333549 | 0.00244431 |
| 169 | **Fbn1** | 2.083288437 | 0.005073924 |
| 170 | **Niban2** | 2.079473232 | 0.001411055 |
| 171 | **Lrp1** | 2.07844752 | 0.001462694 |
| 172 | **Rrad** | 2.071250314 | 0.001876125 |
| 173 | **Dnajc3** | 2.067793474 | 0.029976837 |
| 174 | **Serpina3n** | 2.067435731 | 0.001227373 |
| 175 | **Arpc3** | 2.06232008 | 0.00319564 |
| 176 | **Fscn1** | 2.059721266 | 0.006484299 |
| 177 | **Raly** | 2.050901698 | 0.002166624 |
| 178 | **Nop56** | 2.045038693 | 0.022899313 |
| 179 | **Clptm1** | 2.03531164 | 0.012075634 |
| 180 | **Cndp2** | 2.033503305 | 0.001789432 |
| 181 | **Khsrp** | 2.032020366 | 0.001813703 |
| 182 | **Nans** | 2.018420462 | 0.003536193 |
| 183 | **Tkt** | 2.017967559 | 0.001411055 |
| 184 | **Ddx17** | 2.00678603 | 0.007866668 |
| 185 | **Rcn1** | 2.003583377 | 0.001227373 |
| 186 | **Fhl2** | 2.000846829 | 0.003488926 |

| | | | |
|---|---|---|---|
| 139 | **Myom1** | -1.322519684 | 0.002703907 |
| 140 | **Hmga1** | -1.322808756 | 0.018356261 |
| 141 | **Xxylt1** | -1.326781426 | 0.049907926 |
| 142 | **Ephx2** | -1.327705591 | 0.004052234 |
| 143 | **Eci2** | -1.337787348 | 0.003726037 |
| 144 | **Adss1** | -1.338413463 | 0.012997067 |
| 145 | **Etfdh** | -1.340898391 | 0.005694914 |
| 146 | **Ndufa7** | -1.341259753 | 0.014458496 |
| 147 | **Maob** | -1.342834531 | 0.006410651 |
| 148 | **Ndufaf3** | -1.345926976 | 0.006593127 |
| 149 | **Atp5pd** | -1.34659577 | 0.005389611 |
| 150 | **Plin2** | -1.347626211 | 0.007639493 |
| 151 | **Mrpl37** | -1.359593577 | 0.004785257 |
| 152 | **Aldh5a1** | -1.359775464 | 0.005389611 |
| 153 | **Dmd** | -1.361139901 | 0.005073924 |
| 154 | **Ndufa6** | -1.364861595 | 0.002228601 |
| 155 | **Prxl2b** | -1.364930058 | 0.045535704 |
| 156 | **Sccpdh** | -1.368363726 | 0.009954278 |
| 157 | **Fabp3** | -1.372073931 | 0.019032196 |
| 158 | **Suclg1** | -1.376052713 | 0.006965342 |
| 159 | **Tpm1** | -1.379017189 | 0.009940133 |
| 160 | **Mrpl58** | -1.381865585 | 0.033416847 |
| 161 | **Got1** | -1.382206009 | 0.002773631 |
| 162 | **Ak1** | -1.386722965 | 0.002126664 |
| 163 | **Atp5me** | -1.387666268 | 0.006310413 |
| 164 | **Phb** | -1.391902256 | 0.005694914 |
| 165 | **Uqcrb** | -1.391937686 | 0.008323225 |
| 166 | **Uqcr10** | -1.394454106 | 0.014675013 |
| 167 | **Rtn4ip1** | -1.395556418 | 0.011852921 |
| 168 | **Acads** | -1.397096212 | 0.003146495 |
| 169 | **Tfam** | -1.397362345 | 0.003157642 |
| 170 | **Micos13** | -1.399387242 | 0.004716741 |
| 171 | **Prkaa2** | -1.406658786 | 0.039762646 |
| 172 | **Bcat2** | -1.406724931 | 0.003438574 |
| 173 | **Hhatl** | -1.407281332 | 0.002467198 |
| 174 | **Acyp2** | -1.408218772 | 0.002970429 |
| 175 | **Eno3** | -1.408355782 | 0.006880988 |
| 176 | **Ttn** | -1.409559781 | 0.001789432 |
| 177 | **Fech** | -1.409941609 | 0.003586226 |
| 178 | **Mdh1** | -1.410634781 | 0.002506478 |
| 179 | **Mlycd** | -1.411166254 | 0.002257246 |
| 180 | **Taco1** | -1.411255106 | 0.010332099 |
| 181 | **Zadh2** | -1.412374079 | 0.013738644 |
| 182 | **Upf2** | -1.413003146 | 0.029791981 |
| 183 | **Krt1** | -1.415765396 | 0.013263167 |
| 184 | **Hdhd2** | -1.416821372 | 0.027955251 |
| 185 | **Tsfm** | -1.418189313 | 0.020868901 |
| 186 | **Tango2** | -1.421585031 | 0.046756585 |

| | | | |
|---|---|---|---|
| 187 | **Upf1** | 1.99355291 | 0.037068755 |
| 188 | **Actr2** | 1.991317979 | 0.001411055 |
| 189 | **Atp2b1** | 1.990582786 | 0.001411055 |
| 190 | **Lamtor4** | 1.980805055 | 0.035689714 |
| 191 | **Epb41l2** | 1.976317979 | 0.002578606 |
| 192 | **Cap1** | 1.972410972 | 0.001227373 |
| 193 | **Plek** | 1.964460903 | 0.004312154 |
| 194 | **Srp54** | 1.963918231 | 0.036802917 |
| 195 | **Eftud2** | 1.954070201 | 0.00194433 |
| 196 | **Nrdc** | 1.943275788 | 0.034897143 |
| 197 | **Spcs2** | 1.942636328 | 0.002087105 |
| 198 | **Actr3** | 1.936875191 | 0.001411055 |
| 199 | **Vma21** | 1.928324867 | 0.031468343 |
| 200 | **Ewsr1** | 1.912917386 | 0.003091002 |
| 201 | **Hnrnpa0** | 1.911102735 | 0.003438574 |
| 202 | **Cbx3** | 1.901595141 | 0.007718755 |
| 203 | **Cfl1** | 1.895450237 | 0.001462694 |
| 204 | **Rbbp4** | 1.88280046 | 0.00215446 |
| 205 | **Ppie** | 1.872868394 | 0.022662261 |
| 206 | **Nucb1** | 1.86624839 | 0.016246197 |
| 207 | **Prkcsh** | 1.862574514 | 0.008052273 |
| 208 | **Nt5c2** | 1.855729885 | 0.001411055 |
| 209 | **Hp** | 1.847376406 | 0.005108636 |
| 210 | **Snd1** | 1.846309469 | 0.001411055 |
| 211 | **Atp6v1c1** | 1.837618535 | 0.020347515 |
| 212 | **P4hb** | 1.83118329 | 0.001411055 |
| 213 | **Stk24** | 1.83072382 | 0.004964058 |
| 214 | **Acp2** | 1.827693067 | 0.007039775 |
| 215 | **Mapk1** | 1.827466995 | 0.001723939 |
| 216 | **Man2b1** | 1.825509196 | 0.035469277 |
| 217 | **Hnrnpdl** | 1.825347875 | 0.003588533 |
| 218 | **S100a13** | 1.814234061 | 0.001723939 |
| 219 | **Xirp1** | 1.814004525 | 0.001723939 |
| 220 | **NARS1** | 1.803154168 | 0.001411055 |
| 221 | **Erp44** | 1.802420727 | 0.003146495 |
| 222 | **Atp6v0d1** | 1.799506851 | 0.003213299 |
| 223 | **Pitpna** | 1.799120573 | 0.00206644 |
| 224 | **Tars1** | 1.798751527 | 0.001913633 |
| 225 | **Bag1** | 1.798206875 | 0.002542629 |
| 226 | **Emd** | 1.795013687 | 0.006473309 |
| 227 | **Hspb1** | 1.784869626 | 0.001852133 |
| 228 | **Por** | 1.784770069 | 0.002840352 |
| 229 | **Dbnl** | 1.782914241 | 0.003168885 |
| 230 | **Mrc1** | 1.782032368 | 0.029259834 |
| 231 | **Pecam1** | 1.773262878 | 0.003146495 |
| 232 | **Rraga** | 1.770806736 | 0.002434267 |
| 233 | **Pdia3** | 1.770025071 | 0.001411055 |
| 234 | **Ccdc22** | 1.766440748 | 0.039792304 |

| | | | |
|---|---|---|---|
| 187 | **Romo1** | -1.422738954 | 0.01002268 |
| 188 | **Ndufs5** | -1.4241073 | 0.002940966 |
| 189 | **Cpt2** | -1.427318961 | 0.002108928 |
| 190 | **Speg** | -1.429418324 | 0.01229201 |
| 191 | **Chchd7** | -1.430794099 | 0.013138395 |
| 192 | **Mtnd5** | -1.430894833 | 0.043646507 |
| 193 | **Pdha1** | -1.437807 | 0.002582278 |
| 194 | **Ppp2r5a** | -1.44008623 | 0.005631229 |
| 195 | **Slc25a11** | -1.441298132 | 0.002384229 |
| 196 | **Cdh5** | -1.44575075 | 0.022052418 |
| 197 | **Fkbp3** | -1.446303871 | 0.00981529 |
| 198 | **Gstm2** | -1.446906963 | 0.015132793 |
| 199 | **Ndufc2** | -1.448932048 | 0.005631229 |
| 200 | **Bola1** | -1.450960281 | 0.022581868 |
| 201 | **Tmem14c** | -1.456170364 | 0.008737773 |
| 202 | **Timm50** | -1.458122649 | 0.002355372 |
| 203 | **Meioc** | -1.462931319 | 0.024388495 |
| 204 | **Slc25a20** | -1.467820215 | 0.010596045 |
| 205 | **Dbt** | -1.471626268 | 0.001723939 |
| 206 | **Psmb5** | -1.472447295 | 0.006113682 |
| 207 | **Mrps26** | -1.475018413 | 0.029259834 |
| 208 | **Nceh1** | -1.480719531 | 0.001937701 |
| 209 | **Phb2** | -1.481000704 | 0.00229412 |
| 210 | **Pccb** | -1.481675404 | 0.002932108 |
| 211 | **Coq5** | -1.486439018 | 0.02474173 |
| 212 | **Mlip** | -1.49258657 | 0.003571267 |
| 213 | **Immt** | -1.49446325 | 0.002703907 |
| 214 | **Nqo1** | -1.497223884 | 0.040900158 |
| 215 | **Mtx1** | -1.498582858 | 0.010287246 |
| 216 | **Pfkm** | -1.498785028 | 0.002257246 |
| 217 | **Mtres1** | -1.502113721 | 0.02464866 |
| 218 | **Ecsit** | -1.502133251 | 0.008737773 |
| 219 | **Decr1** | -1.503999753 | 0.001852133 |
| 220 | **Mtfr1l** | -1.504790685 | 0.00181256 |
| 221 | **Ndufs4** | -1.505206078 | 0.001691139 |
| 222 | **Ndufs7** | -1.513105506 | 0.008323225 |
| 223 | **Fundc2** | -1.514823476 | 0.008723164 |
| 224 | **Eci1** | -1.516833164 | 0.004005926 |
| 225 | **Acy3** | -1.522027527 | 0.029259834 |
| 226 | **Ca4** | -1.522881351 | 0.006931118 |
| 227 | **Gcdh** | -1.523727507 | 0.003157642 |
| 228 | **Opa3** | -1.524893595 | 0.006922571 |
| 229 | **Atpaf1** | -1.525565901 | 0.010205985 |
| 230 | **Dld** | -1.526316016 | 0.002087105 |
| 231 | **Acadvl** | -1.531366845 | 0.001903534 |
| 232 | **Atp5f1c** | -1.537099289 | 0.004312154 |
| 233 | **Mrpl52** | -1.539451166 | 0.014637172 |
| 234 | **Ddt** | -1.541392635 | 0.008731198 |

| | | | |
|---|---|---|---|
| 235 | **Gnao1** | 1.760966807 | 0.015338549 |
| 236 | **Atp6v1e1** | 1.76017851 | 0.029259834 |
| 237 | **Serpinh1** | 1.754584594 | 0.001411055 |
| 238 | **Lars1** | 1.753984168 | 0.003876002 |
| 239 | **Pabpc1** | 1.748536661 | 0.002108928 |
| 240 | **Ybx3** | 1.740900595 | 0.010028856 |
| 241 | **Snrnp70** | 1.738873297 | 0.010527375 |
| 242 | **Myl6** | 1.737224843 | 0.020332997 |
| 243 | **Prmt1** | 1.735584321 | 0.00600137 |
| 244 | **Copg2** | 1.731653922 | 0.026125184 |
| 245 | **Akt1** | 1.731239687 | 0.003723688 |
| 246 | **Lamp2** | 1.72676761 | 0.002258852 |
| 247 | **Ubqln2** | 1.726132093 | 0.00181256 |
| 248 | **Gmppa** | 1.725203215 | 0.005073924 |
| 249 | **Lamp1** | 1.721389516 | 0.002506478 |
| 250 | **H2-D1** | 1.718298424 | 0.001715592 |
| 251 | **Fus** | 1.715602123 | 0.002621846 |
| 252 | **Calu** | 1.710225473 | 0.002304252 |
| 253 | **Copg1** | 1.709235689 | 0.002822823 |
| 254 | **Rbm3** | 1.708379515 | 0.030839023 |
| 255 | **Copb2** | 1.706781982 | 0.002257246 |
| 256 | **H1-3** | 1.701588357 | 0.001411055 |
| 257 | **Iah1** | 1.700244907 | 0.004100534 |
| 258 | **Syk** | 1.695129402 | 0.029976837 |
| 259 | **Arpc2** | 1.693911413 | 0.001652328 |
| 260 | **Vim** | 1.679679287 | 0.00229412 |
| 261 | **Esyt1** | 1.679348504 | 0.001723939 |
| 262 | **Vwf** | 1.677551402 | 0.003544625 |
| 263 | **Lpp** | 1.675277168 | 0.010268394 |
| 264 | **Capza1** | 1.675271991 | 0.001715592 |
| 265 | **Fabp5** | 1.670459174 | 0.001411055 |
| 266 | **Manf** | 1.664275246 | 0.001859971 |
| 267 | **Aars1** | 1.657321297 | 0.013772291 |
| 268 | **Ftl1** | 1.654200561 | 0.002434267 |
| 269 | **Eif3b** | 1.653296693 | 0.001652328 |
| 270 | **Fgb** | 1.645843131 | 0.00229412 |
| 271 | **Sh3bgrl** | 1.644302658 | 0.001715592 |
| 272 | **Vasp** | 1.642013859 | 0.005694914 |
| 273 | **Tomm34** | 1.641738838 | 0.041868774 |
| 274 | **Arhgap1** | 1.638424517 | 0.003700485 |
| 275 | **Ilf3** | 1.635552798 | 0.032006845 |
| 276 | **Apex1** | 1.632838973 | 0.007368793 |
| 277 | **Srprb** | 1.626632409 | 0.029282088 |
| 278 | **Tmsb4x** | 1.624552709 | 0.002228601 |
| 279 | **Hnrnpab** | 1.622547162 | 0.001859971 |
| 280 | **Septin9** | 1.619867741 | 0.008194002 |
| 281 | **Psmb8** | 1.619583895 | 0.002087105 |
| 282 | **Hprt1** | 1.616618866 | 0.002970429 |

| | | | |
|---|---|---|---|
| 235 | **Aifm1** | -1.543773 | 0.002228601 |
| 236 | **Ndufa11** | -1.546489829 | 0.003877091 |
| 237 | **Ndufv2** | -1.547677936 | 0.003184261 |
| 238 | **Pdp1** | -1.549157566 | 0.002108928 |
| 239 | **Atp2a2** | -1.549339169 | 0.001784651 |
| 240 | **Krt42** | -1.550952284 | 0.006852624 |
| 241 | **Pygm** | -1.552657024 | 0.003157642 |
| 242 | **Mtco2** | -1.559110643 | 0.012229546 |
| 243 | **Mcat** | -1.55911484 | 0.020099706 |
| 244 | **Ogdh** | -1.560574491 | 0.00680545 |
| 245 | **Chchd3** | -1.562399191 | 0.004082264 |
| 246 | **Mrpl22** | -1.562764925 | 0.013772291 |
| 247 | **Echdc** | -1.569604988 | 0.005073924 |
| 248 | **Nipsnap2** | -1.576405781 | 0.005203129 |
| 249 | **L2hgdh** | -1.579582024 | 0.001670911 |
| 250 | **Myot** | -1.580379557 | 0.01525981 |
| 251 | **mCoq6 m** | -1.580637749 | 0.010744841 |
| 252 | **Pdk1** | -1.582024526 | 0.009923547 |
| 253 | **Ttc38** | -1.585140471 | 0.046543572 |
| 254 | **Cdh13** | -1.5913135 | 0.00229412 |
| 255 | **Ndufaf4** | -1.596141294 | 0.004677152 |
| 256 | **Dlat** | -1.597233336 | 0.002611587 |
| 257 | **Ech1** | -1.599229834 | 0.002108928 |
| 258 | **mOxct1 m** | -1.600243966 | 0.007006066 |
| 259 | **mTrim54** | -1.601519224 | 0.045409603 |
| 260 | **Ldhb** | -1.603197944 | 0.003115387 |
| 261 | **Aldh4a1** | -1.603894031 | 0.001419231 |
| 262 | **Tufmd** | -1.604476898 | 0.002108928 |
| 263 | **Nudt8** | -1.60488777 | 0.029035015 |
| 264 | **Tgfb1i1** | -1.605686683 | 0.018639968 |
| 265 | **Hadhb** | -1.606413317 | 0.002257246 |
| 266 | **Uqcr11** | -1.608551413 | 0.037084067 |
| 267 | **Fahd2** | -1.609709917 | 0.033383574 |
| 268 | **Srpx** | -1.609756557 | 0.029220756 |
| 269 | **Acsf3** | -1.614176521 | 0.033121369 |
| 270 | **Gstk1** | -1.614988094 | 0.001723939 |
| 271 | **Naxd** | -1.616421668 | 0.005668554 |
| 272 | **Ndufs1** | -1.617504031 | 0.002840352 |
| 273 | **Acaa2** | -1.617721212 | 0.001582284 |
| 274 | **Bphl** | -1.619922218 | 0.009246952 |
| 275 | **Ndufv1** | -1.621157486 | 0.001723939 |
| 276 | **Mrpl21** | -1.622257231 | 0.00298111 |
| 277 | **Ndrg2** | -1.626781149 | 0.003438574 |
| 278 | **Mtch2** | -1.631397782 | 0.002173193 |
| 279 | **Crat** | -1.632984018 | 0.00206644 |
| 280 | **Mpv17** | -1.636075629 | 0.027480621 |
| 281 | **Bckdha** | -1.636587939 | 0.001789432 |
| 282 | **Fh** | -1.636817326 | 0.002166624 |

| | | | |
|---|---|---|---|
| 283 | **Pycr2** | 1.613369778 | 0.012314018 |
| 284 | **Pxdn** | 1.612445135 | 0.003146495 |
| 285 | **Tubb2a** | 1.605747666 | 0.001715592 |
| 286 | **Prpf8** | 1.602891872 | 0.005699789 |
| 287 | **F13a1** | 1.602545671 | 0.002659633 |
| 288 | **Irgm1** | 1.595151273 | 0.001419231 |
| 289 | **Tgm2** | 1.591256534 | 0.002036073 |
| 290 | **Ace** | 1.584943356 | 0.011983905 |
| 291 | **Nomo1** | 1.580579675 | 0.012627545 |
| 292 | **Ywhah** | 1.580044369 | 0.00345321 |
| 293 | **Flnc** | 1.578519916 | 0.001723939 |
| 294 | **Cul1** | 1.573860968 | 0.001936982 |
| 295 | **Cpsf6** | 1.572840736 | 0.005114561 |
| 296 | **Mapre1** | 1.566549518 | 0.005073924 |
| 297 | **Aprt** | 1.564047329 | 0.005548437 |
| 298 | **Flna** | 1.560086254 | 0.003024906 |
| 299 | **Api5** | 1.554575749 | 0.002355372 |
| 300 | **Calr** | 1.551910669 | 0.002840352 |
| 301 | **Srp14** | 1.551125213 | 0.040884859 |
| 302 | **Sirt2** | 1.549928442 | 0.011304084 |
| 303 | **Kpna4** | 1.548079566 | 0.019991786 |
| 304 | **Ncl** | 1.53802485 | 0.001876125 |
| 305 | **Map2k1** | 1.536025438 | 0.00432025 |
| 306 | **Lsm8** | 1.53311132 | 0.041478242 |
| 307 | **Vamp8** | 1.531959846 | 0.015132793 |
| 308 | **Dpp3** | 1.531210356 | 0.002228601 |
| 309 | **Lbr** | 1.529743508 | 0.001915504 |
| 310 | **Mtdh** | 1.527445448 | 0.049376088 |
| 311 | **Vat1** | 1.527092751 | 0.002840352 |
| 312 | **Pdcd10** | 1.523771158 | 0.046065084 |
| 313 | **Khdrbs1** | 1.520560926 | 0.003587143 |
| 314 | **Copa** | 1.519475461 | 0.002108928 |
| 315 | **Rps27** | 1.518209819 | 0.014458496 |
| 316 | **Vtn** | 1.513257864 | 0.02730931 |
| 317 | **Ankrd17** | 1.510687022 | 0.03728351 |
| 318 | **Stt3b** | 1.51038079 | 0.008323225 |
| 319 | **Dazap1** | 1.504644874 | 0.008384234 |
| 320 | **Tubb5** | 1.487301882 | 0.004357056 |
| 321 | **Sec23a** | 1.486924865 | 0.007190919 |
| 322 | **Cnot1** | 1.480964912 | 0.031313704 |
| 323 | **Rps14** | 1.479788206 | 0.003571267 |
| 324 | **Sec31a** | 1.478891456 | 0.001911421 |
| 325 | **Eif5b** | 1.476600573 | 0.003601707 |
| 326 | **Usp4** | 1.470715435 | 0.002840352 |
| 327 | **Itgb2** | 1.461430467 | 0.009066564 |
| 328 | **Tbcb** | 1.45565217 | 0.00575296 |
| 329 | **Eif3i** | 1.455575785 | 0.003700485 |
| 330 | **Arg1** | 1.452864907 | 0.012892768 |

| | | | |
|---|---|---|---|
| 283 | **Slc25a13** | -1.637172175 | 0.003758485 |
| 284 | **Ndufa13** | -1.637313076 | 0.008552061 |
| 285 | **Selenbp1** | -1.638860302 | 0.004697639 |
| 286 | **Atp5f1a** | -1.64015685 | 0.001876125 |
| 287 | **Aco2** | -1.640761786 | 0.002228601 |
| 288 | **Ndufb10** | -1.644314153 | 0.002685551 |
| 289 | **Rmdn1** | -1.647703033 | 0.029259834 |
| 290 | **Tars3** | -1.650199403 | 0.042369332 |
| 291 | **Ndufa4** | -1.658669981 | 0.002228601 |
| 292 | **Cyc1** | -1.658866271 | 0.002058898 |
| 293 | **Nudt2** | -1.658872536 | 0.024019193 |
| 294 | **Spryd4** | -1.659975903 | 0.038106543 |
| 295 | **Gstm1** | -1.663926562 | 0.002933447 |
| 296 | **Prdx3** | -1.66543428 | 0.005607633 |
| 297 | **Grhpr** | -1.666889017 | 0.002642729 |
| 298 | **Sdha** | -1.670522345 | 0.002621846 |
| 299 | **Smarcd1** | -1.671445644 | 0.017029432 |
| 300 | **Eif4a2** | -1.672399929 | 0.007931601 |
| 301 | **Abcb8** | -1.673513529 | 0.002108928 |
| 302 | **Auh** | -1.674222187 | 0.00665093 |
| 303 | **Clybl** | -1.674255482 | 0.002126664 |
| 304 | **Ptcd3** | -1.677603798 | 0.00432025 |
| 305 | **Myh6** | -1.681451792 | 0.00222969 |
| 306 | **Ndufs8** | -1.683798423 | 0.027436727 |
| 307 | **Cbr4** | -1.684527866 | 0.005389611 |
| 308 | **Lactb** | -1.685477924 | 0.029707918 |
| 309 | **Ndufb4** | -1.689441254 | 0.001691139 |
| 310 | **Txnrd2** | -1.694714074 | 0.01525981 |
| 311 | **Tnnt2** | -1.698199106 | 0.001789432 |
| 312 | **Nnt** | -1.698920906 | 0.001652328 |
| 313 | **Pcca** | -1.699810064 | 0.001914341 |
| 314 | **Sucla2** | -1.700456237 | 0.001715592 |
| 315 | **Acsl1** | -1.701115756 | 0.001852133 |
| 316 | **Bcs1l** | -1.702277213 | 0.039189805 |
| 317 | **Sars2** | -1.70378272 | 0.020886163 |
| 318 | **Uqcrc2** | -1.704252644 | 0.002355372 |
| 319 | **Iars2** | -1.707236662 | 0.011483911 |
| 320 | **Coq9** | -1.707312353 | 0.003198889 |
| 321 | **Sdhc** | -1.713962432 | 0.005389611 |
| 322 | **Mccc1** | -1.715262137 | 0.001723939 |
| 323 | **Krt10** | -1.716250603 | 0.006284673 |
| 324 | **Idh3g** | -1.716525138 | 0.008838398 |
| 325 | **Smim4** | -1.716857036 | 0.023444652 |
| 326 | **Plin3** | -1.718860292 | 0.014458496 |
| 327 | **Gstm7** | -1.720269019 | 0.006183988 |
| 328 | **Etfrf1** | -1.725920746 | 0.024277598 |
| 329 | **Chchd6** | -1.726386477 | 0.020886163 |
| 330 | **Mtnd1** | -1.730272956 | 0.005304655 |

| | | | |
|---|---|---|---|
| 331 | **Fga** | 1.450312304 | 0.00229412 |
| 332 | **Eef1b** | 1.446351031 | 0.002794495 |
| 333 | **Cpq** | 1.44630497 | 0.002697184 |
| 334 | **Rnh1** | 1.443698096 | 0.003508536 |
| 335 | **Myl12b** | 1.43765223 | 0.003159247 |
| 336 | **Atp6v1b2** | 1.434669342 | 0.001913633 |
| 337 | **Tagln2** | 1.429037155 | 0.005374572 |
| 338 | **Fgg** | 1.422819883 | 0.002703907 |
| 339 | **Wfs1** | 1.422746743 | 0.035611323 |
| 340 | **Fam98a** | 1.422468663 | 0.011029063 |
| 341 | **Snrpa** | 1.420756897 | 0.005073924 |
| 342 | **Abcf1** | 1.41952447 | 0.003198889 |
| 343 | **Pepd** | 1.419234935 | 0.023249824 |
| 344 | **Myh9** | 1.418773107 | 0.003536193 |
| 345 | **Lgals1** | 1.412311454 | 0.006484299 |
| 346 | **Ncstn** | 1.410864869 | 0.012683584 |
| 347 | **Rbm14** | 1.40891285 | 0.008790679 |
| 348 | **Nap1l1** | 1.403249707 | 0.003437428 |
| 349 | **Nmi** | 1.397699662 | 0.011570466 |
| 350 | **Tap2** | 1.39698861 | 0.038620529 |
| 351 | **Ap3b1** | 1.395157331 | 0.014644977 |
| 352 | **Rps5** | 1.393164145 | 0.002974051 |
| 353 | **Numa1** | 1.387846141 | 0.00628681 |
| 354 | **Hspb7** | 1.384781823 | 0.004534312 |
| 355 | **S100a10** | 1.380760028 | 0.009908977 |
| 356 | **Col6a1** | 1.380727161 | 0.002093653 |
| 357 | **Sec61b** | 1.374224088 | 0.006649944 |
| 358 | **Rab8b** | 1.369900276 | 0.011696402 |
| 359 | **Gsn** | 1.364928568 | 0.002355372 |
| 360 | **Chmp3** | 1.364767872 | 0.025777431 |
| 361 | **Hnrnpu** | 1.364513478 | 0.001913633 |
| 362 | **Fkbp8** | 1.364203989 | 0.005711621 |
| 363 | **Ykt6** | 1.359481114 | 0.035807362 |
| 364 | **Rpn1** | 1.355142171 | 0.001928588 |
| 365 | **B2m** | 1.352470284 | 0.005073924 |
| 366 | **Msn** | 1.351531438 | 0.002370797 |
| 367 | **Serpinf1** | 1.347808258 | 0.007121008 |
| 368 | **Cpne1** | 1.347478534 | 0.011814939 |
| 369 | **Uso1** | 1.340785324 | 0.003645066 |
| 370 | **Itih2** | 1.339538322 | 0.006552946 |
| 371 | **Srsf3** | 1.335456109 | 0.006552946 |
| 372 | **Fam177a1** | 1.328274925 | 0.037114226 |
| 373 | **Apoe** | 1.328013746 | 0.008926329 |
| 374 | **Aip** | 1.324349108 | 0.007815228 |
| 375 | **Erap1** | 1.320065349 | 0.00575296 |
| 376 | **Adprh** | 1.319049745 | 0.038106543 |
| 377 | **Itih4** | 1.31643578 | 0.003684766 |
| 378 | **Eif2a** | 1.315897518 | 0.004267995 |

| | | | |
|---|---|---|---|
| 331 | **Atl2** | -1.731939945 | 0.030539797 |
| 332 | **Slc25a3** | -1.736775216 | 0.012032687 |
| 333 | **Ndufs3** | -1.74229226 | 0.001411055 |
| 334 | **Opa1** | -1.742374936 | 0.002087105 |
| 335 | **Katnb1** | -1.744842176 | 0.002840352 |
| 336 | **Atp5pf** | -1.745031394 | 0.020099706 |
| 337 | **Prkar2a** | -1.748672239 | 0.018449622 |
| 338 | **Mrpl47** | -1.749532943 | 0.022918342 |
| 339 | **Lrpprc** | -1.749938348 | 0.002207375 |
| 340 | **Mrps27** | -1.750968349 | 0.005950447 |
| 341 | **Atp5f1b** | -1.75578212 | 0.004106883 |
| 342 | **Aldh6a1** | -1.758345324 | 0.006064076 |
| 343 | **Mrps22** | -1.759419812 | 0.004357056 |
| 344 | **Krt16** | -1.759579329 | 0.009540221 |
| 345 | **Acat1** | -1.762675902 | 0.001723939 |
| 346 | **Ak4** | -1.766884544 | 0.01314478 |
| 347 | **Dhrs11** | -1.772411155 | 0.009863002 |
| 348 | **Atp5mg** | -1.777891532 | 0.003437428 |
| 349 | **Mrps36** | -1.77825415 | 0.003157642 |
| 350 | **Brf2** | -1.779089505 | 0.022383825 |
| 351 | **C1qbp** | -1.779418091 | 0.008576062 |
| 352 | **Pdh** | -1.779992944 | 0.001864912 |
| 353 | **Mrrf** | -1.783195372 | 0.002490104 |
| 354 | **Prdx2** | -1.783509881 | 0.006644354 |
| 355 | **Mpc2** | -1.787891067 | 0.026876696 |
| 356 | **Lipe** | -1.789987124 | 0.024274536 |
| 357 | **Letm1** | -1.791292905 | 0.003441797 |
| 358 | **Atp5mf** | -1.794915262 | 0.017373406 |
| 359 | **Cs** | -1.79682407 | 0.009954278 |
| 360 | **Slc25a12** | -1.807817919 | 0.001411055 |
| 361 | **Tspan9** | -1.808890749 | 0.029122742 |
| 362 | **Slc16a1** | -1.810616112 | 0.006250288 |
| 363 | **Mtnd4** | -1.810814343 | 0.04070394 |
| 364 | **Iba57** | -1.81407939 | 0.00527745 |
| 365 | **Vwa8** | -1.816828654 | 0.002228601 |
| 366 | **Ndufs2** | -1.82622666 | 0.002967451 |
| 367 | **Hibadh** | -1.828387072 | 0.002207375 |
| 368 | **Nudt7** | -1.833276746 | 0.005082952 |
| 369 | **Ckmt2** | -1.836991417 | 0.001876125 |
| 370 | **Akr1e2** | -1.837034682 | 0.026608912 |
| 371 | **Plin4** | -1.837374721 | 0.002334416 |
| 372 | **Hadha** | -1.838827544 | 0.00355206 |
| 373 | **Tmod1** | -1.840827438 | 0.001927502 |
| 374 | **Cluh** | -1.841971226 | 0.007956575 |
| 375 | **Lyrm7** | -1.848669414 | 0.018825669 |
| 376 | **Ppcs** | -1.85357401 | 0.004534312 |
| 377 | **Mccc2** | -1.853883861 | 0.001784651 |
| 378 | **Acadl** | -1.861691603 | 0.001652328 |

| | | | |
|---|---|---|---|
| 379 | **Txnip** | 1.312657407 | 0.010055962 |
| 380 | **Tor1aip1** | 1.310101392 | 0.028341757 |
| 381 | **Snap23** | 1.308276712 | 0.018329795 |
| 382 | **Erh** | 1.308090166 | 0.006782685 |
| 383 | **Ube2m** | 1.306959259 | 0.017581569 |
| 384 | **Apoh** | 1.303750121 | 0.004445379 |
| 385 | **Orm1** | 1.300971685 | 0.045409603 |
| 386 | **Hmgb2** | 1.298497587 | 0.003922594 |
| 387 | **Pdia6** | 1.292567983 | 0.008385668 |
| 388 | **H2az1** | 1.291372589 | 0.005937974 |
| 389 | **Dad1** | 1.290725629 | 0.008576062 |
| 390 | **Psmb1** | 1.290686031 | 0.007642505 |
| 391 | **Txnrd1** | 1.285626584 | 0.002993465 |
| 392 | **Lyn** | 1.28547693 | 0.003441797 |
| 393 | **Tmx3** | 1.284750345 | 0.0120702 |
| 394 | **Ddx6** | 1.28164994 | 0.002355372 |
| 395 | **Pfn1** | 1.279722662 | 0.005835914 |
| 396 | **Clic4** | 1.279265663 | 0.043968204 |
| 397 | **Snu13** | 1.279065188 | 0.009001636 |
| 398 | **Gmppb** | 1.275709105 | 0.009273987 |
| 399 | **Snrpn** | 1.275483536 | 0.046437238 |
| 400 | **Rab5c** | 1.274787171 | 0.003797927 |
| 401 | **Dnm2** | 1.269869194 | 0.002792241 |
| 402 | **Fubp1** | 1.268590959 | 0.009001636 |
| 403 | **Rps6ka3** | 1.256930281 | 0.021619711 |
| 404 | **Srm** | 1.254271111 | 0.01314478 |
| 405 | **Anxa2** | 1.251586723 | 0.004683371 |
| 406 | **Fhl1** | 1.249863775 | 0.005389611 |
| 407 | **Hnrnpul2** | 1.248186854 | 0.015245017 |
| 408 | **Ddx5** | 1.247254266 | 0.004312154 |
| 409 | **Tpm4** | 1.245825188 | 0.004272328 |
| 410 | **Ppib** | 1.24497404 | 0.006071365 |
| 411 | **Npm1** | 1.244455015 | 0.030613138 |
| 412 | **Rhoa** | 1.241213167 | 0.005709066 |
| 413 | **Twf2** | 1.235090478 | 0.003645066 |
| 414 | **Ubr4** | 1.232830748 | 0.043924397 |
| 415 | **Eif2s3x** | 1.231992589 | 0.032385799 |
| 416 | **Ssb** | 1.229150057 | 0.002840352 |
| 417 | **Atox1** | 1.228496222 | 0.014458496 |
| 418 | **Rpl10** | 1.227079958 | 0.008956781 |
| 419 | **Snrpg** | 1.224343338 | 0.027157504 |
| 420 | **Pfkp** | 1.218282788 | 0.005389611 |
| 421 | **Hsp90b1** | 1.214748817 | 0.003157642 |
| 422 | **Flnb** | 1.213337899 | 0.006484299 |
| 423 | **Mars1** | 1.212974899 | 0.045230154 |
| 424 | **Hnrnph1** | 1.208006031 | 0.002921578 |
| 425 | **Capn2** | 1.190357818 | 0.005026849 |
| 426 | **Taldo1** | 1.183992307 | 0.004445379 |

| | | | |
|---|---|---|---|
| 379 | **Jph2** | -1.865112707 | 0.003198889 |
| 380 | **Nadk2** | -1.86899195 | 0.002355372 |
| 381 | **Kdm1a** | -1.875315726 | 0.010016444 |
| 382 | **Ndufa8** | -1.876957197 | 0.002937489 |
| 383 | **Myoz2** | -1.879445189 | 0.005389611 |
| 384 | **Maip1** | -1.881942058 | 0.025005872 |
| 385 | **Acot2** | -1.886468602 | 0.001652328 |
| 386 | **Crip2** | -1.888591181 | 0.009856785 |
| 387 | **Tmem65** | -1.891110956 | 0.00665093 |
| 388 | **Atp5po** | -1.891595177 | 0.0029388 |
| 389 | **Guf1** | -1.896603808 | 0.00680545 |
| 390 | **Krt73** | -1.896837892 | 0.007121008 |
| 391 | **Etfb** | -1.897908256 | 0.00229412 |
| 392 | **Timm44** | -1.899899416 | 0.005199169 |
| 393 | **Sh3bgr** | -1.914939484 | 0.021045125 |
| 394 | **Ndufa2** | -1.917464402 | 0.00423447 |
| 395 | **Asap2** | -1.919176317 | 0.015596598 |
| 396 | **Nebl** | -1.926396967 | 0.003758485 |
| 397 | **Mcee** | -1.93035201 | 0.010527375 |
| 398 | **Ryr2** | -1.931526389 | 0.002037933 |
| 399 | **Mdh2** | -1.933922575 | 0.002355372 |
| 400 | **Me3** | -1.940270393 | 0.003011038 |
| 401 | **Echs1** | -1.940341488 | 0.002166624 |
| 402 | **Ppa2** | -1.942351869 | 0.001715592 |
| 403 | **Acsl6** | -1.949114268 | 0.032177193 |
| 404 | **Acss1** | -1.951513782 | 0.00175894 |
| 405 | **Atpaf2** | -1.953542443 | 0.033875528 |
| 406 | **Ank2** | -1.961652041 | 0.029958435 |
| 407 | **Gja1** | -1.965055294 | 0.002471417 |
| 408 | **Isoc2a** | -1.96990381 | 0.03248888 |
| 409 | **Htra2** | -1.969946675 | 0.022052418 |
| 410 | **Coq3** | -1.971699656 | 0.001715592 |
| 411 | **Rab3a** | -1.973253008 | 0.005649341 |
| 412 | **Acadm** | -1.979091357 | 0.002129144 |
| 413 | **Etfa** | -1.981379458 | 0.001789432 |
| 414 | **Mrps35** | -1.983368587 | 0.006310413 |
| 415 | **Acacb** | -1.98703495 | 0.009602489 |
| 416 | **Atp1a2** | -1.988838128 | 0.002937489 |
| 417 | **Bcl2l13** | -1.990799106 | 0.011875199 |
| 418 | **Mrpl4** | -1.995378553 | 0.004088571 |
| 419 | **Lars2** | -1.9959694 | 0.031144652 |
| 420 | **Igbp1** | -1.997965164 | 0.031199898 |
| 421 | **Pc** | -2.003339274 | 0.021014405 |
| 422 | **Mmut** | -2.005835603 | 0.001723939 |
| 423 | **Slc9a3r2** | -2.006627736 | 0.025764045 |
| 424 | **Vars2** | -2.013939167 | 0.001784651 |
| 425 | **Ndufb6** | -2.023463217 | 0.00131716 |
| 426 | **Mtif2** | -2.025108979 | 0.014644977 |

| | | | |
|---|---|---|---|
| 427 | **Rpl15** | 1.183630197 | 0.003157642 |
| 428 | **Gna13** | 1.173276046 | 0.007761453 |
| 429 | **Sri** | 1.173232651 | 0.005699789 |
| 430 | **Ap1m1** | 1.172744975 | 0.029585646 |
| 431 | **Rps19** | 1.172367484 | 0.006199316 |
| 432 | **Tspo** | 1.16969782 | 0.045230154 |
| 433 | **Aga** | 1.165879136 | 0.022662261 |
| 434 | **Hnrnpc** | 1.165596586 | 0.029350423 |
| 435 | **Rps3a** | 1.165415097 | 0.008292097 |
| 436 | **Inpp5a** | 1.165287078 | 0.010905893 |
| 437 | **Eif3e** | 1.16286096 | 0.005324987 |
| 438 | **Bola2** | 1.161706839 | 0.045230154 |
| 439 | **Rplp1** | 1.157626551 | 0.008953341 |
| 440 | **C4b** | 1.153685889 | 0.008427858 |
| 441 | **Lrrc47** | 1.151741133 | 0.049054047 |
| 442 | **Nek7** | 1.147741049 | 0.022899313 |
| 443 | **Ddost** | 1.144124354 | 0.008052273 |
| 444 | **Zyx** | 1.143106674 | 0.013712958 |
| 445 | **Fau** | 1.139058404 | 0.026428368 |
| 446 | **Strap** | 1.136260391 | 0.01187943 |
| 447 | **Pa2g4** | 1.132914641 | 0.004639753 |
| 448 | **Nono** | 1.130438345 | 0.008133481 |
| 449 | **Etf1** | 1.120149437 | 0.013398365 |
| 450 | **Septin8** | 1.113286769 | 0.021619711 |
| 451 | **Rps4x** | 1.112497781 | 0.007829355 |
| 452 | **Gimap4** | 1.103737244 | 0.018351853 |
| 453 | **Ap2m1** | 1.099330506 | 0.006580593 |
| 454 | **Tax1bp3** | 1.098387322 | 0.020611673 |
| 455 | **Rpl19** | 1.096899836 | 0.026586215 |
| 456 | **Anxa5** | 1.092779776 | 0.006552946 |
| 457 | **Srsf2** | 1.089497123 | 0.005928307 |
| 458 | **Rps21** | 1.087790676 | 0.004673422 |
| 459 | **Arpc4** | 1.086931124 | 0.01196736 |
| 460 | **Rpl34** | 1.086808009 | 0.037578517 |
| 461 | **Eif4a1** | 1.084491638 | 0.004683371 |
| 462 | **Map1lc3a** | 1.081443891 | 0.022720604 |
| 463 | **Rps12** | 1.080923469 | 0.005463204 |
| 464 | **Gsr** | 1.070615206 | 0.006393349 |
| 465 | **Septin7** | 1.069912682 | 0.00694121 |
| 466 | **Myl9** | 1.069745919 | 0.006931118 |
| 467 | **Clint1** | 1.069729266 | 0.041103626 |
| 468 | **Mcam** | 1.069636541 | 0.008956781 |
| 469 | **Hnrnpa3** | 1.068452764 | 0.007926799 |
| 470 | **U2af2** | 1.067674232 | 0.008999625 |
| 471 | **Ctnnd1** | 1.066800846 | 0.011814939 |
| 472 | **Elavl1** | 1.064464262 | 0.021341239 |
| 473 | **Rer1** | 1.064103882 | 0.009201323 |
| 474 | **Uchl5** | 1.063238511 | 0.009142182 |

| | | | |
|---|---|---|---|
| 427 | **Ndufa5** | -2.032057907 | 0.009677682 |
| 428 | **Tnni3** | -2.03335948 | 0.003946323 |
| 429 | **Timmdc1** | -2.034414378 | 0.009913486 |
| 430 | **Tmod4** | -2.03503 | 0.002970429 |
| 431 | **Cd36** | -2.045631999 | 0.001411055 |
| 432 | **Atp5mj** | -2.047664367 | 0.026015707 |
| 433 | **Cpt1b** | -2.058666334 | 0.001897093 |
| 434 | **Sdr39u1** | -2.070509505 | 0.002340455 |
| 435 | **Hadh** | -2.073520096 | 0.005711589 |
| 436 | **Pnpt1** | -2.081615161 | 0.026428368 |
| 437 | **Myl3** | -2.090088297 | 0.003115387 |
| 438 | **Akr1b7** | -2.090962797 | 0.006671056 |
| 439 | **Ctnna3** | -2.099151788 | 0.001876125 |
| 440 | **Pdk2** | -2.121377497 | 0.001227373 |
| 441 | **Hsdl2** | -2.132048732 | 0.004100534 |
| 442 | **Pdha2** | -2.140621829 | 0.034240962 |
| 443 | **Hsd17b8** | -2.145531396 | 0.008263247 |
| 444 | **Mrps30** | -2.157683691 | 0.011225625 |
| 445 | **Myl2** | -2.157726366 | 0.001916552 |
| 446 | **Tmem126a** | -2.158038319 | 0.025322924 |
| 447 | **Mrpl33** | -2.162492354 | 0.017707735 |
| 448 | **Ndufa10** | -2.16811886 | 0.007060906 |
| 449 | **Endog** | -2.183719234 | 0.002542629 |
| 450 | **Inmt** | -2.192884859 | 0.020347515 |
| 451 | **Ckm** | -2.202126659 | 0.003438574 |
| 452 | **Serpina1e** | -2.221065146 | 0.01229201 |
| 453 | **Slc27a1** | -2.226855389 | 0.003146495 |
| 454 | **Myzap** | -2.231182192 | 0.017535593 |
| 455 | **Ndufa9** | -2.244230917 | 0.004611931 |
| 456 | **Far1** | -2.245258994 | 0.007805047 |
| 457 | **Fgf1** | -2.246038701 | 0.005061734 |
| 458 | **Hdhd5** | -2.252538664 | 0.011814939 |
| 459 | **Mtnd2** | -2.260084381 | 0.024742012 |
| 460 | **Acadsb** | -2.26252373 | 0.001411055 |
| 461 | **Bcam** | -2.273275323 | 0.001227373 |
| 462 | **Mrpl39** | -2.277127042 | 0.014435403 |
| 463 | **Tefm** | -2.277357566 | 0.029976837 |
| 464 | **Aldh1a1** | -2.278603506 | 0.006964891 |
| 465 | **Ldb3** | -2.282240802 | 0.001411055 |
| 466 | **Acad10** | -2.285216583 | 0.001227373 |
| 467 | **Uqcc2** | -2.28552287 | 0.019826301 |
| 468 | **Kif15** | -2.295357649 | 0.030107934 |
| 469 | **Idh2** | -2.302636396 | 0.002547795 |
| 470 | **Aqp1** | -2.308311845 | 0.008020207 |
| 471 | **Ptgr2** | -2.310489336 | 0.005451226 |
| 472 | **Krt6b** | -2.311434884 | 0.04756289 |
| 473 | **Spta1** | -2.316474881 | 0.020041509 |
| 474 | **Acp6** | -2.319376873 | 0.005073924 |

| | | | |
|---|---|---|---|
| 475 | **Mob1b** | 1.061567424 | 0.020715883 |
| 476 | **Actn4** | 1.052025326 | 0.01229201 |
| 477 | **Lsm4** | 1.05184089 | 0.040791457 |
| 478 | **Snrpd3** | 1.049260308 | 0.013772291 |
| 479 | **Vwa5a** | 1.048711902 | 0.005776718 |
| 480 | **Dctn4** | 1.048703582 | 0.009323728 |
| 481 | **Gnpda1** | 1.047354244 | 0.004456677 |
| 482 | **Ddx39b** | 1.046601442 | 0.006651092 |
| 483 | **Pdcd6ip** | 1.033390894 | 0.006338526 |
| 484 | **Rps11** | 1.031176733 | 0.011478157 |
| 485 | **Ptms** | 1.029691056 | 0.012997067 |
| 486 | **Prrc1** | 1.028164049 | 0.009494331 |
| 487 | **Rpl35a** | 1.023349914 | 0.047703447 |
| 488 | **Ruvbl1** | 1.021238051 | 0.017643875 |
| 489 | **Iars1** | 1.019559343 | 0.007956575 |
| 490 | **Caprin1** | 1.012944156 | 0.015132793 |
| 491 | **Snrpf** | 1.007333038 | 0.009954278 |
| 492 | **Dek** | 1.007054277 | 0.004962052 |
| 493 | **Cst3** | 1.005484465 | 0.00628681 |
| 494 | **Ndrg1** | 1.003793517 | 0.049460654 |
| 495 | **Eif3a** | 1.001763655 | 0.009677682 |

| | | | |
|---|---|---|---|
| 475 | **Aoc3** | -2.330108677 | 0.005389611 |
| 476 | **Mmab** | -2.34147371 | 0.015305581 |
| 477 | **Mavs** | -2.351996648 | 0.002228601 |
| 478 | **Fhod3** | -2.35412972 | 0.002041831 |
| 479 | **Tmem38a** | -2.377207463 | 0.008050904 |
| 480 | **Mrpl10** | -2.41062825 | 0.036790759 |
| 481 | **Casq2** | -2.416983083 | 0.001366147 |
| 482 | **Selenbp2** | -2.419535583 | 0.001419231 |
| 483 | **Mtfp1** | -2.425983253 | 0.004673422 |
| 484 | **Sptb** | -2.43378399 | 0.020116815 |
| 485 | **Epb41** | -2.444881693 | 0.036815974 |
| 486 | **Ank1** | -2.444924761 | 0.029493911 |
| 487 | **Hint2** | -2.462270773 | 0.001199851 |
| 488 | **Gstz1** | -2.465833313 | 0.001715592 |
| 489 | **Krt79** | -2.471841333 | 0.042369332 |
| 490 | **Tmem143** | -2.471903216 | 0.002106262 |
| 491 | **Fbp2** | -2.477893633 | 0.010234097 |
| 492 | **Cmya5** | -2.478303748 | 0.001411055 |
| 493 | **Mrpl53** | -2.493752844 | 0.034897143 |
| 494 | **Gpt** | -2.502103808 | 0.003009159 |
| 495 | **Mgst3** | -2.542882744 | 0.003986567 |
| 496 | **Cacna2d1** | -2.560769945 | 0.001411055 |
| 497 | **Slc30a9** | -2.562741374 | 0.011788604 |
| 498 | **Sirt3** | -2.56343943 | 0.011209309 |
| 499 | **Slc2a4** | -2.577464618 | 0.001652328 |
| 500 | **Dhdh** | -2.598250491 | 0.005073924 |
| 501 | **Cox20** | -2.626761828 | 0.003249071 |
| 502 | **Coq8a** | -2.631238937 | 0.001729565 |
| 503 | **Gcat** | -2.633993705 | 0.011612404 |
| 504 | **Tmem177** | -2.664703041 | 0.015004261 |
| 505 | **Slc4a1** | -2.677979252 | 0.037023178 |
| 506 | **Uqcc1** | -2.686371275 | 0.022824156 |
| 507 | **Nt5c1a** | -2.69672827 | 0.044998491 |
| 508 | **Mrps5** | -2.699111876 | 0.033395422 |
| 509 | **Ca2** | -2.733239922 | 0.006484299 |
| 510 | **Mterf2** | -2.738166865 | 0.002840352 |
| 511 | **Ppp1r3a** | -2.763084233 | 0.001723939 |
| 512 | **A2m** | -2.780739652 | 0.001911421 |
| 513 | **Fundc1** | -2.785761993 | 0.016351034 |
| 514 | **Ucp3** | -2.803798574 | 0.006444507 |
| 515 | **Adh1** | -2.838886265 | 0.002108928 |
| 516 | **Ces1d** | -2.84686589 | 0.002207375 |
| 517 | **Acad11** | -2.864880178 | 0.001227373 |
| 518 | **Nudt13** | -2.869980131 | 0.03775942 |
| 519 | **Rtn2** | -2.881138891 | 0.001587411 |
| 520 | **Perm1** | -3.020784808 | 0.00896379 |
| 521 | **Bpgm** | -3.071094498 | 0.01362999 |
| 522 | **Ndufaf7** | -3.083896763 | 0.003617644 |

| | | | |
|---|---|---|---|
| 523 | **Armc1** | -3.143425809 | 0.010597493 |
| 524 | **Gpcpd1** | -3.194128737 | 0.001477059 |
| 525 | **Ca1** | -3.236000785 | 0.00486635 |
| 526 | **Art3** | -3.310486076 | 0.001876125 |
| 527 | **Pm20d2** | -3.332954658 | 0.001723939 |
| 528 | **Supv3l1** | -3.365926414 | 0.003877091 |
| 529 | **Pds5b** | -3.401677848 | 0.013774942 |
| 530 | **Ppp1r14c** | -3.412939474 | 0.00229412 |
| 531 | **Slc25a42** | -3.452560372 | 0.001227373 |
| 532 | **Fn3k** | -3.490237075 | 0.001789432 |
| 533 | **Ears2** | -3.604692549 | 0.001411055 |
| 534 | **Med31** | -3.627477786 | 0.002840352 |
| 535 | **Mapt** | -3.645429189 | 0.001419231 |
| 536 | **Mars2** | -3.735639024 | 0.006867105 |
| 537 | **Mbd2** | -3.910390255 | 0.005424337 |
| 538 | **Fblim1** | -4.0257472 | 0.001411055 |
| 539 | **Adck1** | -4.045278338 | 0.001419231 |
| 540 | **Mff** | -4.12181551 | 0.001723939 |
| 541 | **Plin5** | -4.780225435 | 0.001227373 |
| 542 | **Hbb-b2** | -5.850826048 | 0.001961005 |

**Supplementary Table 5.** List of downregulated proteins with the molecular function of DNA or RNA binding and transcriptional or translational regulator function

| Downregulated Proteins | Molecular Function | | | | |
|---|---|---|---|---|---|
| **Naca** | enables DNA binding | transcription coactivator activity | protein binding | enables unfolded protein binding | |
| **Mrpl12** | enables RNA binding | enables mRNA binding | enables structural constituent of ribosome | enables protein binding | |
| **Csrp1** | enables RNA binding | enables protein binding | enables zinc ion binding | enables structural constituent of muscle | enables actinin binding |
| **Alyref** | nucleic acid binding | enables RNA binding | enables mRNA binding | enables protein binding | enables C5-methylcytidine-containing RNA binding |
| **Pdcd5** | enables DNA binding | enables protein binding | enables heparin binding | enables acetyltransferase activator activity | enables beta-tubulin binding |
| **Trap1** | nucleotide binding | enables RNA binding | enables tumor necrosis factor receptor binding | enables protein binding | enables ATP binding |
| **Prdx5** | enables RNA polymerase III transcription regulatory region sequence-specific DNA binding | enables peroxidase activity | enables protein binding | enables thioredoxin peroxidase activity | antioxidant activity |
| **Glrx3** | enables RNA binding | enables protein kinase C binding | enables protein binding | enables identical protein binding | enables metal ion binding |
| **Ptges2** | enables DNA binding | enables protein binding | enables lyase activity | isomerase activity | enables heme binding |
| **Smyd1** | enables DNA binding | enables transcription corepressor activity | enables protein binding | methyltransferase activity | transferase activity |
| **Hsd17b10** | enables tRNA binding | enables RNA binding | enables 3-hydroxyacyl-CoA dehydrogenase activity | enables estradiol 17-beta-dehydrogenase activity | enables protein binding |
| ***Got2*** | enables RNA binding | catalytic activity | enables L-aspartate:2-oxoglutarate aminotransferase activity | transaminase activity | enables kynurenine-oxoglutarate transaminase activity |
| **Lonp1** | nucleotide binding | enables mitochondrial promoter sequence-specific DNA binding | DNA binding | enables single-stranded DNA binding | enables single-stranded RNA binding |

| | | | | | |
|---|---|---|---|---|---|
| **Huwe1** | enables DNA binding | enables RNA binding | enables ubiquitin-protein transferase activity | enables protein binding | transferase activity |
| **Gfm1** | nucleotide binding | enables RNA binding | enables translation elongation factor activity | enables GTPase activity | enables protein binding |
| **Mrpl28** | enables RNA binding | enables structural constituent of ribosome | enables protein binding | | |
| **Cast** | enables RNA binding | enables endopeptidase inhibitor activity | cysteine-type endopeptidase inhibitor activity | cysteine-type endopeptidase inhibitor activity | enables calcium-dependent cysteine-type endopeptidase inhibitor activity |
| **Grsf1** | nucleic acid binding | enables RNA binding | enables mRNA binding | enables protein binding | |
| **Lpin1** | enables transcription coactivator activity | enables protein binding | enables phosphatidate phosphatase activity | hydrolase activity | |
| **Hmga1** | enables RNA polymerase II cis-regulatory region sequence-specific DNA binding | enables cis-regulatory region sequence-specific DNA binding | enables transcription coregulator binding | enables DNA binding | enables minor groove of adenine-thymine-rich DNA binding |
| **Mrpl37** | enables RNA binding | enables structural constituent of ribosome | | | |
| **Tfam** | enables transcription cis-regulatory region binding | enables mitochondrial promoter sequence-specific DNA binding | enables transcription coactivator binding | DNA binding | enables chromatin binding |
| **Prkaa2** | nucleotide binding | enables chromatin binding | enables protein kinase activity | enables protein serine/threonine kinase activity | enables AMP-activated protein kinase activity |
| **Taco1** | enables mRNA binding | enables protein binding | enables rRNA binding | enables mitochondrial ribosome binding | |
| **Upf2** | enables RNA binding | enables protein binding | enables telomeric DNA binding | enables molecular adaptor activity | |
| **Tsfm** | enables RNA binding | enables translation elongation factor activity | enables protein binding | | |
| **Slc25a11** | enables RNA binding | enables protein binding | enables sulfate transmembrane transporter activity | enables thiosulfate transmembrane transporter activity | enables oxaloacetate transmembrane transporter activity |
| **Fkbp3** | enables RNA binding | enables peptidyl-prolyl cis-trans isomerase activity | enables protein binding | enables FK506 binding | isomerase activity |

| | | | | | |
|---|---|---|---|---|---|
| **Timm50** | enables RNA binding | enables phospho protein phosphatase activity | enables protein serine/threonine phosphatase activity | enables protein tyrosine phosphatase activity | enables interleukin -2 receptor binding |
| **Mrps26** | enables RNA binding | | | | |
| **Mlip** | enables transcri ption corepressor activity | protein binding | enables lamin binding | | |
| **Immt** | enables RNA binding | enables protein binding | | | |
| **Nqo1** | enables RNA binding | enables NAD(P) H dehydro- genase (quinone)activit y | enables cytochr ome-b5 reductase activity, acting on NAD(P)H | enables superoxide dismutase activity | enables protein binding |
| **Mtres1** | enables tRNA binding | enables RNA binding | enables protein binding | enables ribosomal large subunit binding | |
| **Atp5f1c** | enables RNA binding | enables protein binding | contributes_to ATP hydrolysis activity | contributes_to proton -transporting ATP synthase activity, rotational mechanism | |
| **Aifm1** | enables DNA binding | enables NADH dehydrogenase activity | enables protein binding | enables NAD(P)H oxidase H2O2-forming activity | oxidoreductase activity |
| **Mcat** | enables RNA binding | enables fatty acid synthase activity | enables [acyl-carrier-protein] S-malonyltransfer ase activity | S-malonyltransferase activity | transferase activity |
| **Mrpl22** | enables RNA binding | enables structur al constituent of ribosome | | | |
| **Tgfb1i1** | enables transcri ption coregulator activity | enables transcri ption co-activator activity | enables protein binding | enables metal ion binding | enables Roundabo ut binding |
| **Hadhb** | enables RNA binding | enables 3-hydroxyacyl-CoA dehydrogenase activity | enables acetyl-CoA C-acety-ltransferase activity | enables acetyl-CoA C-acyltransferase activity | enables enoyl-CoA hydratase activity |
| **Acaa2** | enables RNA binding | enables acetyl-CoA C-acetyl-transferase activity | enables acetyl-CoA hydrolase activity | enables acetyl-CoA C-acyltransferase activity | enables protein binding |
| **Mrpl21** | enables RNA binding | enables structur al constituent of ribosome | | | |
| **Atp5f1a** | nucleotide binding | enables protease binding | enables RNA binding | enables protein binding | enables ATP binding |
| **Smarcd1** | enables chroma tin binding | enables transcri ption coregulator activity | enables transcri ption coactivator activity | enables signaling receptor binding | enables protein binding |

| | | | | | |
|---|---|---|---|---|---|
| **Eif4a2** | nucleotide binding | nucleic acid binding | enables RNA binding | enables RNA helicase activity | enables translation initiation factor activity |
| **Auh** | RNA binding | enables mRNA 3'-UTR binding | catalytic activity | enables enoyl-CoA hydratase activity | enables methylglut aconyl-CoA hydratase activity |
| **Ptcd3** | enables RNA binding | enables protein binding | enables rRNA binding | enables ribosomal small subunit binding | |
| **Lrpprc** | DNA binding | enables single-stranded DNA binding | enables RNA binding | enables mRNA 3'-UTR binding | enables protein binding |
| **Mrps27** | enables tRNA binding | RNA binding | enables protein binding | enables rRNA binding | enables mitochond rial ribosome binding |
| **Aldh6a1** | enables fatty-acyl-CoA binding | enables RNA binding | enables methyl malonate semia-ldehyde dehydrogenase (acylating) activity | oxidoreductase activity | oxidoreductase activity, acting on the aldehyde or oxo group of donors, NAD or NADP as acceptor |
| **C1qbp** | enables comple ment component C1q complex binding | enables transcri ption corepressor activity | enables mRNA binding | enables protein kinase C binding | enables protein binding |
| **Cs** | enables RNA binding | enables citrate (Si)-synthase activity | transferase activity | citrate synthase activity | acyltransferase activity, acyl groups converted into alkyl on transfer |
| **Iba57** | enables RNA binding | enables protein binding | enables transfer ase activity | | |
| **Cluh** | RNA binding | enables mRNA binding | | | |
| **Mdh2** | enables RNA binding | catalytic activity | oxidoreductase activity | malate dehydrogenase activity | oxidoreductase activity, acting on the CH-OH group of donors, NAD or NADP as acceptor |
| **Mrps35** | enables RNA binding | enables structur al constituent of ribosome | | | |
| **Mrpl4** | enables RNA binding | enables structur al constituent of ribosome | enables protein binding | | |
| **Mtif2** | nucleotide binding | enables RNA binding | enables translati on initiation factor activity | enables GTPase activity | |
| **Pnpt1** | enables 3'-5'-exoribonucleas e activity | nucleic acid binding | enables RNA binding | nuclease activity | exonuclease activity |

| | | | | | |
|---|---|---|---|---|---|
| **Mrps30** | enables RNA binding | enables structur al constituent of ribosome | | | |
| **Endog** | enables single-stranded DNA endo-deoxyribonucle ase activity | enables magnesi um ion binding | enables nucleic acid binding | nuclease activity | enables endonucle ase activity |
| **Mrpl39** | enables nucleot ide binding | enables RNA binding | | | |
| **Tefm** | nucleic acid binding | enables RNA binding | enables protein binding | NOT enables crossover junction endodeoxy-ribonuclease activity | enables DNA polymerase processivity factor activity |
| **Mrpl10** | enables RNA binding | enables structur al constituent of ribosome | enables protein binding | | |
| **Slc30a9** | enables chroma tin binding | enables monoat omic cation transmembrane transporter activity | enables nuclear receptor binding | nuclear receptor coactivator activity | |
| **Mrps5** | enables RNA binding | enables structur al constituent of ribosome | | | |
| **Mterf2** | enables nucleic acid binding | enables DNA binding | enables DNA binding | enables protein binding | |
| **Supv3l1** | nucleotide binding | enables DNA binding | enables DNA helicase activity | enables RNA binding | enables RNA helicase activity |
| **Pds5b** | enables DNA binding | enables protein binding | | | |
| **Med31** | enables transcri ption coregulator activity | enables protein binding | enables ubiquiti n protein ligase activity | | |
| **Mapt** | enables DNA binding | enables minor groove of adenine-thymine-rich DNA binding | enables double-stranded DNA binding | enables single-stranded DNA binding | enables RNA binding |
| **Mbd2** | DNA binding | chromatin binding | enables satellite DNA binding | enables mRNA binding | enables protein binding |
